# PIEZO1 regulates leader cell formation and cellular coordination during collective keratinocyte migration

**DOI:** 10.1101/2022.10.13.512181

**Authors:** Jinghao Chen, Jesse R. Holt, Elizabeth L. Evans, John S. Lowengrub, Medha M. Pathak

**Author notes:** These authors contributed equally to this work (Jinghao Chen, Jesse R. Holt).

## Abstract

The collective migration of keratinocytes during wound healing requires both the generation and transmission of mechanical forces for individual cellular locomotion and the coordination of movement across cells. Leader cells along the wound edge transmit mechanical and biochemical cues to ensuing follower cells, ensuring their coordinated direction of migration across multiple cells. Despite the observed importance of mechanical cues in leader cell formation and in controlling coordinated directionality of cell migration, the underlying biophysical mechanisms remain elusive. The mechanically-activated ion channel PIEZO1 was recently identified to play an inhibitory role during the reepithelialization of wounds. Here, through an integrative experimental and mathematical modeling approach, we elucidate PIEZO1’s contributions to collective migration. Time-lapse microscopy reveals that PIEZO1 activity inhibits leader cell formation at the wound edge. To probe the relationship between PIEZO1 activity, leader cell formation and inhibition of reepithelialization, we developed an integrative 2D continuum model of wound closure that links observations at the single cell and collective cell migration scales. Through numerical simulations and subsequent experimental validation, we found that coordinated directionality plays a key role during wound closure and is inhibited by upregulated PIEZO1 activity. We propose that PIEZO1-mediated retraction suppresses leader cell formation which inhibits coordinated directionality between cells during collective migration.

**Author summary:** During the healing of a wound, cells called keratinocytes that make up the outer layer of the skin migrate collectively to close the wound gap. The mechanically activated ion channel PIEZO1 was previously found to inhibit wound closure. Here, through a combined modeling and experimental approach, we investigate the role of PIEZO1 in regulating collective migration. Specialized cells called leader cells, which typically form along the wound edge, are important for guiding the migration of neighboring cells. These leader cells dictate the coordinated directionality, or the cohesiveness of the migration direction between neighboring cells, through the transmission of mechanical and biochemical cues. We find that PIEZO1 activity inhibits the formation of these leader cells and, as a result, inhibits cell coordinated directionality causing the collective movement of cells to become disorganized and less effective in closing the wound. Our findings shed light on the complex mechanisms underlying collective migration, providing valuable insight into how mechanical cues affect the movement of cells during wound closure.

## Introduction

Cell migration plays an essential role in driving a diverse range of physiological processes including embryonic morphogenesis, tissue formation, repair and regeneration [1, 2]. This multistep process of cellular locomotion relies upon the coordination between several cellular processes including: actin polymerization, exertion of actomyosin-based contractile forces, and the dynamics of adhesion complexes [3]. During single cell migration, cells migrate directionally by becoming polarized. Located at the front of polarized cells, the leading edge drives forward locomotion while the rear, or retracting region, underlies the physical translocation of the cell body [4–6]. Under many physiological contexts, cells increase their migration efficiency by migrating together as a multicellular unit. During this collective form of cell migration, cells locomote while maintaining cell-cell contacts thus enabling subpopulations of cells to move interdependently [7, 8]. In addition to each cell polarizing individually, collectively migrating populations of cells become uniformly polarized due to the communication of mechanical and biochemical information through cell-cell contacts [9, 10]. This multicellular polarization is initiated by the highly specialized leader cells which are located at the front of groups of collectively migrating cells [11]. Leader cells are located at the tip of cellular outgrowths that develop along the wound edge and these cells are distinct from neighboring cells, as they display increased polarity and large lamellipodial protrusions [12]. Through the local coordination of intercellular mechanical forces, leader cells dictate the speed and the directional migration of individual follower cells located behind them [13–19]. Here, we use the term “coordinated directionality” to refer to how cohesively cells migrate in a direction similar to neighboring cells. This large-scale polarization and coordination of motion by leader cells is able to span across multiple cells, covering hundreds of micrometers in length [11, 20]. Thus the collective behaviors and dynamics of migrating sheets of cells are largely dependent upon the formation and dynamics of leader cells, and the transduction of guidance cues to the ensuing followers.

The collective movements of cells during epithelial sheet migration play a central role in guiding keratinocyte migration during reepithelialization, an essential component underlying the repair of wounded skin, wherein the cutaneous epidermal barrier is reinstated [21]. Recent work from our group identified the mechanically activated ion channel, PIEZO1, as a key regulator of the reepithelialization process [22]. Wounds generated in skin-specific *Piezo1* knockout mice (*Krt14*^*Cre*^*;Piezo1*^*fl/fl*^; hereafter *Piezo1* -cKO) were found to close faster than those in littermate Control (Control_cKO_) mice. On the other hand, *Krt14*^*Cre*^*;Piezo1*^*cx/+*^ and *Krt14*^*Cre*^*;Piezo1*^*cx/cx*^ mice (hereafter *Piezo1* -GoF) which express a skin-specific *Piezo1* gain-of-function mutation exhibited slower wound closure relative to littermate Control (Control_GoF_) mice (Fig 1A; [22]).

**Fig 1.**
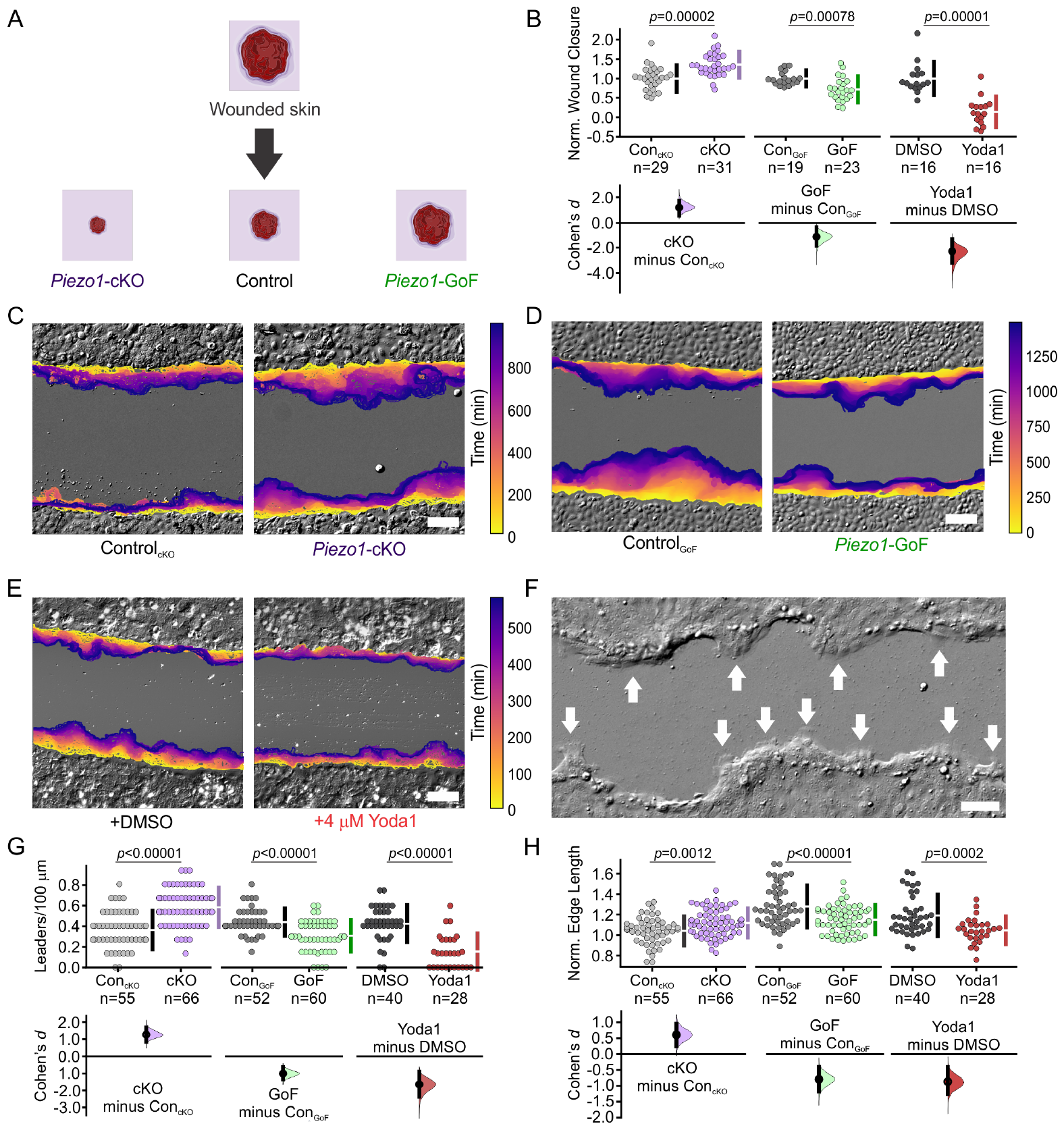
PIEZO1 activity inhibits wound edge dynamics and leader cell formation. **(A)** Summary schematic depicting PIEZO1’s effect on keratinocyte reepithelialization reported in Holt *et al*., 2021 [22]. **(B)** Reproduced from Fig 1L in [22] under a Creative Commons Attribution license, Cumming plot illustrating wound closure during *in vitro* scratch assays utilizing keratinocytes isolated from: Control_cKO_ and *Piezo1* -cKO mice (*left* ; *p* value calculated via two-sample t-test; Cohen’s *d* = 1.19; images from three independent experiments), Control_GoF_ and *Piezo1* -GoF mice (*middle*; *p* value calculated via two-sample t-test; Cohen’s *d* = -1.13; images from four independent experiments), and DMSO-treated and 4 *μ*M Yoda1-treated keratinocytes (*right* ; *p* value calculated via Mann-Whitney test; Cohen’s *d* = -2.28; images from three independent experiments). *n* in B denotes the number of unique fields of view imaged. **(C)** Representative overlay of the leading edge detected and segmented from DIC time-lapse images taken during *in vitro* scratch assay experiments in Control_cKO_ (left) and *Piezo1* -cKO (right) monolayers. Color of the cell boundary outline indicates passage of time. Scale bar = 100 *μ*m. The data in C are representative of three independent experiments. **(D)** Similar to C but for scratch assay experiments performed in Control_GoF_ (*left*) and *Piezo1* -GoF (*right*) monolayers. The data in D are representative of four independent experiments. **(E)** Similar to C but for scratch assay experiments performed in DMSO-treated (*left*) and 4 *μ*M Yoda1-treated (*right*) monolayers. The data in E are representative of three independent experiments. **(F)** Representative DIC image of wound closure during an *in vitro* scratch assay showing the appearance of finger-like protrusions led by leader cells (shown by white arrows). Scale bar = 100 *μ*m. See also S1 Fig. **(G)** Cumming plot showing the number of leader cells per 100 *μ*m which were manually identified from DIC time-lapse images along the wound margin in monolayers of: Control_cKO_ vs. *Piezo1* -cKO keratinocytes (*left* ; *p* value calculated via two-sided permutation t-test; Cohen’s *d* = 1.26), Control_GoF_ vs. *Piezo1* -GoF keratinocytes (*middle*; *p* value calculated via Mann Whitney test; Cohen’s *d* = -1), DMSO-treated vs. 4 *μ*M Yoda1-treated keratinocytes (*right* ; *p* value calculated via Mann Whitney test; Cohen’s *d* = -1.65). **(H)** Cumming plot showing quantification of the normalized edge length in monolayers of: Control_cKO_ vs. *Piezo1* -cKO keratinocytes (*left* ; *p* value calculated via two-sided permutation t-test; Cohen’s *d* = 0.6), Control_GoF_ vs. *Piezo1* -GoF keratinocytes (*middle*; *p* value calculated via two-sided permutation t-test; Cohen’s *d* = -0.8), DMSO-treated vs. 4 *μ*M Yoda1-treated keratinocytes (*right* ; *p* value calculated via two-sided permutation t-test; Cohen’s *d* = -0.9). To account for differences in the starting edge length which might occur when scratching monolayers in H, data are normalized by dividing the scratch length at either the end of the imaging period, or at the moment the wound edges touch, by the starting scratch length. A higher normalized edge length indicates a more featured wound edge, corresponding to the presence of more leader cells. *n* in G & H denotes the number of monolayer sheets imaged. See also Table 1.

Scratch wound assays performed in monolayers of keratinocytes isolated from these mice recapitulate the *in vivo* results, confirming that PIEZO1 activity inhibits keratinocyte reepithelialization (Fig 1B; [22]). Moreover, treatment of monolayers with Yoda1, a chemical agonist of PIEZO1, also resulted in delayed wound closure further indicating the channel’s involvement in regulating wound closure (Fig 1B) [23, 24]. Through a combined series of *in vitro* experimentation and bioimage analyses we determined that PIEZO1 channel activity increases localized cell retraction along the wound edge during *in vitro* wound closure assays, inhibiting advancement of cells and thus slowing wound closure. Our finding that PIEZO1 enhances retraction provided a working mechanism for how PIEZO1 activation slows wound closure, while the absence of the channel accelerates wound closure.

**Table 1.** Coordinated directionality is the key model parameter which replicates PIEZO1 reepithelialization phenotypes. *Top*: Summary table of monolayer experimental results on normalized wound closure and wound edge length. See also Fig 1. *Bottom*: Summary table of simulation results, depicting the effect of model parameters on normalized wound closure and wound edge length. For model parameters, single cell parameters (retraction strength, retraction duration, inter-retraction duration and cell motility) are separated from parameters which come from collective cell settings (cell-cell adhesion and coordinated directionality). A “+” indicates the wound feature is positively correlated with the model parameter, e.g., wound edge length increases with increased retraction strength, whereas “ *−* “ indicates a negative correlation, e.g., normalized wound closure is reduced with increasing retraction strength. Bolded italicized text denotes model parameters which correspond with experimental trends. See also Figs 2G, 2H and S3.

| Experimental Observations | Norm. Wound Closure | Norm. Edge Length |
| --- | --- | --- |
| *Piezo1*-cKO relative to Control<sub>cKO</sub> | + | + |
| *Piezo1*-GoF relative to Control<sub>GoF</sub> | − | − |
| Yoda1-treated relative to DMSO-treated | − | − |
| Model Parameter | Norm. Wound Closure | Norm. Edge Length |
| retraction strength | − | + |
| retraction duration | − | + |
| inter-retraction duration | + | − |
| cell motility | + | − |
| cell-cell adhesion | − | + |
| ***coordinated directionality*** | **+** | **+** |

Interestingly, several experimental studies have highlighted that mechanical cues play a role in leader cell formation [25, 26], guiding directional motion [27, 28], and governing the length scale of correlated motion during collective migration [14]. Given PIEZO1’s function as a key mechanotransducer and the channel’s contribution to keratinocyte reepithelialization, we asked whether PIEZO1 may affect the mechanoregulation of leader cells and cellular coordination during collective migration. Here, we take a combined theoretical and experimental approach to probe PIEZO1’s contribution to the biophysical mechanisms underlying keratinocyte collective migration.

Mathematical modeling has emerged as a powerful technique to systematically probe how biological factors contribute to the complex orchestration of collective migration [29–32]. Here, we build upon these previous works and develop a novel two-dimensional continuum model of reepithelialization. This model is derived by upscaling from a discrete model, incorporating key factors such as cell motility, retraction, cell-cell adhesion, and coordinated directionality. While motility and retraction are determined by single cell behaviors, cell-cell adhesion and coordinated directionality are influenced by the presence of neighboring cells. An upscaling procedure enables us to identify the contributions of these components to cell migration at the monolayer scale. We calibrated the cell-scale parameters in the model using data from experiments on single cells and performed parameter studies to investigate the influence of cell-cell adhesion and coordinated directionality, which are harder to measure experimentally. Our numerical simulations revealed that coordinated directionality is a critical factor in recapitulating the influence of PIEZO1 on wound closure and that elevated PIEZO1 activity leads to the inhibition of coordinated directionality. These predictions of the model were experimentally validated. Experiments also revealed that PIEZO1 activity suppresses the formation of leader cells, contributing further to the inhibition of collective migration during keratinocyte reepithelialization.

## Results

### PIEZO1 activity inhibits wound edge dynamics and leader cell formation

Efficient collective migration is driven by the formation of leader cells [33–35]. These highly specialized cells are distinct from their surrounding follower cells and play a key role in dictating collective dynamics [12]. In light of our previous finding that PIEZO1 activity inhibits wound closure [22], we sought to further characterize the effect of PIEZO1 on collective migration by investigating how PIEZO1 activity may affect leader cell formation. Since the emergence of leader cells drives collective migration and increased PIEZO1 activity results in delayed wound closure, we hypothesized that the number of leader cells would be affected by PIEZO1 activity levels. We generated scratch wounds in *Piezo1* -cKO, *Piezo1* -GoF, Yoda1-treated keratinocyte monolayers and their respective controls and utilized differential interference contrast (DIC) time-lapse imaging to examine the evolution of the wound margin (Fig 1C-E). During reepithelialization, multicellular finger-like protrusions often form along the wound margin as cells work together to close the wound area [36]. At the front of these cellular outgrowths, leader cells can be identified by their specialized phenotypic morphology in which they display a larger size, increased polarity, and prominent lamellipodia (Figs 1F and S1) [11, 35]. Leader cells were manually identified in time-lapse images of wound closure, similar to methods other groups have used for leader cell quantification within migrating collectives [37]. Given the different genetic backgrounds between conditions (i.e., *Piezo1* -cKO, *Piezo1-*GoF, Yoda1-treated) and the differences observed in migration properties across the Control samples of these different backgrounds [22], keratinocytes are only compared to control conditions of the same genetic background for all analyses. As hypothesized, we found that in *Piezo1* -cKO monolayers, the monolayer edge shows an increase in the number of leader cells compared to those from Control_cKO_ keratinocyte monolayers (Figs 1C, 1G and S1A). On the other hand in both *Piezo1* -GoF and Yoda1-treated monolayer conditions, where PIEZO1 activity is increased, the wound edge remains relatively flat throughout the imaging period due to a decrease in the formation of leader cells at the wound edge compared to respective control monolayers (Figs 1D, 1E, 1G, S1B and S1C). Furthermore, we noticed in *Piezo1-*cKO monolayers, that leader cells appeared to recruit more follower cells as seen by an increase in the width of fingering protrusions relative to Control_cKO_ monolayers (Figs 1C and S1).

To quantify the effect that PIEZO1 activity has on wound edge dynamics and leader cell protrusions, we also measured the change in the length of the wound edge within a field of view over the course of the imaging period, similar to methods employed by other groups [38]. The presence of leader cells, which are located at the front of cellular outgrowths, increases the length of the wound edge. Therefore, a shorter wound edge length would indicate fewer leader cells while a longer wound edge would indicate an increase in leader cells along the wound margin. We found that *Piezo1* -cKO monolayers have a longer wound edge length relative to Control_cKO_ monolayers, which further supports our observation that the absence of PIEZO1 results in increased leader cells along the wound edge (Fig 1H, *left*). Conversely, in both *Piezo1* -GoF and Yoda1-treated monolayers we found that edge lengths are significantly shorter than the respective control monolayers (Fig 1H, *middle, right*). Thus, we find that PIEZO1 inhibits the formation of leader cells, resulting in a shorter and flatter wound edge, while the absence of the channel results in a longer and more featured wound edge due to an increase in leader cell protrusions.

### Modeling PIEZO1’s influence on keratinocyte collective dynamics

Due to the many intricacies underlying the biological phenomena of collective cell migration, we adopted a theoretical approach as a framework for characterizing the biophysical relationship between PIEZO1 activity, leader cell initiation and wound closure. We designed a mathematical model of keratinocyte reepithelialization in order to study how PIEZO1 activity influences this process. We first separated reepithelialization into essential phenomenological components which could be incorporated into the design of the model as manipulable variables. As such we accounted for cell motility, and cellular retraction, a process central to the migration process and one which we previously found PIEZO1 activity promoted [22]. In our experimental data, we found that retraction varied in intensity such that in some instances it led to small regions of individual cells retracting while in other cases it led to the entire cell body pulling back away from the wound area [22]. Therefore we modeled retraction as a stochastic process at the leading edge associated with backward cell motion. We incorporated into our model design: (1) the average duration of retraction events at the monolayer edge, (2) the interval of time between sequential edge retractions, and (3) the strength of retraction. We also incorporated two hallmarks of collective cell migration: cell-cell adhesion and the coordination of keratinocyte migration direction, or coordinated directionality, both of which have been central to mathematical models proposed by other groups [32, 39–41]. Instead of modeling the mechanical forces involved in adhesion and retraction explicitly, we encoded the mechanistic effects such as cell motility, coordinated directionality, cell-cell adhesion, and retraction into model parameters. We then systematically manipulate these biological components of wound closure within our model and compare simulation results to experimental data garnered from scratch assays of PIEZO1 mutant keratinocytes (Table 1).

Due to the inherent multivariate nature of our system, we utilized a partial differential equation (PDE) model to spatiotemporally describe PIEZO1’s effect on reepithelialization. The PDE governing collective cell migration, which describes behavior at the monolayer scale, is derived by upscaling a discrete model at the single cell level. Simulations of wound closure, obtained by solving the nonlinear PDE numerically, enables a deeper understanding of how each model parameter contributes at both the single cell and monolayer levels. Furthermore, integration of experimental data at the single cell and monolayer scales allows for calibration of the model. We present a dimensionless version of the model here. We rescale the cell density by its maximal value, which can be quantified by counting the maximum number of cells in squares of a grid in the monolayer region away from the wound edge, where we expect cell density to exhibit spatial and temporal uniformity. The characteristic length scale *l* is defined as the distance from the wound edge to the region where the cells reach the maximal density in the monolayer (typically *∼* 10 cell lengths). Hence, our computational domain is a small region around the wound edge. From the experimental data, we can extract a characteristic wound edge velocity *v*, which allows us to derive a characteristic time *λ*^*−*1^ = *l/v*. See Section 6 in S1 Text for additional details.

The two-dimensional spatial discretization of a field of view containing a monolayer covered by a uniform grid of size *h* allows the labeling of indices (*i, j*) in space as ***x*** = ***x***_*i*,*j*_ = (*ih, jh*), and cell density, *ρ* = *ρ*(***x***, *t*) = *ρ*(*x, y, t*), a function of space, ***x*** = (*x, y*)^*T*^, and time, *t*, can be represented by *ρ*_*i*,*j*_ = *ρ*(***x***_*i*,*j*_, *t*) at time *t* (Fig 2A). The dimensionless experimental field of view is a unit square domain: [0, 1] *×* [0, 1] *∈* R^2^ (see Section 6 in S1 Text for the details of nondimensionalization). By incorporating the essential biological components of reepithelialization (Table 1, *bottom*), we construct the following discrete master equation (Fig 2A, *left* ; Eq. 1), which demonstrates the change rate of cell density over time (Eq. 1; left hand side) in response to the net flux of cells (Eq. 1; right hand side):

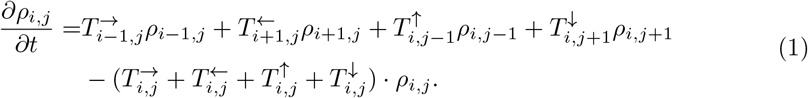

Here, the *T* ‘s are transitional probabilities per unit time associated with given directions of movement (i.e., 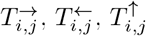 and 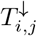) for cells migrating between adjacent grid points (e.g., from 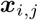 to 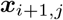 for 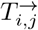). Each transitional probability accounts for cell motility, cell-cell adhesion, coordinated directionality, retraction events, and volume filling limitations.

**Fig 2.**
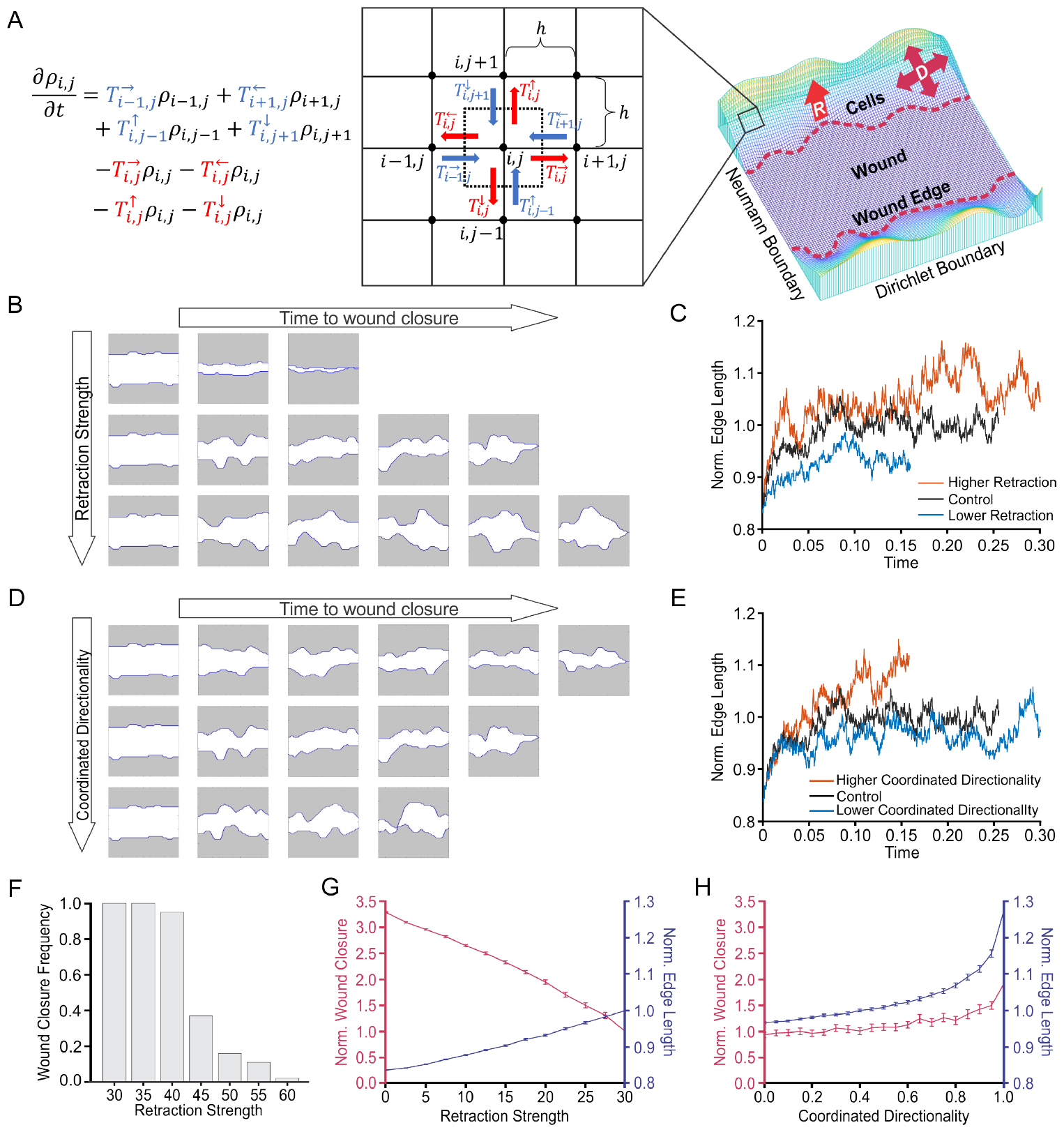
Coordinated directionality is the key model parameter which replicates PIEZO1 reepithelialization phenotypes. **(A)** Schematic showing a simplified visual of the modeling approach and visualization of the simulation domain. In the semi-discrete master equation (*left* ; Eq. 1), transitional probabilities associated with cell influx are highlighted in blue, while cell efflux related transitional probabilities are in red. Corresponding arrows depict this process on the grid (*middle*), indicating that the net flux is equal to the change in cell density over time at grid point (*i, j*). ***D*** represents coordinated directionality, and ***R*** represents retraction. **(B)** Simulation snapshots taken at equidistant time intervals depicting the evolution of the wound edge until wound closure (the moment interfaces touched) under low (*top*), Control (*middle*) and increased (*bottom*) levels of retraction strength. Shaded areas represent cell monolayers, while unshaded areas denote the cell-free space. **(C)** Plots showing quantification of the normalized edge length of simulated wounds, a measurement indicative of the number of leader cells, under different levels of retraction strength as a function of time. Shorter lines indicate simulation ending earlier due to faster wound closure. **(D, E)** Same as for (B) and (C), but under different levels of coordinated directionality. **(F)** The proportion of wound closure cases under different retraction magnitudes. The proportion of open wound closure cases start to decline after increasing retraction strength to 40, and almost no closure cases occur as retraction strength approaches 60. See also S2 Fig. **(G)** Line graphs showing the mean of 100 simulation results depicting the effect of retraction strength on normalized wound closure (*red; left axes*) and normalized edge length (*blue; right axes*). Error bars depict the standard error of mean. **(H)** Similar to (G) but for coordinated directionality. In C, E, F-H, all numbers have no unit because the model is dimensionless. See also S3 Fig and Table 1.

In the discrete master equation (Eq. 1), 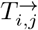 the transitional probability for cells traveling from ***x***_*i*,*j*_ to ***x***_*i*+1,*j*_ is defined as the following:

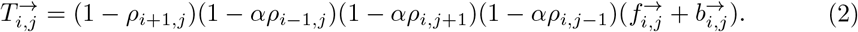

The term (1 *− ρ*_*i*+1,*j*_) models the effects of volume filling, e.g., if the cell density at ***x***_*i*+1,*j*_ has reached its maximal value, it restricts further cell movement into that point. The term (1 *− αρ*_*i−*1,*j*_)(1 *− αρ*_*i*,*j*+1_)(1 *− αρ*_*i*,*j−*1_) models cell-cell adhesion from three directions that hinder the cell migration, where *α ∈* [0, 1] is the adhesion coefficient, which is assumed to be the same in each direction [29]. In the last term 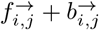, the vector 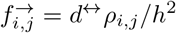 models diffusive cell motion while 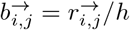 models cell movement due to retraction. The dependence on *h* reflects diffusive (*𝒪* (1*/h*^2^)) and advective (*𝒪* (1*/h*)) scaling of the equations, respectively. The diffusive component 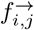, scaled as (𝒪 (1*/h*^2^), generates a diffusion flux that depends on the gradient of cell density. The advective component 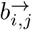, scaled as 𝒪 (1*/h*), results in an advection velocity independent of cell density that mimics the influence of retraction events (see Section 4 in S1 Text for details). Further, *d*^*↔*^ represents the magnitude of movement in the horizontal coordinate direction, while 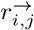 accounts for cell retraction. Note that the cell density term *ρ*_*i*,*j*_ in 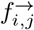 models the moving front that connects a region of zero cell density (wound) with a region of non-zero density (monolayer), e.g., [42]. The other transitional probabilities, 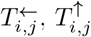 and 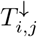 are defined analogously. Hence, Eq. 2 can be rewritten as

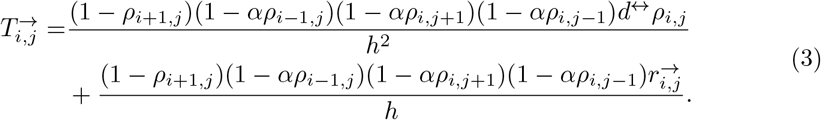

A continuum limit can be obtained by taking *h→* 0 in the discrete master equation (Eq. 1) to yield the partial differential equation

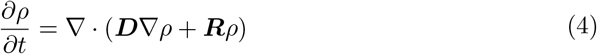

which is a diffusion-advection equation where the diffusion, ***D***, models cellular locomotion and coordinated directionality, whereas the advection velocity, ***R***, models retraction of the leading edge. The diffusion coefficient, or diffusivity, ***D***, is a 2 *×* 2 positive definite matrix given by

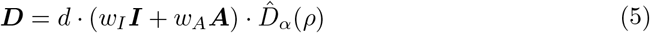

where *d >* 0 models cell motility during collective migration. The diffusion decomposition *w*_*I*_ ***I*** + *w*_*A*_***A*** combines the diffusion isotropy, where the identity matrix ***I*** = ***I***_2_ models the randomness of cellular migration, and diffusion anisotropy, where the matrix ***A*** models directed cellular migration. During wound closure, directional cues received from leader cells promote the migration of followers into the cell-free space to close the wound, thus promoting cells to have a higher probability of moving into the wound area and resulting in an anisotropic direction of diffusion.

The information regarding coordinated directionality is transmitted from the discrete level through the incorporation of distinct magnitudes of movement in the coordinate directions (*d*^*↔*^ and *d*^*↕*^ in Eq. 2, 3) that influence the transitional probabilities in two directions. Considering that cells receive signals to migrate towards the wound gap, we assume a larger magnitude of movement in the vertical direction (*d*^*↕*^ *≥ d*^*↔*^) based on our experimental configuration (Fig 2A, *right*). This assumption facilitates the following decomposition:

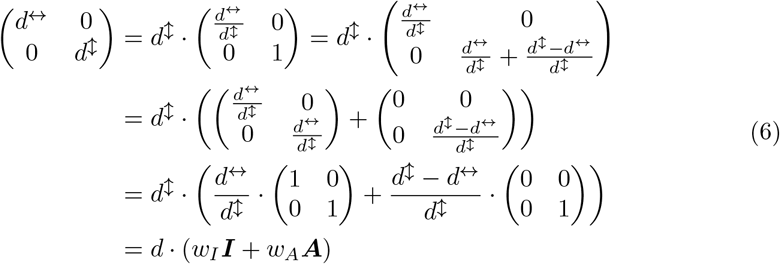

where the continuous coefficients of cell motility *d*, isotropic strength *w*_*I*_, and anisotropic strength *w*_*A*_ (representing coordinated directionality) are derived from the discrete coefficients of the magnitudes of movement in the coordinate directions *d*^*↔*^ and *d*^*↕*^ through the following relation:

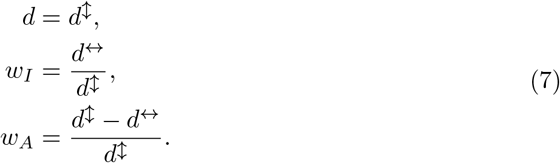

Here, the directionality assumption *d*^*↕*^ *≥ d*^*↔*^ guarantees the weights *w*_*I*_ and *w*_*A*_ are non-negative and bounded by 1, and the convex weighting relation *w*_*I*_ + *w*_*A*_ = 1 naturally holds from the derivation of *w*_*I*_ and *w*_*A*_. From these relations, we can observe how *w*_*A*_ measures coordinated directionality. Systems with stronger coordinated directionality, e.g., larger *w*_*A*_ that results from large relative differences between between *d*^*↕*^ and *d*^*↔*^, are more likely to migrate towards the direction of closure.

The scalar diffusion coefficient 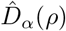 in Eq. 5 is a polynomial of cell density *ρ*:

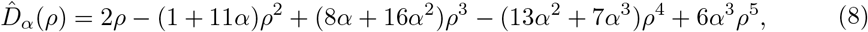

which is derived through a multi-scale modeling process from the scaled cell density *ρ*_*i*,*j*_*/h*^2^, cell-cell adhesion (e.g., (1 *− αρ*_*i−*1,*j*_)(1 *− αρ*_*i*,*j*+1_)(1 *− αρ*_*i*,*j−*1_) in 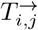) and volume filling (e.g., 1 − *ρ*_*i*+1,*j*_ in 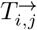). The adhesion coefficient, *α*, which lies in the range [0, 1], models the adhesion forces between adjacent cells, with a larger *α* corresponding to larger adhesion forces. Volume-filling limitations to cell movement are also modeled in 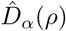 to hinder cells from migrating into a cell-dense area. In order to maintain a positive diffusivity, the value of *α* is bounded by *∼* 0.66 from above (see the detailed derivation in Section 3 in S1 Text).

Analogous to the derivation of diffusion, retraction, ***R*** (Fig 2A, *right*), is derived from the *𝒪* (1*/h*) component of the discrete transitional probability (Eq. 3) by taking the limit *h →* 0:

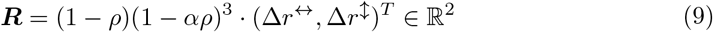

where 1 *− ρ* and (1 *− αρ*)^3^ model the effects of volume filling and cell-cell adhesion respectively. The retraction magnitude and directions are modeled phenomenologically in Δ*r*^*↔*^ and Δ*r*^*↕*^ as being localized in space and time, motivated by our previous work [22]. In particular, we assume:

1. Retraction occurs locally along the wound edge. This means only a part of wound edge cells are involved in retraction events at each time, while the rest of the cells on the edge and cells within the monolayer away from the edge just migrate by diffusion.
2. Retraction occurs intermittently in time. This means no retraction event is endless, i.e., no regions retract indefinitely. Hence at a wound edge point, there is a finite interval of duration time for each retraction event, and there is also a finite interval of time between two consecutive retraction events.

Because the computational domain is a small region around the wound edge (*∼* 10 cell lengths, see Section 6 in S1 Text), we assume there is one localized retraction region of fixed width that occurs at random times and locations on each side of the wound edge.

Following the localization assumptions (1) and (2), a choice for Δ*r*^*↔*^ and Δ*r*^*↕*^ is

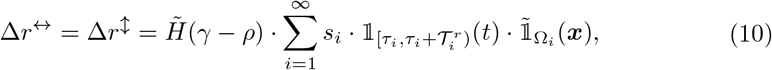

where *H* is a Heaviside function

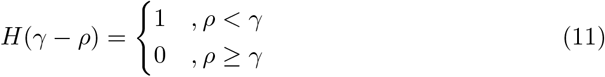

with threshold *γ*, which localizes the retraction to the wound edge (*γ* = 0.4 was adapted in the simulation). In particular, *H*(*γ − ρ*) = 0 turns off the retraction for *ρ > γ*, which is the high cell density region far away from the wound edge, while *H*(*γ − ρ*) = 1 turns on the retraction for *ρ < γ*, which is the low cell density region near the wound edge.

By labeling retraction events in chronological order with positive integers *i* = 1, 2, 3, …, indicator functions 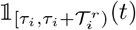 and 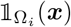 localize the regions where the edge retracts in time and space, respectively. We take the retraction to be localized in a region Ω_*i*_ := [*c*_*i*_ *− ω*_*r*_*/*2, *c*_*i*_ + *ω*_*r*_*/*2] *×* [0, 1] about a line segment *x* = *c*_*i*_ with width *ω*_*r*_ (*ω*_*r*_ = 0.2 was used in the simulation):

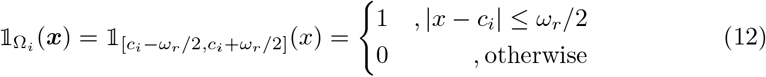

where we account for randomness by taking the uniform distribution *c*_*i*_ *∼ 𝒰* (0, 1). This allows the region Ω_*i*_ to randomly slide around [0, 1] to localize the retraction events.

The retractions are assumed to occur at particular times *τ*_*i*_ with durations 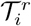.

Accordingly, we take

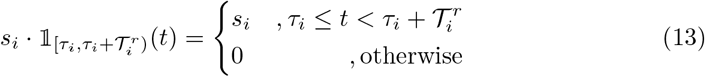

where *s*_*i*_ is the speed (or strength) of the retraction, and the next retraction occurs at 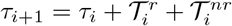 where 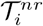 is the inter-retraction duration. To account for randomness, we assume:

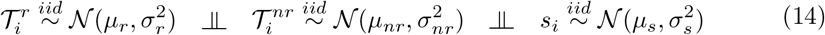

where 𝒩 (*μ, σ*^2^) denotes the normal distribution with mean *μ* and standard deviation *σ*, and all random variables are independent and identically distributed (*iid*). Thus, the mean strength of the retraction forces is *μ*_*s*_ and a single retraction event is sustained for a random duration with mean *μ*_*r*_. Any subsequent retraction will only start after a random idle duration with mean *μ*_*nr*_. The corresponding variances are 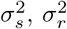, and 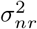, respectively. To ensure that our model incorporates only physically meaningful events, any negative duration or strength values that arise during the simulation are promptly discarded.

In summary, Δ*r*^*↔*^ and Δ*r*^*↕*^ are designed to model retractions such that cell movement would be governed by a diffusion-advection equation that guides the migrating cells in the retraction region near the wound edge:

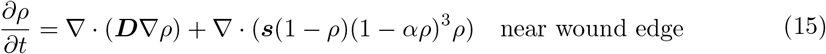

where ***D*** is the diffusivity (Eq. 5) and 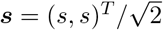 where the retraction strength, *s*, regulates the magnitude of advection velocity. On the other hand, cells far from the wound edge (e.g., interior of the monolayer) migrate following a simple diffusion equation

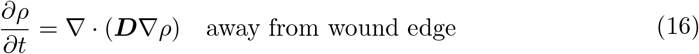

with the same diffusivity (Eq. 5). In fact, our model passes retraction information from the discrete to the continuous level. For example, if 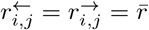 then there is no retraction at *x*_*i*,*j*_, e.g., monolayer region far away behind the wound edge. In this case, the governing equation is a pure diffusion equation without advection (Eq. 16) since 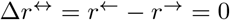, which appears in the continuum limit. In the retraction region, 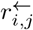 increases and 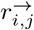 decreases, so Δ*r*^*↔*^ = *r*^*←*^ *− r*^*→*^*≠*0 and the governing equation turns into a diffusion-advection equation (Eq. 15).

Note that both Heaviside function *H*(*γ − ρ*) and characteristic functions 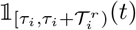 and 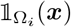 are discontinuous. To preserve differentiability, we smooth *H* using a hyperbolic tangent function 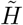 (Eq. 34 in Section 5 in S1 Text) and smooth 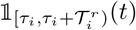 and 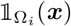 using generalized bell-shaped functions]. 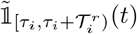 and 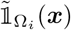 (Eq. 35 in Section 5 in S1 Text). In addition to the definition of Δ*r*^*↔*^ and Δ*r*^*↕*^ (Eq. 10) given above, there are alternative choices that can be adapted to interpolate the advection velocity between the retraction and non-retraction regions. However, the qualitative results of the model are not sensitive to the choice of Δ*r*^*↔*^ and Δ*r*^*↕*^ under assumptions (1) and (2) and the model reduces to Eq. 15 near the wound edge and to Eq. 16 far from the wound edge.

Since we only model a subset of the observation domain in the experiment, e.g., the region close to the wound edges as opposed to the whole experimental domain, we impose the following conditions at the boundaries of the computational domain:

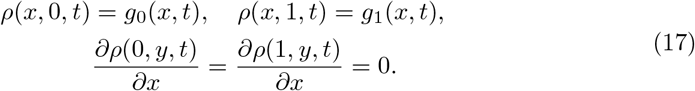

Horizontally on the top and bottom of the domain, time-dependent Dirichlet boundary conditions at ***x*** = (*x*, 0) and ***x*** = (*x*, 1) assign cell densities to the boundary points by functions *g*_0_ and *g*_1_, which mimic the effect of cells that flow into the observation domain area from the monolayer roughly perpendicular to the wound edge. The functions *g*_0_ and *g*_1_ are random functions of space and time (See Section 1 in S1 Text for the definitions of *g*_0_ and *g*_1_). Whilst vertically on the left and right sides of the domain, no-flux (Neumann) boundary conditions are used to approximate a net balance of cell influx and efflux into the observation domain roughly parallel to the wound edge, as suggested by the experiments.

The initial condition is generated by solving the PDE without retraction events for a short time period, which produces a banded heterogeneous monolayer with a cell-free region in the middle mimicking the initial wound (see Section 2 in S1 Text for details).

Summarizing, the model depends on the following biological parameters: (1) the mean retraction duration, *μ*_*r*_, (2) the mean inter-retraction duration, *μ*_*nr*_, (3) the mean retraction strength, *μ*_*s*_, (4) cell motility, *d*, in the absence of retraction (pure diffusion context), (5) cell-cell adhesion, *α*, and (6) the strength of coordinated directionality, *w*_*A*_. The governing equation (Eq. 4) is a nonlinear stochastic PDE, where stochasticity arises from the random coefficients. We solve the equations numerically using a finite difference method to obtain the cell density *ρ*(***x***, *t*) on the simulation domain over time until wound closure. Multiple simulations are performed under each condition to quantify the variability for the subsequent data analysis (See Methods Section *Numerical scheme* for details), from which we investigate how each model parameter influences collective migration during reepithelialization.

### Coordinated directionality is the key model parameter which replicates PIEZO1 reepithelialization phenotypes

Simulations of wound closure provide insight into how individual model parameters affect the wound closure process (Fig 2B-E). Thus, through a parameter study, we can explore the effects of model parameters on two experimentally-measured phenotypes affected by PIEZO1 activity during keratinocyte reepithelialization: (1) the rate of normalized wound closure and (2) normalized wound edge length, a measurement to characterize the degree of cellular protrusions and retractions during wound healing, which is correlated with leader cell presence (Fig 1G and 1H; [26]). These metrics were chosen because they can be directly measured and compared to experimental data.

During simulations we found that wounds would fail to close if parameters exceed a reasonable range (Figs 2F and S2). For instance, when retraction strength is set too high, cells are unable to overcome retractions of the wound edge which causes wounds to remain open indefinitely (Fig 2F). This model prediction is consistent with experimental results where Yoda1 treatment sometimes resulted in an increase in wound area during wound closure assays (Fig 1B; [22]).

By plotting the average rate of wound closure and edge length across multiple simulations we can see how the setting of individual model parameters compares to experimental trends we observe (Figs 2G, 2H and S3). We find that increasing the retraction strength parameter hinders wound closure, a result which is in line with the mechanism proposed in Holt *et al*., 2021 [22] (Fig 2G). However, our parameter study also shows that increased retraction strength results in a longer wound edge length, suggesting an increase in leader cell-like protrusions along the simulated wound margin. This contradicts our experimental observations in which a shorter wound edge length with fewer leader cells accompanies delayed wound closure (Fig 2G; Table 1). Similarly, we find that lower retraction strength elicited faster wound closure with shorter edge lengths due to fewer leader cell-like protrusions which also contradicts our experimental results (Fig 2G). Together, these results indicate that there is more to PIEZO1’s role in cell migration than retraction alone.

To identify possible contributors of wound closure regulation influenced by PIEZO1 activity, we performed an extensive parameter study in which we made adjustments to the model parameters of: cell-cell adhesion, retraction duration, inter-retraction duration, cell motility and coordinated directionality. We found that manipulation of all parameters aside from coordinated directionality fail to replicate the observed experimental results, i.e., faster wound closure accompanying a longer edge length, or conversely, delayed closure occurring with a shorter edge length (Table 1; S3 Fig). By increasing the coordinated directionality parameter within our model, wounds close faster with longer edge lengths due to the presence of more leader cell-like protrusions, replicating experimental observations in *Piezo1* -cKO monolayers (Fig 2H). On the other hand, under low coordinated directionality parameter conditions cells migrate more aimlessly, with formation of fewer leader-cell like protrusions along the wound edge and with inhibited closure, similar to observations from *Piezo1* -GoF and Yoda1-treated wounds (Fig 2H). Taken together, our parameter study predicts that while other model parameters, including retraction strength, affect keratinocyte migration, coordinated directionality plays a key role in modeling PIEZO1 inhibition of keratinocyte reepithelialization.

### PIEZO1 activity is predicted to regulate wound closure by hindering coordinated directionality

Through numerical simulation, our modeling parameter study reveals how altering individual model parameters one at a time while keeping the remaining parameters at their base values (S4 Fig) affects wound closure. However, experimental results reveal that PIEZO1 activity may alter more than one model parameter, which may generate compensating effects that reduce the contribution of coordinated directionality in the experimental setting. Therefore, we sought to further constrain the mathematical model by incorporating model parameters derived from experimental data. To this end, we utilized and expanded upon analyses performed on single migrating keratinocytes in our previous study [22], to compile an experimental dataset characterizing PIEZO1’s effect on: cell motility, retraction duration, inter-retraction duration and cell retraction strength (Figs 3A and S5; Table 2). Cell motility parameters were calculated by extracting cell speed information from single cell tracking experiments which were previously performed using single *Piezo1* -cKO and *Piezo1* -GoF keratinocytes [22] (Figs 3A, *left* and S5; Table 1). We expanded upon this work by also tracking individually migrating Yoda1-treated and DMSO-treated keratinocytes to incorporate the effect of Yoda1 on cell motility into our model predictions. Similar to our observations in *Piezo1* -GoF keratinocytes, Yoda1 treatment had no effect on the motility of single migrating keratinocytes compared to DMSO-treated control cells (S6 Fig). To find the average duration of retractions and intervals between successive retractions for all experimental conditions (*Piezo1* -cKO, *Piezo1* -GoF, Yoda1-treatment and the respective controls), we utilized two analysis methods performed in our previous study [22]: (1) kymographs (Fig 3A, *right*), which graphically depict the retraction and inter-retraction durations of the leading edge of migrating keratinocytes, and (2) a cell protrusion quantification software, ADAPT [43], which quantifies the strength of retraction events at the leading edge. Thus, from these measurements (Figs 3A and S5; Table 2), we can calibrate our model parameters based on experimental measurements, enabling us to make experimentally relevant predictions regarding PIEZO1’s influence on wound closure behavior.

**Table 2.** PIEZO1 activity affects single cell migration. Summary table presenting experimental results obtained from quantitative analysis of single cell migration experiments (e.g., kymograph, cell protrusion analyses, single cell tracking assays). A “+” indicates an increase, “ *−* “ indicates a decrease, and “ *∼* “ indicates no statistically significant change between Control and Test condition. All data aside from DMSO-treated and Yoda1-treated cell motility (S6 Fig) was initially published in [22]. Actual data values for each condition can be found listed in S5 Fig.

| Experimental Observations | Retraction Duration | Inter-retraction Duration | Retraction Strength | Cell Motility |
| --- | --- | --- | --- | --- |
| *Piezo1*-cKO relative to Control<sub>cKO</sub> | + | + | + | + |
| *Piezo1*-GoF relative to Control<sub>GoF</sub> | + | — | + | ~ |
| Yoda1-treated relative to DMSO-treated | — | — | + | ~ |

**Fig 3.**
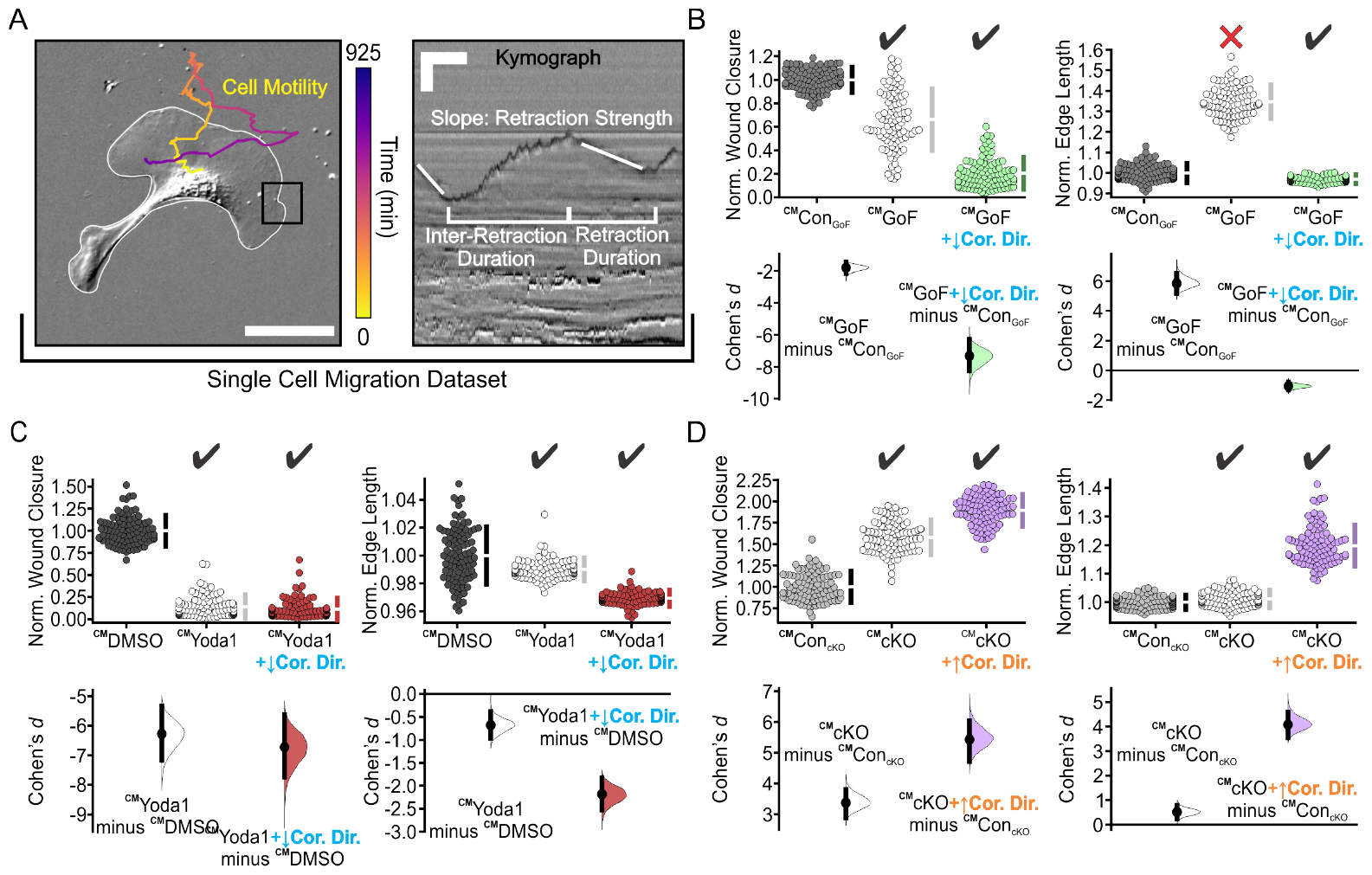
PIEZO1 activity is predicted to regulate wound closure by hindering coordinated directionality. **(A)** Schematic depicting experimentally measured features used to generate the single cell migration dataset. Left, representative still image of migrating keratinocyte with overlaid cell trajectory. Trajectory is derived from tracking cell motility during time-lapse experiments. Color denotes passage of time such that yellow is the starting position and purple denotes track end position. Cell boundary is in white. Scale bar = 100 *μ*m. Kymographs (*right*) taken at the leading edge of migrating cells (e.g., similar to black box in the left image) are used to obtain information regarding inter-retraction duration and retraction duration. The cell protrusion quantification software, ADAPT [43] was used to gain information regarding retraction strength. Scale bar = 10 *μ*m, Time bar = 5 min. **(B)** Cumming plots showing simulation results using the calibrated model (^CM^) to predict how PIEZO1 affects normalized wound closure (*left plots*) and wound edge length (*right plots*) in simulated Control_GoF_ monolayers (*dark gray*), *Piezo1* -GoF monolayers without altered coordinated directionality parameters (*white*), and *Piezo1* -GoF monolayers with coordinated directionality decreased (*green*). See Methods Section *Model parameter adjustment* for the details. **(C)** Similar to B but using simulation results from DMSO-treated monolayers (*black*), Yoda1-treated monolayers without altered coordinated directionality parameters (*white*), and Yoda1-treated monolayers with coordinated directionality decreased (*red)*. **(D)** Similar to B but using simulation results from Control_cKO_ monolayers (*light gray*), *Piezo1* -cKO monolayers without altered coordinated directionality parameters (*white*), and *Piezo1* -cKO monolayers with coordinated directionality increased (*purple)*. In B-D, n = 100 simulation results for each condition, and ^CM^ denotes “Calibrated Model”. To account for differences between control cases, data are normalized by rescaling to the mean of the corresponding control. Larger normalized wound closure indicates faster wound closure, while a smaller normalized wound closure indicates slower wound closure. Similarly, a larger normalized edge length indicates a more featured wound while a smaller normalized edge length indicates a flatter or less featured wound. Black check marks at the top of each plot condition indicate that simulation results match experimental trends while a red cross indicates simulation fails to match the experiment trends. See also Table 3. For comparison with experimental data see Fig 1B, 1G and 1H.

To calibrate our model, we created a respective simulation control for each experimental condition (*Piezo1* -cKO, *Piezo1* -GoF and Yoda1-treated) by fixing the values of model parameters to a basecase, where the frequency of retraction was set to be the same as the corresponding experimental control. For a given experimental condition, the model parameters related to retraction (retraction duration, inter-retraction duration, retraction strength) and cell motility were adjusted from the control condition by the same proportions as their experimentally-measured changes relative to the control condition (Figs 3A and S5; Table 2). In particular, the mean retraction and inter-retraction durations *μ*_*r*_ and *μ*_*nr*_, the cell motility *d* and the mean retraction strength *μ*_*s*_ are changed proportionally in the model (see Methods Section *Model parameter adjustment* for details). With cell-cell adhesion and coordinated directionality unchanged compared to Control_GoF_, we find that while we can replicate simulated monolayers of *Piezo1* -GoF keratinocytes having slower wound closure compared to simulated Control_GoF_ monolayers, we fail to observe the expected decreasing change in leader cell-like protrusions as indicated by a smaller simulated monolayer edge length (Fig 3B). However, by lowering the collective migration parameter of coordinated directionality, we recapitulate the experimental phenotype of both a shorter edge length and slower wound closure in simulated *Piezo1* -GoF monolayers (Fig 3B). On the other hand, we see that model simulations calibrated by the single cell migration dataset for both *Piezo1* -cKO and Yoda1-treated keratinocytes reproduce the expected experimental trends; however, by incorporating changes to coordinated directionality we observe a stronger effect (Fig 3C and 3D). Notably, we observe that adjustment of cell-cell adhesion parameters, another model parameter integral to collective migration, fails to replicate all experimental results, reinforcing that coordinated directionality plays a primary role in PIEZO1’s effect on reepithelialization (Table 3; S7 Fig). The more retraction regions generated, the slower the wound healing process and the longer the wound edge length. Matching the experimental results when PIEZO1 signaling is upregulated still required a decrease in coordinated directionality. Taken together, these studies demonstrate that only by including alterations to coordinated directionality are we able to mimic all experimental phenotypes.

**Table 3.**
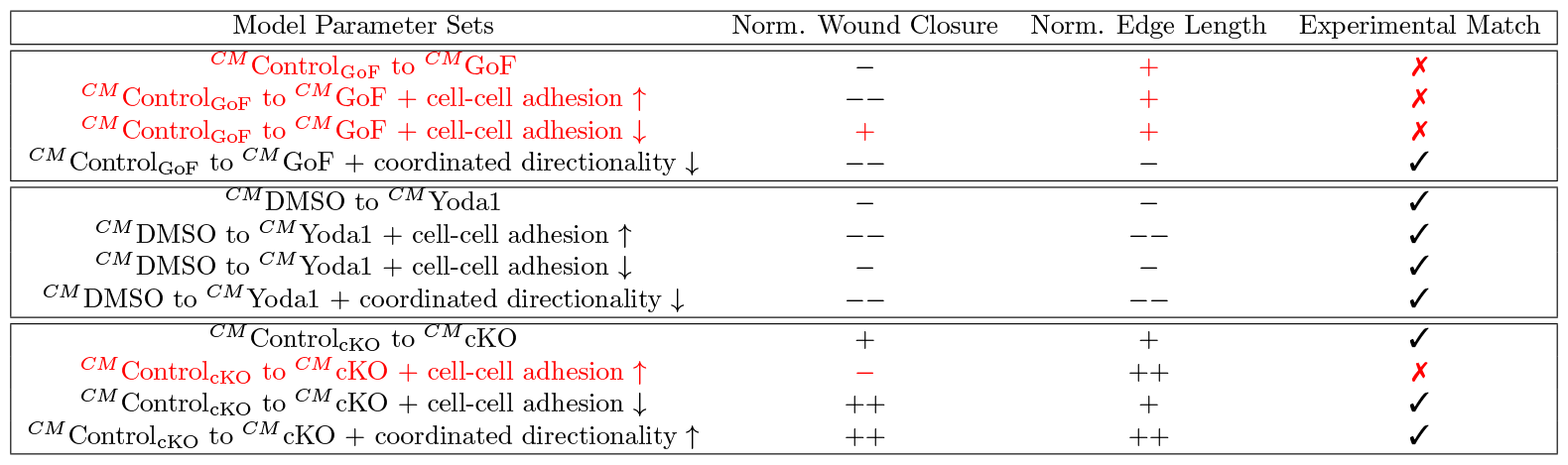
Coordinated directionality recovers monolayer closure behavior from single cell data. Summary table depicting simulation results using the calibrated model (^CM^) to predict how PIEZO1 affects normalized wound closure and normalized edge length with altered adhesion and coordinated directionality parameters. A “+” indicates a parameter set has a predicted increase upon an experimental measure while a “*−* “ indicates a predicted decrease. Double signs (++/*−−*) represent a stronger observed effect on the simulated measure than single signs (+/*−*). Red font and cross mark ✗ indicate that model predictions calibrated by the “Single Cell Migration” dataset do not match experimental trends (Table 1), while a check mark **✓**indicates that model predictions are consistent with experimental results. See also Figs 3, S7 and S8.

To test whether the effect of higher PIEZO1 activity hindering coordinated directionality is sensitive to the details of the mathematical model of cell-cell adhesion, we also considered a phenomenological continuum modeling framework (see Section 7 in S1 Text for details) in which the diffusion coefficient was assumed to be a decreasing function of cell-cell adhesion instead of upscaling from a discrete model. This alternative approach follows that of Amereh *et al*., 2021 [44] and does not rely on upscaling from a discrete model. We repeated all the simulations using this new model of adhesion and found that the results are qualitatively consistent with the upscaled adhesion model we originally considered (S14 and S15 Figs), supporting the idea that the hindering effect of cell-cell adhesion can be modeled in various ways without altering the qualitative results.

We also tested the sensitivity of our conclusion regarding the role of PIEZO1 on coordinated directionality with respect to the details of the mathematical model of cell motilities and retraction processes. We re-calibrated the model using cell motilities from monolayer experiments (S9 Fig) rather than single cell experiments (S5 Fig). Consistent with the single cell data, cell motility within the monolayer increased in *Piezo1* -cKO compared to Control_cKO_. In *Piezo1* -GoF and Yoda1-treated conditions, the cell motilities decreased in the monolayer (compared to respective controls), which is different from the behavior of single cells where the motilities were the same. Nevertheless, the re-calibrated model still predicts that PIEZO1 activity decreases coordinated directionality (see Section 8 in S1 Text for details).

We also varied the magnitudes of the retraction processes (retraction duration, inter-retraction duration and retraction strength). Because our previous experimental work [22] indicated qualitative consistency between single-cell and monolayer retraction processes but did not measure these features quantitatively in monolayers, we tested three different magnitudes of these processes in the model (S18 Fig), rather than using magnitudes drawn directly from the single cell experiments as previously done. The results still predict that PIEZO1 hinders coordinated directionality (S18 Fig, also see Section 8 in S1 Text for details).

Thus, our model predicts that PIEZO1 activity affects coordinated directionality within monolayers such that increased PIEZO1 activity inhibits the cells ability to move cohesively during collective migration, ultimately delaying wound closure. On the other hand, in monolayers which lack PIEZO1 expression, cells are predicted to have stronger directionality signals and recruit more follower cells to close the wound faster.

### PIEZO1 activity inhibits persistence of direction during keratinocyte collective migration

To test our model’s prediction we first utilized a cell tracking assay to examine the motility of individual cells during collective migration. To track the movement of individual cells within monolayers we utilized the live-cell DNA stain SiR-Hoechst to label individual nuclei within monolayers [45]. After imaging collective cell migration over the course of several hours, we tracked the movement of individual nuclei and analyzed the resulting cell trajectories (Fig 4A-C). The mean squared displacement (MSD) is a common metric for analyzing cell displacement as a function of time. Replicating our single cell migration observations [22], we observe that individual tracked nuclei within *Piezo1-*cKO monolayers have MSDs that are greater than that of Control_cKO_ cells, demonstrating a larger area explored (Fig 4D). Measurement of the instantaneous cellular speed reveals that, similar to our previous observations, *Piezo1* -cKO cells migrate faster relative to littermate Control_cKO_ cells (S9A Fig). On the other hand, cells from both *Piezo1* -GoF and Yoda1-treated monolayers have MSDs that are significantly smaller (Fig 4E and 4F). This effect is distinct from our observation in single migrating cells, where we observed that *Piezo1* -GoF keratinocytes migrate farther than Control_GoF_ cells [22], and that Yoda1-treatment has no difference relative to DMSO-treated control cells (S6A Fig). Moreover, in both *Piezo1* -GoF and Yoda1-treated monolayers we observe that increased PIEZO1 activity inhibits migration speed (S9B and S9C Fig). This observation also differs from our single cell migration observations in which both *Piezo1* -GoF and Yoda1-treated keratinocytes have no difference in migration speed compared to respective control cells. Our observed differences for PIEZO1’s effect on speed and MSD between single cell and collective migration results may be attributed to additional mechanical information from cell-cell interactions during collective migration affecting activation of PIEZO1.

**Fig 4.**
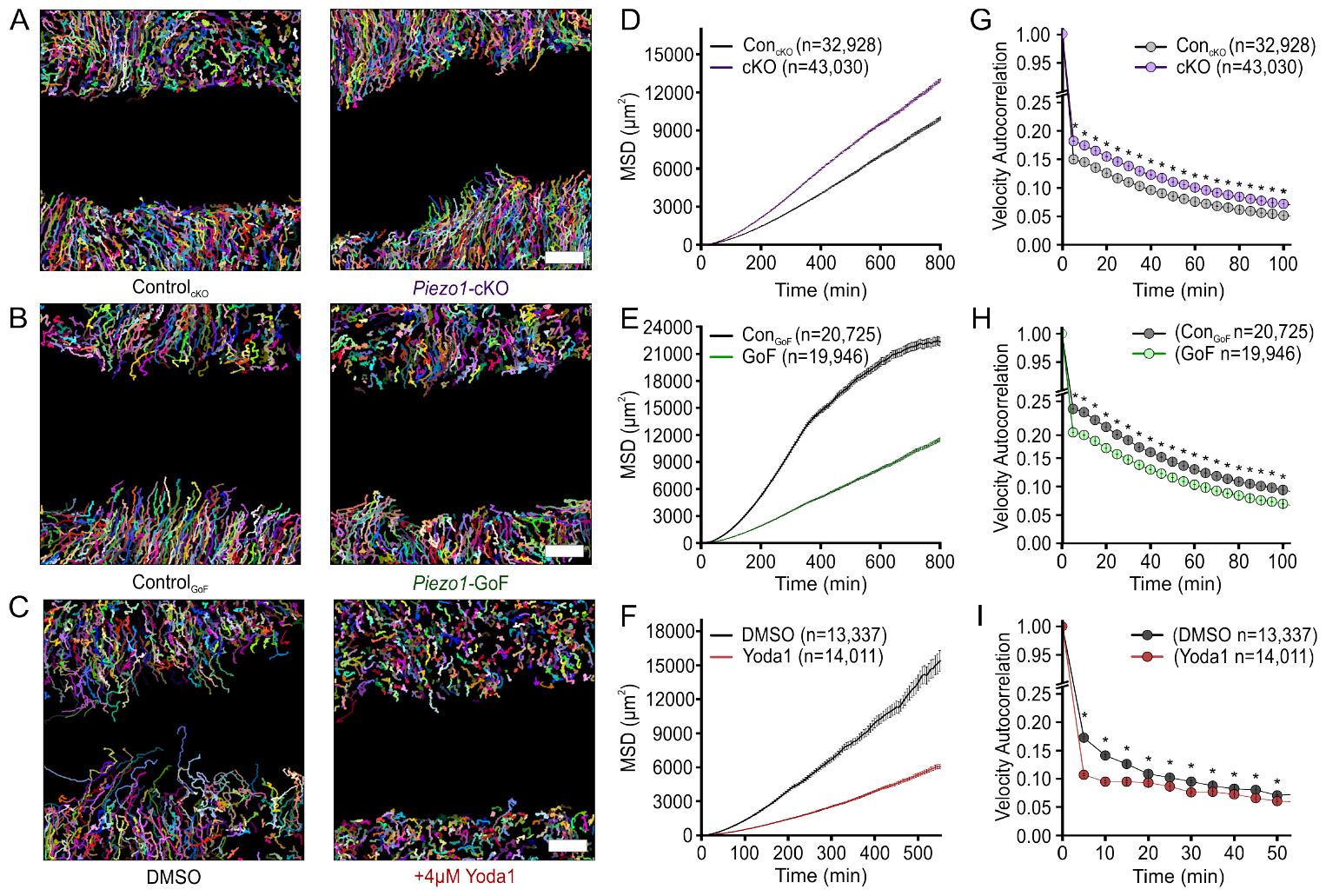
PIEZO1 activity inhibits persistence of direction during keratinocyte collective migration. **(A-C)** Representative field of view depicting individual cell trajectories derived from tracking **(A)** Control_cKO_ (*left*) and *Piezo1* -cKO (*right*) keratinocytes, **(B)** Control_GoF_ (*left*) and *Piezo1* -GoF (*right*) keratinocytes, and **(C)** DMSO-treated (*left*) and 4 *μ*M Yoda1-treated (*right*) keratinocytes during collective migration experiments. Trajectory color depicts individual cell trajectories. Scale bar=100 *μ*m. **(D-F)** Average mean squared displacement (MSD) plotted as a function of time for: **(D)** Control_cKO_ (*gray*) and *Piezo1* -cKO (*purple*) keratinocytes, **(E)** Control_GoF_ (*gray*) and *Piezo1* -GoF (*green*) keratinocytes, and **(F)** DMSO-treated (*gray*) and 4 *μ*M Yoda1-treated (*red*) keratinocytes. All error bars plotted as SEM, in some instances error bars are smaller than symbols. **(G-I)** Average velocity autocorrelation measurement of: **(G)** Control_cKO_ (*gray*) and *Piezo1* -cKO (*purple*) keratinocytes, **(H)** Control_GoF_ (*gray*) and *Piezo1* -GoF (*green*) keratinocytes, and **(I)** DMSO-treated (*gray*) and 4 *μ*M Yoda1-treated keratinocytes, plotted as a function of time (* denotes *p* value*<*0.0001 as calculated via Kolmogorov–Smirnov test). For Control_cKO_ (n=66 unique fields of view) and *Piezo1* -cKO (n=85 unique fields of view) data plotted in A, D, G, images taken from three independent experiments. For Control_GoF_ (n=56 unique fields of view) and *Piezo1* -GoF (n=51 unique fields of view) data plotted in B, E, H, images taken from four independent experiments. For DMSO-treated (n=32 unique fields of view) and 4 *μ*M Yoda1-treated (n=31 unique fields of view) keratinocyte data plotted in C, F, I, images taken from three independent experiments. Plotted n denotes the number of individual cell trajectories. See also S9 Fig.

Since coordinated directionality can, in part, be inferred by how straight the trajectories of cells in a collectively migrating group are, we measured the directional persistence of individual cell trajectories. While coordinated directionality refers to how cohesively cells migrate in a similar direction, directional persistence refers to the directed migration of individual cells or, more simply, how straight individual cell trajectories are. Notably, these two elements are often seen to co-occur in monolayers which show increased wound closure efficiency [46]. The directional persistence of a cell can be quantified by measuring the velocity autocorrelation of cell trajectories [47]. The randomness in direction of a cell’s trajectory is indicated by how rapidly its velocity autocorrelation function decays: autocorrelation curves which decay slower indicate cells that have straighter migration trajectories. Measurement of the velocity autocorrelation shows that *Piezo1* -cKO keratinocytes migrating in cell monolayers move straighter than Control_cKO_ cells (Fig 4G), similar to our previous findings in single migrating cells. In both *Piezo1* -GoF and Yoda1-treated keratinocytes, cells move less straight than their respective controls (Fig 4H and 4I). This finding also differs from findings in single cell migration results wherein Yoda1-treatment does not change directional persistence (S6C Fig) while the *Piezo1* -GoF mutation induces straighter trajectories during single cell migration [22].

Taken together, our results show that PIEZO1 activity inversely correlates with both cell speed and the persistence of migration direction during keratinocyte collective migration. Our observation that the directional persistence of individual keratinocytes within a monolayer is inhibited by PIEZO1 activity during collective cell migration provides initial support for our model’s prediction that coordinated directionality is affected by PIEZO1 activity.

### Increased PIEZO1 activity inhibits the coordination of cellular motion

The coordinated movement of keratinocytes during wound reepithelialization depends on the large-scale interactions of multiple cells as they work together to close wounds. While tracking individual cells in a monolayer provides useful information regarding the locomotion of individual cells, it does not fully describe the dynamics of collectively migrating cells. To further validate our model’s prediction that PIEZO1 activity inhibits coordinated directionality and to characterize the effect of PIEZO1 on large scale cellular interactions during wound closure we utilized particle image velocimetry (PIV). PIV is an optical method of flow visualization which allows us to dynamically map the velocity fields of migrating keratinocytes within a monolayer during wound closure [20, 48, 49]. By isolating the individual velocity vectors comprising a monolayer’s vector field and mapping the frequency of vector directions for samples from different conditions (e.g., *Piezo1-*cKO, *Piezo1-*GoF, and Yoda1-treatment), we can visualize how PIEZO1 affects the coordinated directionality and overall coordination of motion between cells during wound closure (Fig 5A-C). Probability density distributions of velocity directions from Fig 5A-C illustrate that *Piezo1* -cKO cells flow towards the wound margin (denoted by 0 degrees) to a greater extent than littermate Control_cKO_ cells (Fig 5D). Conversely, *Piezo1-*GoF and Yoda1-treated cells flow towards the wound margin to a lesser extent than their corresponding Controls (Fig 5E and 5F).

**Fig 5.**
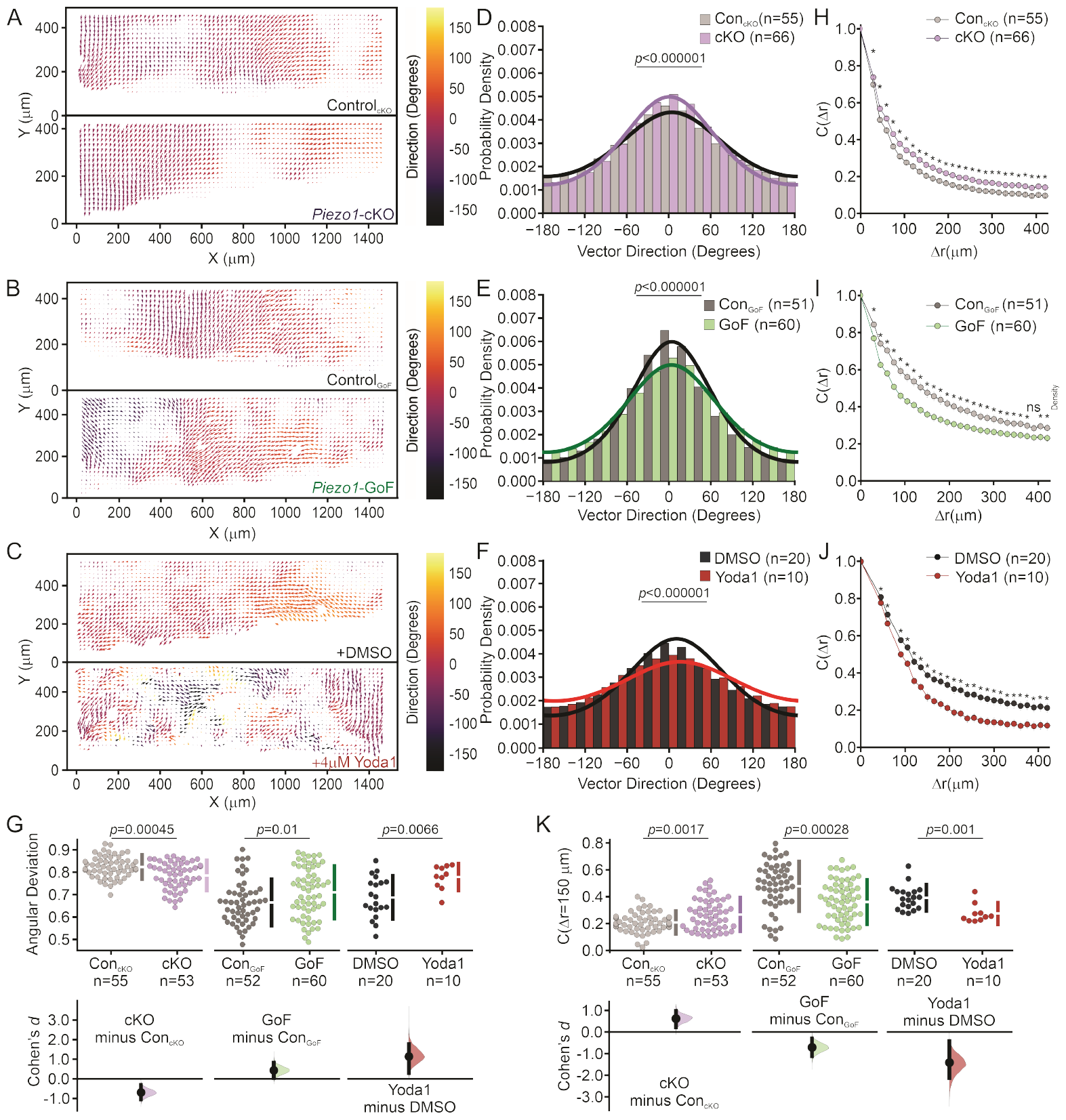
Increased PIEZO1 activity inhibits coordinated cellular motion. **(A-C)** Representative mean Particle Image Velocimetry (PIV) flow fields derived from time-lapse images of labeled nuclei from collectively migrating monolayers of: **(A)** Control_cKO_ (*Top*) and *Piezo1* -cKO (*Bottom*) keratinocytes, **(B)** Control_GoF_ (*Top*) and *Piezo1* -GoF (*Bottom*) keratinocytes, and **(C)** DMSO-treated (*top*) and 4 *μ*M Yoda1-treated keratinocytes (*Bottom*) during time-lapse scratch assay experiments. An individual flow field comprises either the upper or lower monolayer sheet of a scratch assay. Flow fields are oriented such that for the Y-direction, 0 *μ*m is positioned within the cell free region. **(D-F)** Distribution plots showing the probability density of velocity vector direction for: **(D)** Control_cKO_ (*gray* ; *κ* = 0.51) and *Piezo1* -cKO (*purple*; *κ* = 0.71) monolayers, **(E)** Control_GoF_ (*gray* ; *κ* = 1.01) and *Piezo1* -GoF (*green*; *κ* = 0.71) monolayers, and **(F)** DMSO-treated (*gray* ; *κ* = 0.61) and Yoda1-treated (*red* ; *κ* = 0.30) monolayers. The curves depicted in the figure represent the fitting of von Mises distributions, where a smaller reported *κ* corresponds to less directed migration while a larger *κ* indicates an increase in directed migration. For D-F, *p* value calculated by Chi-squared test. **(G)** Cummings plot showing the mean angular deviation, or the variability in velocity direction isolated from PIV flow fields in: Control_cKO_ vs. *Piezo1* -cKO monolayers (*left* ; *p* value calculated via two-sample t-test; Cohen’s *d* = -0.7), Control_GoF_ vs. *Piezo1* -GoF monolayers (*middle*; *p* value calculated via two-sample t-test; Cohen’s *d* = 0.43) or DMSO-treated vs. 4 *μ*M Yoda1-treated monolayers (*right* ; *p* value calculated via two-sample t-test; Cohen’s *d* = 1.14). Data are normalized such that 1 indicates highly random velocity directions and 0 indicates highly uniform velocity directions. **(H-J)** Spatial autocorrelation, *C*(Δ*r*), of the radial velocity component, which is a measure of the spatial coordination of neighboring cells in monolayers, plotted as a function of increasing length scales of: **(H)** Control_cKO_ (*gray*) and *Piezo1* -cKO (*purple*) keratinocytes, **(I)** Control_GoF_ (*gray*) and *Piezo1* -GoF (*green*) keratinocytes, and **(J)** DMSO-treated (*gray*) and Yoda1-treated (*red*) keratinocytes. For H, I, J * denotes a statistically significant difference, and ns denotes not statistically significant as determined by one way ANOVA test. Specific *p* values for plotted points can be found in S11 Fig. See also S10 Fig. (**K**) Local spatial coordination, *C*(Δ*r* = 150 *μm*), of keratinocytes where the correlation value is set at 150 *μ*m to measure the coordination of motion with neighboring cells in: Control_cKO_ vs. *Piezo1* -cKO monolayers (*left* ; *p* value calculated via two-sample t-test; Cohen’s *d* = 0.62), Control_GoF_ vs. *Piezo1* -GoF monolayers (*middle*; *p* value calculated via two-sample t-test; Cohen’s *d* = -0.7) or DMSO-treated vs. 4 *μ*M Yoda1-treated monolayers (*right* ; *p* value calculated via Mann-Whitney test; Cohen’s *d* = -1.4). *n* in B, C, E, F, H, I, J and K denotes the number of monolayer sheets imaged. For Control_cKO_ and *Piezo1* -cKO data plotted in A, D, G (*left*), H, and K (*left*), images are taken from three independent experiments. For Control_GoF_ and *Piezo1* -GoF data plotted in B, E, H (*middle*), I, and K (*middle*), images are taken from four independent experiments. For DMSO-treated and 4 *μ*M Yoda1-treated keratinocyte data plotted in C, F, H (*right*), J, and K (*right*), images are taken from two independent experiments.

To fit our vector direction datasets, we employed the von Mises distribution by minimizing the mean squared error with the von Mises probability density function (Eq. 21 in the Methods Section). The resulting fitted curves (Fig 5D-F) provide the best approximation of the data by adjusting the distribution parameters, including the mean (*μ*) and the concentration (*κ*, indicating the strength of directed migration in our experimental context). A smaller *κ* value corresponds to a flatter bell curve and a distribution closer to uniform, indicating less directed migration. Conversely, a larger *κ* value results in a sharper bump in the probability density function, indicating an increase in directed migration. We find that *Piezo1* -cKO cells show a higher *κ* (*κ* = 0.71) than Control_cKO_ (*κ* = 0.51) indicating that *Piezo1* -cKO cells move with increased coordination relative to Control_cKO_ cells (Fig 5D). On the other hand, we find that both *Piezo1* -GoF and Yoda1-treated monolayers show a loss in directed migration as illustrated by the broader distribution of isolated vector directions and a lower calculated *κ* value for experimental conditions (*κ* = 0.71 for *Piezo1* -GoF, *κ* = 0.30 for Yoda1-treated) relative to the respective control populations (*κ* = 1.01 for Control_GoF_, *κ* = 0.61 for DMSO-treated; Fig 5E and 5F).

PIEZO1’s effect on coordinated directionality can be further parameterized by measuring the angular deviation, or the variability in velocity direction for all vectors within a PIV vector field. Thus, the range of the angular deviation indicates how coordinated the direction of cellular motion is within an entire monolayer field of view such that a higher angular deviation indicates less coordination. We observe that *Piezo1* -cKO monolayers have a lower average angular deviation value relative to Control_cKO_ monolayers, indicating a smaller spread in velocity direction (Fig 5G, *left*). This is opposed to *Piezo1* -GoF and Yoda1-treated monolayers which both show a higher angular deviation than the respective controls, further signifying that PIEZO1 activity promotes less directional migration (Fig 5G, *middle, right*). We note that any difference in the angular deviation between control conditions can likely be attributed to different genetic backgrounds between control conditions.

Recognizing that the synchronized movement of groups of cells during collective migration relies upon the coordination of migration direction across individual cells, we next looked at how PIEZO1 activity affects the distance over which cells align, or correlate, their motion within a monolayer. To do this, we determine how alike the velocity of nearby cells is by calculating the average spatial autocorrelation of velocity vectors, (*C*(Δ*r*)), which measures the degree of correlation between velocity vectors of cells at increasing length scales within a monolayer (Fig 5H-K). If keratinocytes within a monolayer are migrating together with high directional uniformity we expect a higher autocorrelation value, while a lower autocorrelation value indicates that individual keratinocytes are moving more independently of one another. Therefore, the decay rate of the average spatial autocorrelation curve indicates how coordinated a given cell’s direction of motion is to that of another cell located at iteratively increasing distances away (Fig 5H-J). Measurement of the spatial autocorrelation in *Piezo1* -cKO and Control_cKO_ monolayers illustrate that *Piezo1* -cKO cells show an increase in coordination with cells at greater distances relative to Control_cKO_ cells, as indicated by a slower decay of the average *Piezo1* -cKO autocorrelation curve (Fig 5H). The length constant, or distance at which the spatial autocorrelation reaches a value of 0.37, was estimated by fitting an exponential curve to our experimental dataset. Calculations of the length constant for *Piezo1* -cKO cells show an increase in coordination by 21.47 *μ*m farther than Control_cKO_ (Figs 5H and S10). To further quantify the coordination between nearby cells we measure the spatial autocorrelation values at 150 *μ*m, the distance of a few cell-lengths away. Measurement of local autocorrelation values in *Piezo1-*cKO keratinocytes cells show an increased level of coordination of locomotion with neighboring cells compared to cells in Control_cKO_ monolayers (Fig 5H and 5K). In contrast, both *Piezo1* -GoF and Yoda1-treated monolayers exhibit less coordinated movement with neighboring cells when compared to control cells (Fig 5I-K). Length constants in Yoda1-treated and *Piezo1* -GoF cells show a 58.560 *μ*m and 85.54 *μ*m decrease, respectively, in their coordination of motion relative to the respective control monolayers (Figs 5I, 5J and S10). Therefore, we find that PIEZO1 activity disrupts the distance over which cells coordinate their motion during wound closure which inhibits the efficiency of collective migration.

Here, we have uncovered the inhibitory impact of PIEZO1 activity on coordinated cell directionality within a monolayer. This discovery naturally raises additional questions about the consequences of heterogeneous PIEZO1 activity, for instance within monolayers comprising cells with high and low PIEZO1 activity. To explore this, we extended the original model to investigate the impact of heterogeneous PIEZO1 activity in monolayers (we describe the model in Section 9 in S1 Text for details). This new model considers the migration of two cell types, each governed by its own set of equations with distinct model parameters, while interacting through cell-cell adhesion and volume-filling effects. Through simulations involving Control_cKO_ and *Piezo1-*cKO, as well as Control_GoF_ and *Piezo1-*GoF, we observed a correlation between the distribution of edge cells in the monolayers and the level of PIEZO1 activity. In particular, cells with reduced PIEZO1 activity (e.g., *Piezo1-*cKO) are over-represented at the wound edge (S19B Fig), which is consistent with the faster wound closure observed in homogeneous *Piezo1-*cKO monolayers. In contrast, cells with enhanced PIEZO1 activity (e.g., *Piezo1-*GoF) are underrepresented at the leading edge of the monolayer (S19D Fig), which is also consistent with the slower wound closure observed in homogeneous *Piezo1-*GoF monolayers. Testing these model predictions in experiments is deferred to future work.

Taken together, our experimental findings support our model predictions that PIEZO1 inhibits coordinated directionality during collective migration. Moreover, we identify that PIEZO1 activity negatively contributes to leader cell formation and the distance by which keratinocytes can coordinate their migration during 2D epithelial sheet migration.

## Discussion

Mechanical cues have been highlighted to play a prominent role in facilitating the coordinated polarization of individual cells within a collective, regulating the speed and coordinated directionality of collective migration [17]. We recently identified the mechanically activated ion channel PIEZO1 as being a key regulator of wound healing: keratinocytes with increased PIEZO1 activity exhibited delayed wound healing while decreased PIEZO1 activity resulted in faster wound healing [22]. Given PIEZO1’s role in wound healing, we explored PIEZO1’s effect on leader cell formation and coordinated directionality during collective keratinocyte migration. By taking a combined integrative mathematical modeling and experimental approach we identified that PIEZO1 activity suppresses leader cell formation, limits the coordinated directionality of cells during epithelial sheet migration, and reduces the distance by which keratinocytes can coordinate their directionality (Fig 6). This is the first time that PIEZO1 is seen to contribute to the correlation of cellular motions between neighboring cells which underlie the collective movements of cells during epithelial sheet migration. Given that PIEZO1 acts as a key mechanosensor in keratinocytes, this provides further evidence of the channel acting to couple mechanotransduction with correlated migration.

**Fig 6.**
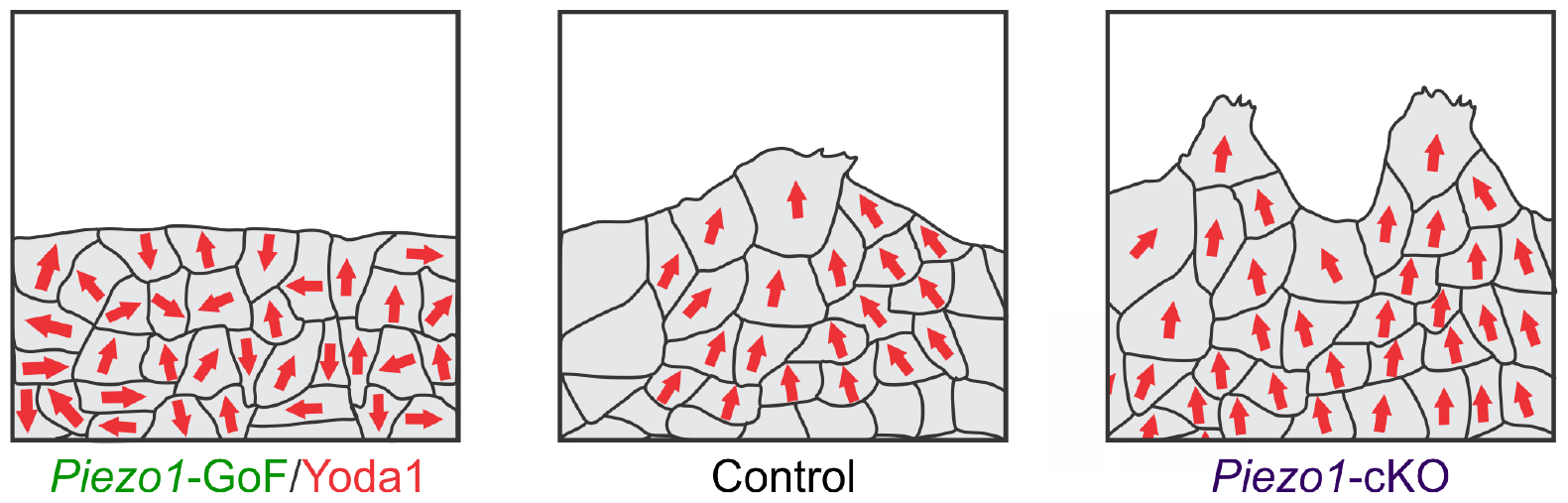
PIEZO1 activity inhibits spatial coordination and leader cell formation during collective migration. Summary schematic of collectively migrating monolayer of keratinocyte cells (gray) with direction of cellular motion overlaid (red arrows) under *Piezo1* -GoF/Yoda1 (*left*), Control (*middle*) and *Piezo1* -cKO (*right*) conditions. Note how as PIEZO1 activity is decreased, the coordinated direction of cells and number of leader cells increases.

When experimentally measuring persistence during single cell migration assays, we found that both *Piezo1* -GoF and *Piezo1* -cKO keratinocytes have increased persistence (Figure 2 & Figure 2—Figure Supplement 3, [22]), while Yoda1-treatment shows no effect on persistence (S6 Fig). On the other hand, measurement of persistence within collectively migrating cells shows that cells within both *Piezo1* -GoF and Yoda1-treated monolayers show less persistence while cells within *Piezo1* -cKO monolayers show an increase in persistence (Fig 4). Taken together, our experimental data indicates that PIEZO1’s effect on cell migration persistence is impacted by the contribution of neighboring cells on cell motion. Given that coordinated directionality is the result of cell-cell interactions, while persistence is an inherent characteristic of single cell migration it appears that coordinated directionality plays a key role in contributing towards the efficiency of collective migration experimentally.

In order to describe the inherent biological complexities underlying keratinocyte reepithelialization we adopted mathematical modeling as a tool to systematically investigate how aspects of collective cell migration affect wound closure. Through the development of a two-dimensional continuum model of wound closure derived through upscaling from a discrete model, we investigated how components of wound closure including cell motility, cell-cell adhesion, cell-edge retraction and the coordination of migration direction between cells, i.e., coordinated directionality, change with manipulation of PIEZO1 activity. Through numerical simulations, we incorporated experimental data to calibrate our model and match keratinocyte monolayer behavior. We examined how model parameters impacted two attributes of wound closure which we experimentally find are affected by PIEZO1 activity: the rate of wound closure and the edge length of simulated monolayers, which served as a measure of leader cell formation. From the modeling studies, the coordinated directionality of cells was identified as a key model parameter predicted to be impaired by PIEZO1 activity during wound closure.

Our model prediction guided the design of validation experiments and subsequent bioimage analyses, in which we confirmed the model prediction and demonstrated that PIEZO1 activity inhibits the ability of local subpopulations of cells to coordinate their movements across distances during collective migration. Altogether, we identified that PIEZO1 activity inversely correlates with the number of leader cells along the wound edge which in turn dictates the directed migration of cell collectives during keratinocyte reepithelialization. Taken together with our previous work demonstrating that enrichment of PIEZO1 at the wound edge triggers local retraction [22], we propose that PIEZO1-mediated retraction inhibits leader cell formation, which disrupts the uniform polarization of groups of cells and underlies the inhibition of collective migration during wound closure. This proposal is consistent with findings by other groups where pharmacologically increasing the contractile forces within monolayers was found to inhibit leader cell formation [25, 26, 37]. In addition, numerical explorations of collective cell migration during wound healing in scenarios where more than one PIEZO1 genotype is present add an intriguing dimension to our study and suggest future experiments combining cells of different genotypes to study the effect of homotypic and heterotypic interactions on cell migration and wound healing.

We developed our mathematical model to describe the dynamics of a straight scratch assay, which was the type of wound used in our experiments. However, for other wound geometries, the directional components of the model, such as the diffusion anisotropy, would need to be modified. In a circular wound, for example, the diffusion anisotropy would tend to be oriented in the radial direction corresponding to the alignment of cells moving radially inward toward the wound region. Furthermore, as a circular wound heals, the length of the wound would decrease over time. However, the roughness of the wound edge would increase, similar to the linear scratch assay considered here. In such a situation, rather than using the raw wound edge length as we do for simplicity, it would be necessary to normalize it, for instance by the perimeter of the circle.

During collective migration, the multicellular movement and corresponding polarization of cell clusters is dependent on signal transduction from leader cells to the ensuing follower cells [17, 34, 50]. Leader cells located at the front of these collectives transmit directional cues to follower cells through intercellular mechanical forces and biochemical cues which are communicated via cell adhesion molecules such as E-cadherin [27, 28, 51–55]. Both theoretical [40, 56] and experimental studies [57] have highlighted the role that cell-cell adhesions play in determining polarization dynamics and motility in multicellular systems. Given our finding that PIEZO1 activity inhibits leader cell formation and coordinated directionality it is possible that PIEZO1 coordinates mechanical forces communicated at cell-cell junctions during the collective migration of keratinocytes; however, further studies would be needed to elucidate this relationship. Consistent with this idea, recent work demonstrates interactions between cadherins and PIEZO1 at cell-cell junctions [58, 59].

Our previous work identified that PIEZO1 enrichment and activity induces cell retraction in single keratinocytes as well as along the wound edge of monolayers during *in vitro* scratch assays [22]. Building on these findings, we demonstrate here that monolayer conditions with elevated PIEZO1 activity lack leader cell formations and display reduced coordinated movement of cells. Interestingly, retraction forces generated by follower cells have been seen to promote the formation of leader cells along the wound edge [26]. Thus, it appears that collective migration requires carefully-regulated and coordinated levels of retraction. Consistent with this, Vishwakarma *et al*. found that pharmacologically adjusting the level of actomyosin contractility within monolayers affected the length-scale by which leader cells can correlate their forces such that actomyosin contractility levels inversely correlate with the frequency of leader cell formations [26]. We propose that altered patterns of PIEZO1-induced retractions within a monolayer may inhibit normal signal transduction by leader cells and disrupt cells from moving cohesively during collective migration. Given that these contractile forces could be communicated through cell-cell adhesions, patterns of cell contractility within the monolayer could be modeled to explore this by incorporating a variable adhesion coefficient in a PDE model or using a discrete approach such as a Vertex Model [60, 61].

The identity of downstream molecules underlying PIEZO1-mediated inhibition of keratinocyte migration during reepithelialization remains an open question. The Rho family of small GTPases, which includes the small molecules Rac1 and RhoA, play several roles during collective migration – regulating cell polarization, intercellular coordination of cellular movement, and leader cell initiation [14, 35, 62, 63]. Previous work has linked PIEZO1-mediated Ca^2+^ influx to impacting both focal adhesion dynamics [64–66] as well as Rac1 and RhoA levels. PIEZO1’s effect on small GTPases has been shown to affect migration in both neural crest cells [67] and cancer cells [68], Cadherin remodeling in lymphatic endothelial cells [69], and macrophage mechanotransduction in iron metabolism [70]. We also observed that total levels of Rac1 and RhoA in healing monolayers are reduced in Yoda1-treated compared to DMSO-treated samples (S12 Fig). While the downregulation of Rho GTPases provides an initial insight into the mechanism underlying PIEZO1-mediated inhibition of leader cells in collective migration, a detailed characterization of this relationship surpasses the scope of work covered within this paper. In future work, we can use mathematical modeling to investigate the relationship between PIEZO1 and Rho GTPases in keratinocyte collective migration by incorporating activator-inhibitor systems for Rho GTPase feedback networks [71] and spatial dynamics [72] into our modeling framework.

Since faster wound healing provides several physiological advantages to an organism, the role of PIEZO1 expression in keratinocytes may seem counterintuitive; however, other groups have reported that too many leader cells results in a disorganized epithelial sheet which affects the quality of wound closure [62]. Recent work examining wound healing in *Drosophila* found that knockout of the *Piezo1* orthologue, *Piezo*, resulted in poorer epithelial patterning and although wounds closed faster, they did so at the expense of epidermal integrity [73]. Therefore, it appears that effective wound healing may require a delicate balance of PIEZO1 activity.

PIEZO1 has been found to influence migration in other cell types, but whether channel activity inhibits or promotes cell migration has been seen to vary [64, 74–80]. Interestingly, recent studies found that PIEZO1 inhibition suppresses collective migration and results in a decrease in the coordinated directionality of migrating *Xenopus* neural crest cells [67, 78]. We note that the tissue-context of collective migration is known to engage distinct spatiotemporal signal transduction pathways [2, 17, 34]. Therefore, our seemingly contradictory findings to the observations in neural crest cells could reflect the inherent differences between the migration of neural crest cells and that of keratinocytes during reepithelialization. This highlights the need for studying PIEZO1 mechanotransduction under different biological contexts of cell migration.

Collective cell migration is an emergent phenomenon occuring at the multicellular level and stems from the large-scale coordination of individual cellular motions. Mechanical forces have been highlighted as playing an important role in shaping collective cell behaviors and influencing the formation and dynamics of both leader and follower cells [12, 14, 17]. Through this work, we have provided the first identification that the upregulation of PIEZO1 activity suppresses leader cell formation and inhibits both the coordinated directionality and the distance by which cells coordinate their cellular motion across length scales during epithelial sheet migration. Moreover, we develop a novel mathematical model for PIEZO1 regulated collective cell migration which is generalizable to studying the role of other proteins or cell types during epithelial sheet migration through analogous simulation and analyses. We propose that elevated PIEZO1-induced cell retraction inhibits the normal long-range coordination between cells during collective migration, disrupting typical mechanochemical activity patterns and the coordinated polarization of neighboring cells. Our findings provide a new biophysical mechanism by which PIEZO1 activity regulates the spatiotemporal dynamics across multiple cells to shape collective migration.

## Acknowledgments

We thank Dr. Ardem Patapoutian and his lab for the generous gift of *Piezo1* -cKO and *Piezo1* -GoF keratinocytes. We thank members of the laboratory for helpful comments on the manuscript.

## Methods and materials

### Ethics statement

All studies were approved by the Institutional Animal Care and Use Committee of University of California at Irvine and The Scripps Research Institute and performed within their guidelines.

### Animals

Keratinocyte samples from *Piezo1* -cKO and *Piezo1* -GoF mice were a gift from Dr. Ardem Patapoutian’s lab, the Scripps Research Institute. *Piezo1* -tdTomato reporter mice (*Piezo1* -tdTomato; JAX stock 029214), *Piezo1* -cKO and *Piezo1* -GoF mice were generated in previous studies [22, 81].

### Keratinocyte isolation and culture

Primary keratinocytes were isolated from the upper dorsal skin of P0-P5 mice as previously described [22]. Briefly, dissected tissue was allowed to dissociate for 15-18 hours. After dissociation, the epidermis was separated and incubated in Accutase (CellnTec CnT-Accutase-100) for 30 minutes at room temperature. Subsequently, the epidermis was transferred to a dish of CnT-Pr media (CellnTec), supplemented with 10% FBS and 1% penicillin/streptomycin where the epidermis was minced and then agitated using a stir plate for 30 min. After agitation, cells were strained through a 70 *μ*m cell strainer (Falcon). Strained cells were spun down and resuspended in CnT-Pr media (CellnTec) supplemented with ISO-50 (1:1000) (CellnTec) and Gentamicin (50 *μ*g/ml) (Thermo Fisher).

Isolated keratinocytes were seeded directly onto the glass region of #1.5 glass-bottom dishes (Mat-Tek Corporation) coated with 10 *μ*g/ml fibronectin (Fisher Scientific, CB-40008A). For single cell migration experiments, isolated cells were sparsely seeded onto the glass region at 1.5 × 10^4^ cells/dish while for monolayer scratch assay experiments, isolated cells were densely seeded onto the glass region at a density of 1.5x10^5^ cells/dish. One day after seeding, CnT-Pr supplemented culture media (see above) was switched to Cnt-Pr-D media (CellnTec) to promote keratinocyte differentiation. Keratinocytes were imaged 3 days after primary isolation, allowing at least 2 days for keratinocyte differentiation in Cnt-Pr-D media (CellnTec).

### Microscopy

For *in vitro* image acquisition, an Olympus IX83-ZDC inverted microscope equipped with a SOLA light engine (Lumencor) was utilized. For time-lapse imaging experiments, a full enclosure stage-top incubator system (Tokai Hit) enabled cells to be imaged at 37°C with 5% CO_2_ to maintain optimal cell health. *μ*Manager, an open-source microscopy controller software, was used for microscope hardware control and image acquisition [82, 83]. For all experimental data, images were taken using a UPlanSApo 10× dry objective with a numerical aperture of 0.40 and acquired using a Hamamatsu Flash 4.0 v2+ scientific CMOS camera.

### Immunofluorescence staining

For immunostaining of healing monolayers in S12 Fig, scratch wounds were generated in confluent monolayers of isolated keratinocytes and then treated with either 4 *μ*M Yoda1, or the equivalent concentration of the solvent DMSO, before allowing the monolayers to collectively migrate. 24 hours after initial wounding the monolayers, monolayers were fixed and then immunostained for total levels of Rac1 (S12 Fig, *left*) and RhoA (S12 Fig, *right*). Immunostaining was performed as previously described [84] using the following antibodies: Mouse anti-Rac1 (Millipore Cat#05-389-25UG, 1:200), Rabbit anti-RhoA (Proteintech Cat#10749-1-AP, 1:100), Donkey anti-Mouse 647 (Abcam Cat#AB150107, 1:500), Goat anti-Rabbit 488 (Life Sciences Cat#A32731, 1:500). Nuclei were stained by Hoechst (Invitrogen Cat#H1399) at 1*μ*g/mL for 5 minutes.

### Single cell migration assay

As previously described [22], time lapse sequences of DIC images were taken at 5 minute intervals. In brief, sparsely seeded keratinocytes were allowed to migrate for 16.67 hr at 37°C with 5% CO_2_ in fibronectin-coated glass-bottom dishes. Cell centroids were tracked using Cell Tracker (https://celltracker.website/index.html, Piccinini2016-dx) and resulting trajectories were analyzed using the cell trajectory analysis software, DiPer [47].

### Wound closure assay

Primary keratinocytes were cultured for 3 days until they formed a confluent monolayer. Prior to imaging experiments, cell nuclei were labeled by addition of SiR-Hoechst [45] (1 *μ*M; Cytoskeleton Inc.) to Cnt-Pr-D+1.2 mM Ca^2+^ bath media for 1 hour prior to imaging. As previously described, monolayer scratches were generated using a 10 *μ*l pipette tip and resulting cell debris was removed by performing three successive washes of culturing media [22, 85]. Time-lapse imaging series of wound closure were acquired by taking sequential DIC and fluorescence images at multiple positions. 1 *μ*M SiR-Hoechst remained in the Cnt-Pr-D+1.2 mM Ca^2+^ bath media throughout the imaging period. For Yoda1 experiments, 4 *μ*M Yoda1 or, as a control, the equivalent concentration of DMSO was supplemented to bath media prior to imaging. Leader cells display broad lamellipodia and are located at the front of protrusions along the leading edge of healing monolayers. During identification, leader cells were identified by manually reviewing time lapse image series and counting the number of cells located at the front of fingering protrusions at the leading edge which display increased polarization and large, prominent lamellipodia. Example leader cells identified during manual review are denoted by white arrows within S1 Fig. The number of leader cell formations is reported at the time point when either the wound interfaces touch or the imaging period finishes.

### Wound edge length analysis

Monolayer sheets were segmented from images taken during wound closure assays using a custom deep-learning based U-net architecture written in Python [86] (https://github.com/Pathak-Lab/PIEZO1-Collective-Migration). The length of the segmented wound edge was calculated by taking the cumulative euclidean distance between all detected pixel positions along the segmented monolayer leading edge. Due to any possible differences in edge length which might arise when manually making scratches in monolayers, each field of view’s edge length was normalized by dividing the edge length at T_final_, the time point when either the wound interfaces touch or the imaging period finishes by T_0_, the starting edge length at the starting time point for a field of view. This normalized edge length was used as a measure of the prevalence of leader cells along the wound edge for a given condition.

### Image analysis

Using the open-source image analysis software Fiji [87] the signal-to-noise ratio of SiR-Hoechst images was increased using Contrast Limited Adaptive Histogram Equalization (CLAHE) (https://imagej.net/plugins/clahe) prior to further analysis. For some images which had poor labeling of SiR-Hoechst, the denoising algorithm Noise2Void was also used to further increase the signal-to-noise ratio of nuclei images [88] (S13 Fig).

### Individual cell tracking

We combined the deep learning-based object detection method StarDist with the cell tracking software TrackMate to perform automated tracking of cells within monolayers [89–91]. Cell trajectories harvested using TrackMate were then exported for further analysis. Due to the technical limitations surrounding Microsoft Excel’s ability to handle large datasets, we developed Cell Pyper (https://github.com/Pathak-Lab/PIEZO1-Collective-Migration) a Pythonic analysis pipeline based on the open-source algorithm DiPer [47] to analyze the Mean Squared Displacement (MSD), Speed and Velocity autocorrelation of harvested cell trajectories.

For efficient computation of a trajectory’s MSD, MSDs are computed according to Eq. 18 (Eq. 4.11 in [92]) where *r*(*k*) *≡ r*(*k*Δ*t*) is a cell trajectory consisting of *N*_*t*_ timepoints and the MSD is calculated for timestep *m*.

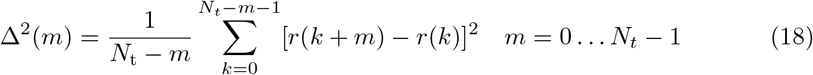

As described by Gorelik & Gautreau (Eq. 6 and 7 in [47], Velocity Autocorrelation analysis is calculated according to equations 19 and 20 for a trajectory consisting of *N* timepoints with a time-step of Δ*t* =5 min. A normalization factor (Norm; Eq. 19) is initially calculated for velocity vector *v*_*i*_ with starting coordinates (*x*_*i*_, *y*_*i*_) which is used to calculate the average velocity autocorrelation coefficient *v*_*ac*_ with step size *n*.

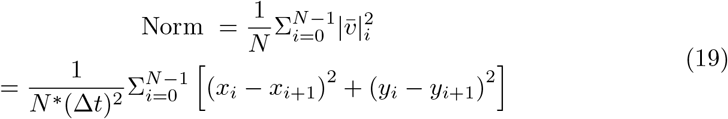

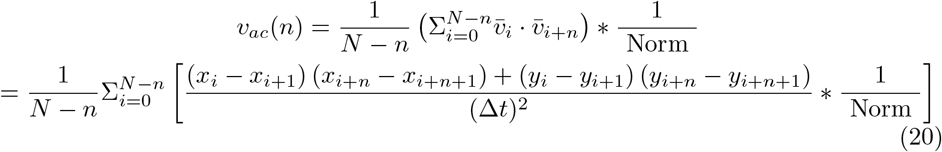

### Particle image velocimetry analysis

Particle Image Velocimetry (PIV) analysis was performed using the Python implementation of OpenPIV [93] (https://github.com/Pathak-Lab/PIEZO1-Collective-Migration). We use multiple passes of interrogation window sizes, initially using first-pass calculations with a 64 pixel x 64 pixel (55.2 *μ*m x 55.2 *μ*m) window followed by two iterations of 32 x 32 (27.6 *μ*m x 27.6 *μ*m) pixel windows and two iterations of 16 x 16 pixel (13.8 *μ*m x 13.8 *μ*m) windows. Each interrogation window was computed with a 50% overlap. A signal-to-noise filter (Threshold=1.3) was used on detected velocity vectors to remove any vector outliers. Outputs produced by OpenPIV analysis were then used to generate PIV flow fields as shown in Fig 5A-C. Working from the flow fields, individual PIV vectors were isolated and PIV vector direction was calculated and normalized to 0° to account for differences in angles of scratches made in monolayers (https://github.com/Pathak-Lab/PIEZO1-Collective-Migration). Vector direction distributions are illustrated as the probability density distribution across experimental replicates in Fig 5D-F. The von Mises distribution was employed to fit vector direction datasets by minimizing the mean squared error between the vector direction data and the von Mises probability density function. The resulting fitted curves represent the best approximation of the data by adjusting the parameters *μ* (mean) and *κ* (concentration or strength) of the von Mises distribution. The parameter *μ* represents the location where the distribution is clustered, while parameter *κ* indicates the level of directionality in our experimental context. The probability density function of the von Mises distribution for the vector direction angle *x* is expressed as:

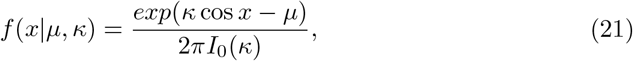

where *I*_0_(*κ*) represents the modified Bessel function of the first kind with order 0

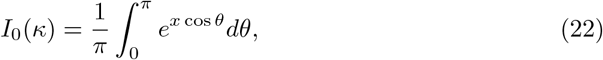

which is selected to ensure the distribution integrates to unity:

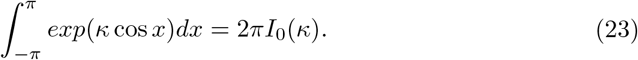

The variance of PIV vector directions within a field of view was calculated as the mean angular deviation, *z*, where *z* is defined in Eq. 24 (Eq. 2 in [48]). Outputs of this equation are bounded such that zero indicates no variability in vector direction within a flow field and one indicates high variability in vector direction.

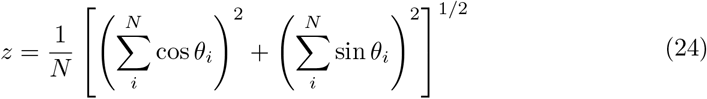

The spatial autocorrelation function, *C*, is computed according to Eq. 25 (Eq. 4 in [48]) using the radial velocity component of a given PIV vector, *v*, within a vector flow field at varying length scales, *r*.

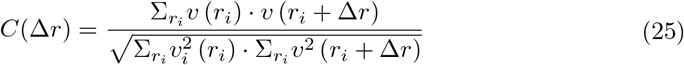

For measurement of the local autocorrelation in vector direction, the spatial autocorrelation at Δ*r* = 150 *μ*m was used to capture correlation of motion at multiple cell lengths. Length constants were calculated by using OriginLab to fit an exponential

function whose exponent is a 2^nd^ order polynomial (Eq. 26) to the spatial autocorrelation dataset and calculating the distance at which *C*(Δ*r*) *≈* 0.37.

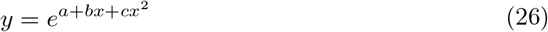

### Numerical scheme

In order to solve the governing equation (Eq. 4), we firstly carry out a forward time discretization (with size Δ*t*) on the left hand side *∂ρ/∂t* by (*ρ*(***x***_*i*,*j*_, *t* + Δ*t*) *− ρ*(***x***_*i*,*j*_, *t*))*/*Δ*t*. In terms of space discretization (right hand side), the transitional probability is proved to be separable (Eq. 3) in the discrete model, which allows us to work on the diffusion part and advection part separately: for the diffusion part, a natural discretization is directly given by the discrete model (e.g., centered finite differences); for the advection part, we apply a 2nd order weighted essentially non-oscillatory (WENO) method [94, 95] to discretize the equation. Hence, an explicit finite difference scheme was used to update the cell density at the nth time step 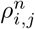 iteratively on the simulation domain [0, 1] *×* [0, 1] until wound closure.

### Model parameter adjustment

In Figs 3B-D and S7, the experimentally derived “Single Cell Migration” dataset guides the changes in model parameters of retraction strength, retraction duration, inter-retraction duration and cell motility when PIEZO1 activity is altered. However, cell-cell adhesion and coordinated directionality were not measured directly in the experiments and instead are inferred by trying to match model and experimental results.

While both cell-cell adhesion and coordinated directionality are designed to range from 0 to 1 in our model, the feasible adhesion coefficient actually needs to be bounded above by 0.66 in order for the diffusivity to be positive definite (see Section 3 in S1 Text for detailed derivation). Since the dependence of wound closure rate and wound edge length with respect to individual model parameters was already numerically shown to be a monotonic function of these parameters (Figs 2G, 2H and S3), it is sufficient to directly use the extrema of the model parameters: 1 for increased coordinated directionality, 0.66 for an increased adhesion, and 0 in the case that coordinated directionality and/or adhesion is decreased. For example, when matching experiments and simulations requires an increased coordinated directionality, we take *w*_*A*_ = 1. Because of the dependency of the outcomes (wound closure rate and edge lengths), if increasing a model parameter to its maxima fails to match the experimental trends, it would be impossible to match with smaller values.

The base values of cell-cell adhesion and coordinated directionality are taken to be 0.2 and 0.4, respectively. In Fig 3B-D, the adhesion coefficient is fixed at the value 0.2, while in S7 Fig, coordinated directionality is fixed at the value 0.4. The base values of the model parameters can be found in S4 Fig.

### Statistical analysis

*P* values, statistical tests, and sample sizes are declared in the corresponding figures. All datasets were tested for normality using the Shapiro-Wilk test prior to statistical analysis. The two-sample t-test was used where data were modeled by a normal distribution and a nonparametric test was used in the case of non-normal distributions. Cumming estimation plots were generated and Cohen’s *d* value was calculated using the DABEST python [96] (https://github.com/ACCLAB/DABEST-python). The Cohen’s *d* effect size is presented as a bootstrap 95% confidence interval (95% CI) on a separate axes. *p* values for Fig 5G-I are declared in S11 Fig.

## S1 Text

### Section 1. Boundary conditions of governing equation

On the Dirichlet boundaries *y* = 0 and *y* = 1 (Eq. 17), the cell density is determined by functions *g*_1_(*x, t*) and *g*_2_(*x, t*) which are continuous on [0, 1] *×* [0, + *∞*). Since both of these are randomly generated from the same approach, without loss of generality, let’s say *g*(*x, t*). Covering [0, 1] *×* [0, +*∞*) with a grid, taking mesh sizes *h*_*x*_ and *h*_*t*_ and labeling grid nodes (*x, t*) = (*x*_*i*_, *t*_*j*_) = (*ih*_*x*_, *jh*_*t*_) by indices (*i, j*), the function values at grid points *g*_*i*,*j*_ = *g*(*x*_*i*_, *t*_*j*_) are taken to follow a normal distribution

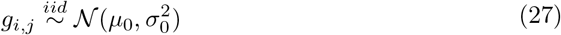

with mean *μ*_0_ and standard deviation *σ*_0_ (*μ*_0_ = 0.6 and *σ*_0_ = 0.3 were adapted in the simulation). This models the variability of the influx of cells from the monolayer moving into the wound region. Thereafter, the function *g*(*x, t*) is given by an interpolation on *g*_*i*,*j*_. Specifically, the boundary conditions on *y* = 0 and *y* = 1 are classical Dirichlet boundary conditions with a constant influx *μ*_0_ if *σ*_0_ is set to be 0.

### Section 2. Initial condition of governing equation

Assume *u*(*x, y, t*) is a function defined on [0, 1] *×* [0, 1] *×* [0, + *∞*) and satisfies the following diffusion equation

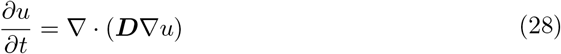

with the same diffusivity ***D*** as in Eq. 5 and the same boundary conditions as in Eq. 17:

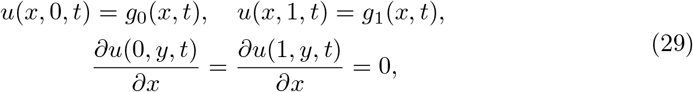

while the initial condition is globally zero:

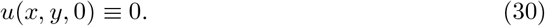

With this setting, the wound region (*u* = 0) is narrowing down from the whole square domain [0, 1] *×* [0, 1] to a heterogeneous horizontal banded region in the middle of the domain, before finally shrinking to zero area and disappearing at *t* = *t*_*end*_. At a certain time point *t* = *t*_0_ *∈* (0, *t*_*end*_) during this process,we set *ρ*(*x, y*, 0) = *u*(*x, y, t*_0_) as the initial condition of our governing equation Eq. 4.

In other words, this initial condition is generated by the governing equation (Eq. 4) but without retraction, starting from zero initial values and diffusing cells without any retraction for a period of time, until retractions were introduced. At the moment right before the first retraction, cell densities across the domain [0, 1] *×* [0, 1] are the initial values for the governing equation. This enables us to start with a variable, and more physiological, initial condition compared to taking a constant values at the wound edge.

### Section 3. Positive definite diffusivity

The matrix *d ·* (*w*_*I*_ ***I*** + *w*_*A*_***A***) is diagonal and has a positive spectrum. Therefore, the diffusivity 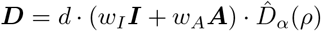 of the governing equation (Eq. 4) is positive definite if and only if the scalar diffusion coefficient 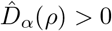, which depends on the value of adhesion coefficient *α*. By inspecting this 5-th degree polynomial, we see that 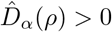 unconditionally holds for all levels of cell density *ρ ∈* (0, 1) as long as

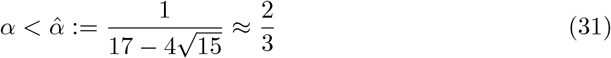

with a critical value 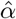. When 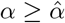, there exists an interval

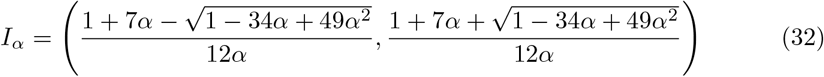

such that 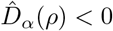 if and only if *ρ ∈ I*_*α*_. That is, the diffusivity is negative definite when cell density *ρ ∈ I*_*α*_, which results in the ill-posedness of the initial value PDE problem. As *α →* 1, the interval *I*_*α*_ expands from a single point 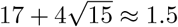 to *I*_1_ = (1*/*3, 1).

### Section 4. Retraction is modeled by advection

Performing a Taylor expansion on the cell density *ρ*, centered at ***x*** = ***x***_*i*,*j*_, in the discrete master equation (Eq. 1) without specifying 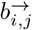, we have

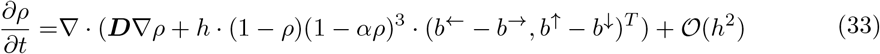

where ***D*** is the same diffusivity as in Eq. 5. By taking *h →* 0, the continuum limit would be a simple diffusion equation *∂ρ/∂t* = *∇ ·* (***D****∇ρ*) without an advection term, unless both Δ*b*^*↔*^ = *b*^*←*^ *− b*^*→*^ and Δ*b*^*↕*^ = *b*^*↑*^ *− b*^*↓*^ are *𝒪* (1*/h*), the advection scaling. Therefore, we define 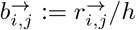 with 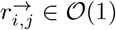. By taking *h →* 0 under this setting, Eq. 33 turns into our continuum limit (Eq. 4), where the retraction is modeled by advection.

### Section 5. Function smoothing

To localize the retraction region (Eq. 10), we smooth the Heaviside function *H*(*γ − ρ*) (Eq. 11) using a hyperbolic tangent function

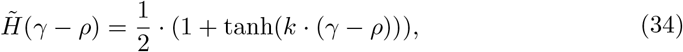

where *k* is the steepness level at transition point *ρ* = *γ* (*k* = 10 was adapted in the simulations). On the other hand, the indicator function 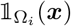 is smoothed using a 2D generalized bell-shaped function:

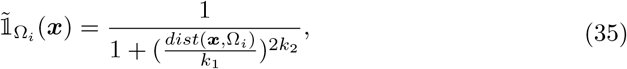

where *k*_1_ and *k*_2_ are parameters determining the width and steepness of the transition region in the smoothing process. The distance between a point ***x*** and a set Ω_*i*_ in 2D Euclidean space is induced by a natural 2-norm *∥ · ∥*:

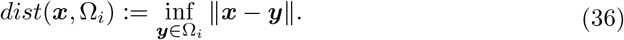

Since the region Ω_*i*_ is banded, the indicator function 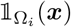 is equivalent to its 1D form 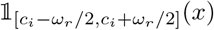 (Eq. 12). Therefore, the 2D generalized bell-shaped function 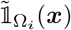 (Eq. 35) can be simplified into a 1D version:

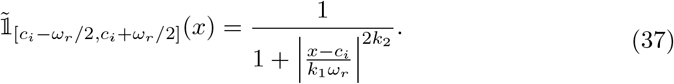

By adjusting the center and the width of the characteristic interval, the generalized bell-shaped function given above can be applied to smooth the indicator function in time 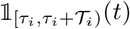 as the following:

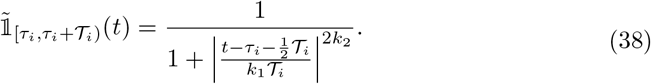

Since the spatial regions of retractions decay away before the next retraction event occurs, shifts in the retraction region *c*_*i*_ *∼ 𝒰* (0, 1) do not introduce discontinuities. Note that the selection of width and steepness parameters (*k*_1_ and *k*_2_) for smoothing indicator functions are different for the spatial and temporal localizations of retraction (S4 Fig).

### Section 6. Model dimensionalization

Recall that Eq. 1 is our non-dimensional master equation with the transitional probability Eq. 2. In order to relate the dimensions in the model to the experiments, we take the dimensional variables to be (1) 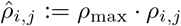 (*ρ*_max_ is the maximal cell density), (2) 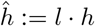 (*l* is the characteristic length) and (3) 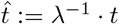 (*λ*^*−*1^ is the characteristic time). Hence, the dimensional transitional probability becomes

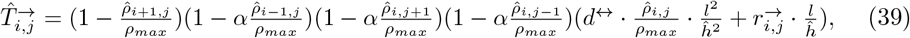

which can be taken into the master equation (Eq. 1) with the dimensional time derivative

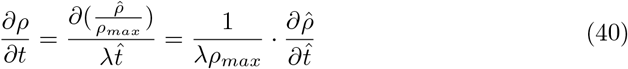

and obtain the continuum limit by taking 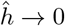 :

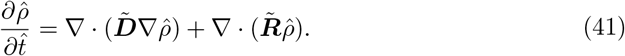

Here, 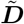 is the dimensional diffusivity (diffusion coefficient) given by

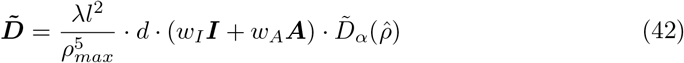

With

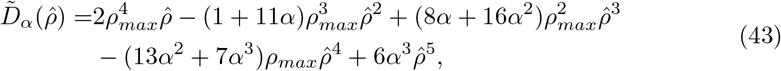

and ***I*** = ***I***_2_, ***A*** = *diag*(0, 1) are defined as before in Eq. 5. On the other hand, 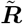 dimensionalized retraction (advection velocity) given by

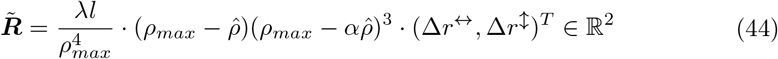

where Δ*r*^*↔*^ and Δ*r*^*↕*^ are defined as in Eq. 10.

In order to connect the model with the experiments and to calculate the effective cell diffusion coefficient as well as the advection velocity, we need to know 3 parameters: *λ, l* and *ρ*_max_. We notice that *λl*^2^ = *v· l*, where *v* := *λl* is actually the characteristic velocity (length over time). Hence, if we have a measurement of the characteristic velocity and the length scale, we can determine the characteristic time by *λ* = *v/l*. In conclusion, we can connect our theory and numerical parameters with the biological experiments in the following way:

- Maximal cell density *ρ*_max_ and dimensional cell density 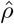: here 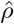 is interpreted as a number density, i.e., 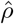 *dxdy* is the number of individuals with the position in the phase area *dxdy* centered at (*x, y*). We can quantify this from the experimental results: put down a grid, count the number of cells in each single square and get a spatial representation of the cell density. In the monolayer region away from the front edge, we expect the cell density should be nearly uniform spatially and temporally, and that value could be used for *ρ*_max_.
- Characteristic length *l*: we define the characteristic length scale to be the distance from the wound edge to the region where the cells reach the maximal density in the monolayer. In our numerical tests, we did not simulate the whole experimental domain, instead, our simulation focused on the region of transition, that is, the region in which the cell density transits from the front to the maximum. Hence, our computational domain is a small region around the wound edge (*∼* 10 cell lengths).
- Characteristic velocity *v*: the velocity of the moving front can be used for this, by averaging the front advancing speed measured by cell shape analysis.
- Characteristic time *λ*^*−*1^: since we already have the way to determine the characteristic length *l* and velocity *v*, the characteristic time can be derived directly by *λ*^*−*1^ = *l/v*.

With the measurements mentioned above, we are able to calculate *ρ*_max_, *l, v, λ* and hence the diffusion coefficient and retraction velocity. At this point, we do not have a direct measurement for the adhesion coefficient *α*. A direct measurement for the cell-cell adhesion is being considered in our future work.

### Section 7. Alternative model: a fully continuum approach

Recall that our model was initially formulated at the discrete level and subsequently upscaled into a continuous PDE. Here, we test a phenomenological continuum model in which cell-cell adhesion is postulated at the continuum level rather than being obtained by upscaling.

In this new model, the governing equation is still a diffusion-advection equation in the same form as Eq. 4, but the hindering effect of cell-cell adhesion on collective migration is modeled by reducing the overall diffusion coefficient as adhesion increases, which is consistent with the approach used by Amereh *et al*. in [44]. While still accounting for coordinated directionality, volume filling effects and the advancing front connecting the wound and the monolayer, the diffusion coefficient in this new continuum model can be specifically expressed as

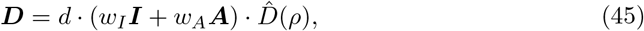

where the overall structure mirrors the diffusion coefficient in our original model (Eq. 5). However, the scalar diffusion coefficient, denoted as 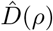, becomes a quadratic polynomial of cell density without depending on any additional parameters:

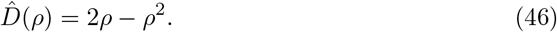

Compared with the original model, *d* in Eq. 45 now contains the combined effects of cell motility and cell-cell adhesion.

Analogously, we assume that the advection velocity would now model the combined effects of retraction strength and cell-cell adhesion. In particular, the advection term in Eq. 4 now becomes

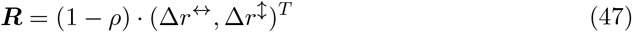

where Δ*r*^*↔*^ and Δ*r*^*↕*^ are defined in the same way as in Eq. 10 but incorporating the combined effects of retraction strength and cell-cell adhesion.

To compare the model predictions with our original model, we replicated all the previous simulations related to *Piezo1* -cKO, *Piezo1* -GoF and Yoda1 using this new phenomenological model. Similar to the calibration process for the original model, we adjusted model parameters based on experimental data (S5 Fig). This involved varying model parameters from a wild type (e.g., Control_GoF_) to a PIEZO1 phenotype (e.g., *Piezo1* -GoF), and measuring changes of wound healing metrics (wound closure and edge length) from repeated simulations. In line with our previous findings, we observed that the simulation results from the Control_GoF_ to *Piezo1* -GoF case are only able to replicate experimental observations if coordinated directionality is reduced. That is, by reducing the parameter of coordinated directionality, we recapitulated the experimental phenotype of both a shorter edge length and slower wound closure in simulated *Piezo1* -GoF monolayers (S14A Fig). Importantly, we noted that changes to the diffusion coefficient, according to changes in cell-cell adhesion, alone failed to replicate all experimental results (S15 Fig), consistent with the results obtained using the upscaled model of adhesion. This underscores the primary role of coordinated directionality in PIEZO1’s impact on reepithelialization and reaffirms our main conclusion that PIEZO1 activity hinders coordinated directionality. Because the dependence of the diffusion coefficient and the retraction strength on cell-cell adhesion could be quite different quantitatively, as suggested by our upscaled model (see Eq. 8 and Eq. 9), for simplicity, here we focused only on the changes in *d* and not on Δ*r*^*↔*^ and Δ*r*^*↕*^. However, we varied the retraction strengths and durations in the context of the original upscaled model (see Section 8 in S1 Text for details) and reached the same conclusion.

### Section 8. Robustness testing for model calibration

The experimental data used for model calibration can be categorized into two main components: cell motility and retraction processes (including retraction duration, inter-retraction duration, and retraction strength). In the process of model calibration, we utilized experimental data at the single-cell level from Table 2 and S5 Fig. To test whether our conclusions depend on the quantitative single cell data, we varied the magnitudes of the motility and retraction processes.

For cell motility, we used experimental data from the monolayers (S9 Fig), which shows that cell motility within the monolayer increased in *Piezo1* -cKO and decreased in *Piezo1* -GoF and Yoda1-treatment compared to their respective experimental controls. Calibrating our original model using motility measured from monolayers, together with the original magnitudes of the retraction processes, and maintaining cell-cell adhesion and coordinated directionality as observed in Control_GoF_, we found that while we could replicate simulated monolayers of *Piezo1* -GoF keratinocytes exhibiting slower wound closure compared to simulated Control_GoF_ monolayers, but we failed to observe the decrease in simulated monolayer edge length seen in experiments (S16A Fig). However, by reducing the parameter of coordinated directionality, we recapitulated the experimental phenotype of both a shorter edge length and slower wound closure in simulated *Piezo1* -GoF monolayers (S16A Fig). On the other hand, our model simulations, calibrated using the monolayer cell motility dataset along with original retraction processes for both *Piezo1* -cKO and Yoda1-treated keratinocytes, reproduced the observed experimental trends. However, with adjustments to coordinated directionality, we observed a more pronounced effect (S16B and S16C Fig). Importantly, we noted that changes to cell-cell adhesion parameters alone failed to replicate all experimental results, underscoring the primary role of coordinated directionality in PIEZO1’s impact on reepithelialization (S17 Fig).

For retraction processes (retraction duration, inter-retraction duration and retraction strength), our previous work [22] indicated qualitative consistency in between single cell data and monolayer experiments. For example, in both single cell and monolayer experiments, retractions in the presence of Yoda1 consistently exhibit shorter durations and larger magnitude retractions compared to DMSO, albeit with variations in the degree of change. Building upon this qualitative observation from experiments, we conducted additional simulations. In these simulations, we recalibrated our original model based on general qualitative trends rather than specific quantitative values for retraction. For instance, Yoda1-treated cells exhibit significantly stronger retraction, with a strength approximately three times (around 2.87) that of its control DMSO-treated cells (S5 Fig). Instead of adhering strictly to this specific ratio of 2.87, we performed simulations using two additional ratios, one larger and and one smaller, while maintaining the qualitative trend in which the retraction strength in Yoda1-treated cells is greater than in DMSO-treated cells. This approach was also applied to the calibration of the retraction duration and the inter-retraction duration, with adjustments made in various ratios rather than relying on specific quantitative values derived from experimental statistics (S5 Fig). Again, two additional values of the durations were used.

Applying this recalibration to simulated monolayers of *Piezo1* -GoF keratinocytes, we observed slower wound closure compared to Control_GoF_ monolayers; however, the expected decrease in simulated monolayer edge length was not observed (S18 Fig). Consistent with our previous findings, simulations from the Control_GoF_ to *Piezo1* -GoF case could replicate experimental observations only when coordinated directionality was reduced. Specifically, by lowering the parameter of coordinated directionality, we recapitulated the experimental phenotype of both a shorter edge length and slower wound closure in simulated *Piezo1* -GoF monolayers, while changes to cell-cell adhesion parameters failed to replicate all experimental results (S18 Fig). The results from this recalibrated model reaffirm the role of coordinated directionality, leading to the same fundamental conclusion that PIEZO1 activity hinders coordinated directionality.

### Section 9. Heterogeneous cell collective migration model

We expanded the original model to explore the influence of PIEZO1 activity in mixed populations. This model considers the migration of two distinct cell types, each governed by its own set of equations and parameters, while interacting through cell-cell adhesion and volume-filling effects. Specifically, the cell densities are denoted as *u* and *v*, and hence there are three types of cell-cell adhesions: (1) between *u* cells (*α*_*uu*_), (2) between *v* cells (*α*_*vv*_), and (3) interaction between *u* cells and *v* cells (*α*_*uv*_). We assume that the interaction adhesion *α*_*uv*_ hinders collective cell migration in the same way as *α*_*uu*_ and *α*_*vv*_. Regarding volume-filling effects, the migration of either *u* cells or *v* cells is impaired by the total density of cells (*u* + *v*) in the front. The spatial position of wound edges are determined by the interface connecting the total cell *u* + *v* and the cell-free region. However, retractions near wound edges are applied individually for *u* cells and *v* cells, following their respective retraction parameters, including the duration and strength of retraction, as well as the inter-retraction duration.

Following the same framework as our original homogeneous cell model, the heterogeneous collective cell migration occurs within a square domain, which is defined by two opposite sides with zero-flux Neumann boundaries and two opposite sides with randomized Dirichlet boundaries. Source cells migrating into the domain from randomized Dirichlet boundaries exhibit stochastic influx and stochastic proportions of cells, with the mean of the *v* cell proportion among source cells designated as *p*_*v*_. The initial distribution of total cells *u* + *v* aligns with our original model (see Section 2 in S1 Text), where cells are evenly mixed with a *p*_*v*_ percentage of *v* cells and consequently, a 1 *− p*_*v*_ percentage of *u* cells.

To simulate monolayers with a mixture of Control_cKO_ and *Piezo1* -cKO cells, we used parameters for the Control_cKO_ and *Piezo1* -cKO cells from the homogeneous monolayers we simulated previously. This ensures that *u* cells exhibit migration behavior akin to Control_cKO_, while *v* cells mirror the characteristics of *Piezo1* -cKO. Throughout this mixed collective migration process, we varied the proportion of *Piezo1* -cKO cells in initial and source cells (*p*_*v*_), and we observed that cases with a higher *Piezo1* -cKO percentage displayed faster wound closure (S19A Fig), which is consistent with the faster wound closure observed in homogeneous *Piezo1* -cKO monolayers. Additionally, *Piezo1* -cKO cells tended to advance to the front and aggregate around the wound edge (S19A Fig).

To quantify this effect, we measured the percentage of *Piezo1* -cKO cells among all wound edge cells at every time step during the wound closure process, and subsequently calculated the average. The term “wound edge cells” pertains to cells situated near the wound edge, where the total density satisfies the condition 0 *< u* + *v < γ*_edge_. To determine the threshold *γ*_edge_ for identifying wound edge cells, we conducted simulations by mixing Control_cKO_ and Control_cKO_ rather than mixing Control_cKO_ and *Piezo1* -cKO. That is, both *u* and *v* cells have the identical phenotype and parameters drawn from the Control_cKO_ experimental data. In this case, for any fraction of *v* in the entire monolayer, we expect to observe the same fraction of *v* cells in wound edge cells where *u* + *v < γ*_edge_. The minimal *γ*_edge_ that satisfies this condition is the threshold we sought, and it was determined to be 0.2. The results revealed that *Piezo1* -cKO cells are over-represented at the wound edge (S19B Fig).

Analogously, we investigate the mixing of Control_GoF_ and *Piezo1* -GoF. Guided by the findings from our original model, in addition to adjusting retraction-related parameters and cell motility based on experimental data (S5 Fig), we also decrease the parameter of coordinated directionality in *Piezo1* -GoF (*v* cells) compared to Control_GoF_ (*u* cells). By varying *p*_*v*_, representing the proportion of *Piezo1* -GoF cells in initial and source cells, we observed that scenarios with a higher *Piezo1* -GoF percentage exhibited slower wound closure (S19C Fig), which is consistent with the slower wound closure observed in homogeneous *Piezo1* -GoF monolayers. Quantitative analysis revealed that, in contrast to the *Piezo1* -cKO case, *Piezo1* -GoF cells are underrepresented at the leading edge of the monolayer (S19C and S19D Fig).

Further insights into this heterogeneous cell migration model, a comprehensive exploration of the intricacies and findings derived from this study, will be provided in future work.

---

**S1 Fig.**
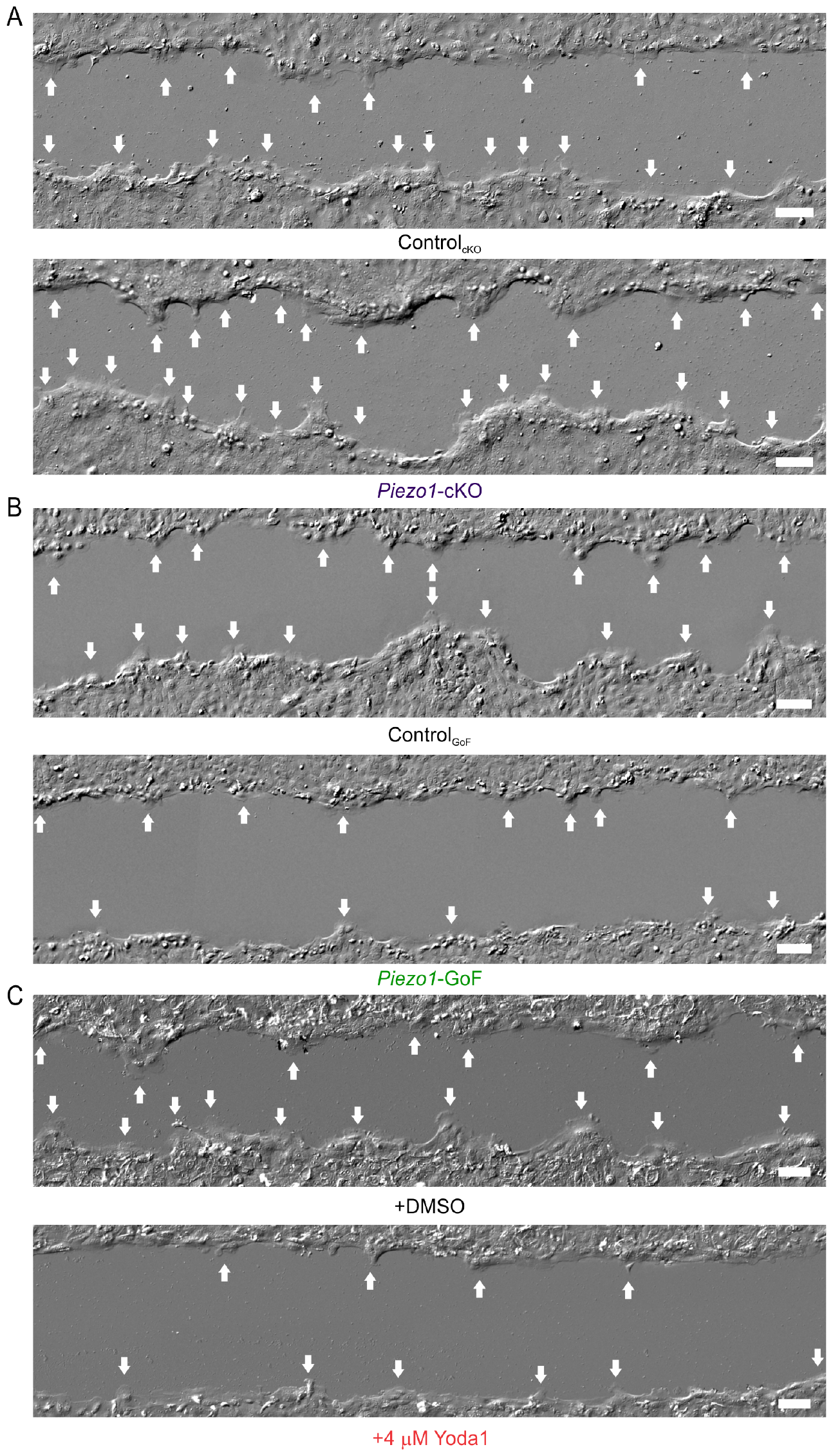
PIEZO1 inhibits leader cell formation at wound margins. Representative DIC images of wounds generated in (**A**; *top*) Control_cKO_, (**A**; *bottom*) *Piezo1* -cKO, (**B**; *top*) Control_GoF_, (**B**; *bottom*) *Piezo1* -GoF, (**C**; *top*) DMSO-treated and (**C**; *bottom*) 4 *μ*M Yoda1-treated monolayers. White arrows indicate leader cell protrusions. Representative images were taken at the same time point as the respective control field of view. Scale bar = 100 *μ*m. Related to Fig 1.

**S2 Fig.**
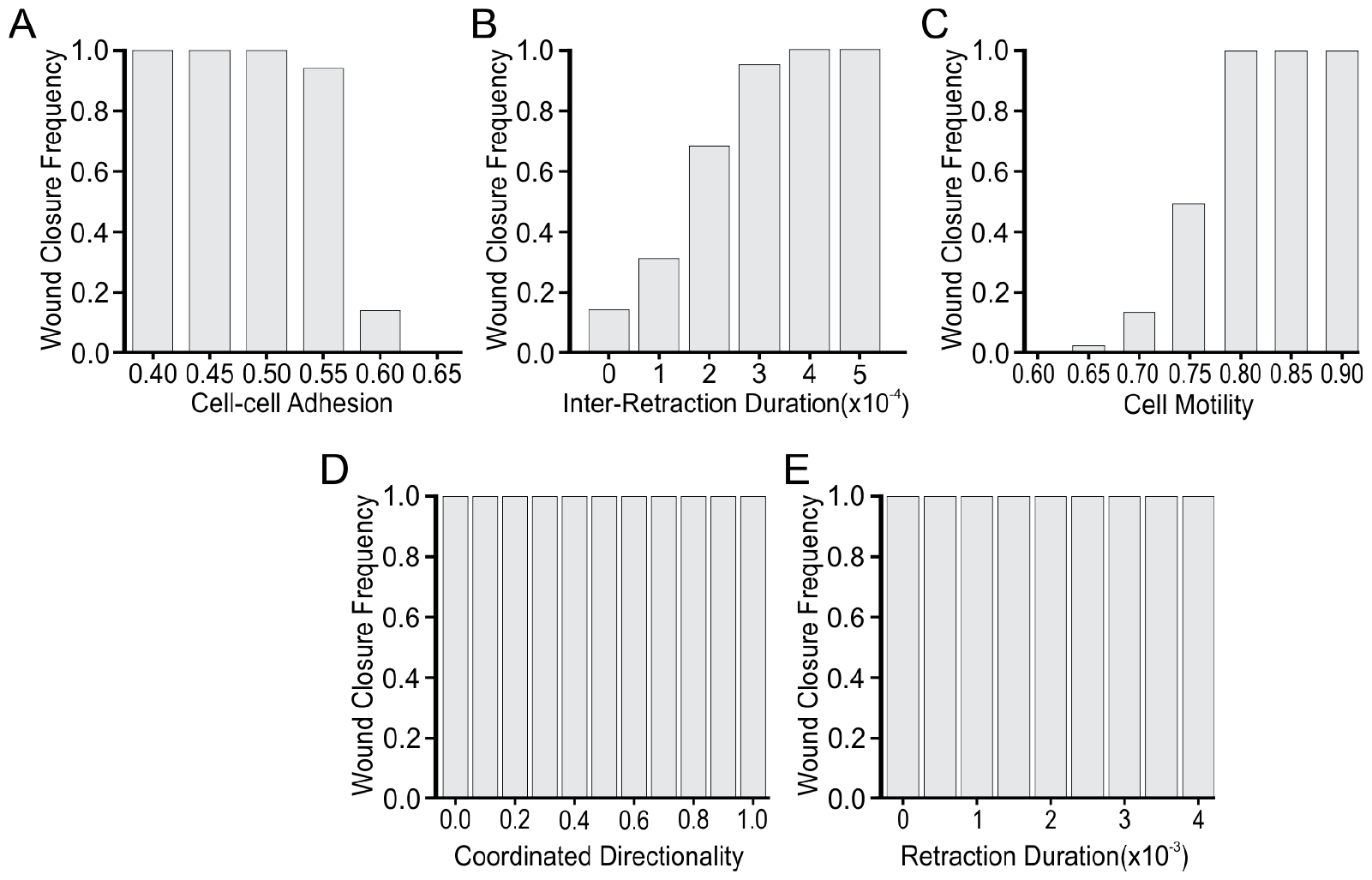
Wounds fail to reach closure if parameter values exceed reasonable ranges. **(A)** The percentage of wound closure cases under different levels of cell-cell adhesion. **(B, C, D, E)** Similar to (A) but for inter-retraction duration, cell motility, coordinated directionality and retraction duration respectively. Related to Fig 2F.

**S3 Fig.**
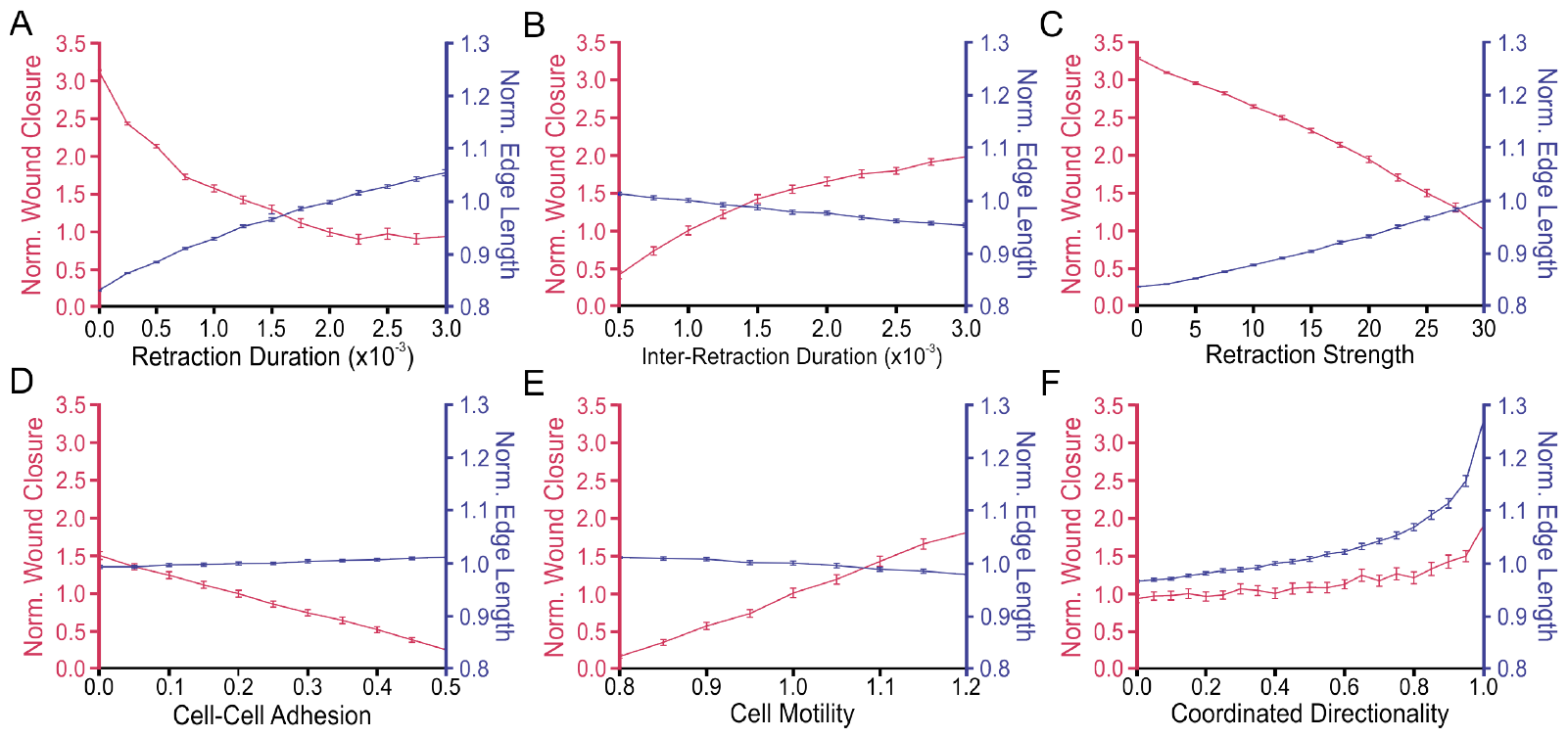
Coordinated directionality is the only parameter which replicates all experimental results. **(A)** The mean of 100 simulation results showing the effect of retraction duration on normalized wound closure (*red; left axes*) and edge length (*blue; right axes*). Error bars depict the standard error of mean. **(B-F)** Similar to (A) but for inter-retraction duration, retraction strength, cell-cell adhesion, cell motility and coordinated directionality, respectively. The data in C and F are also shown in Fig 2G and Fig 2H but are reproduced here for ease of comparison.

**S4 Fig.**
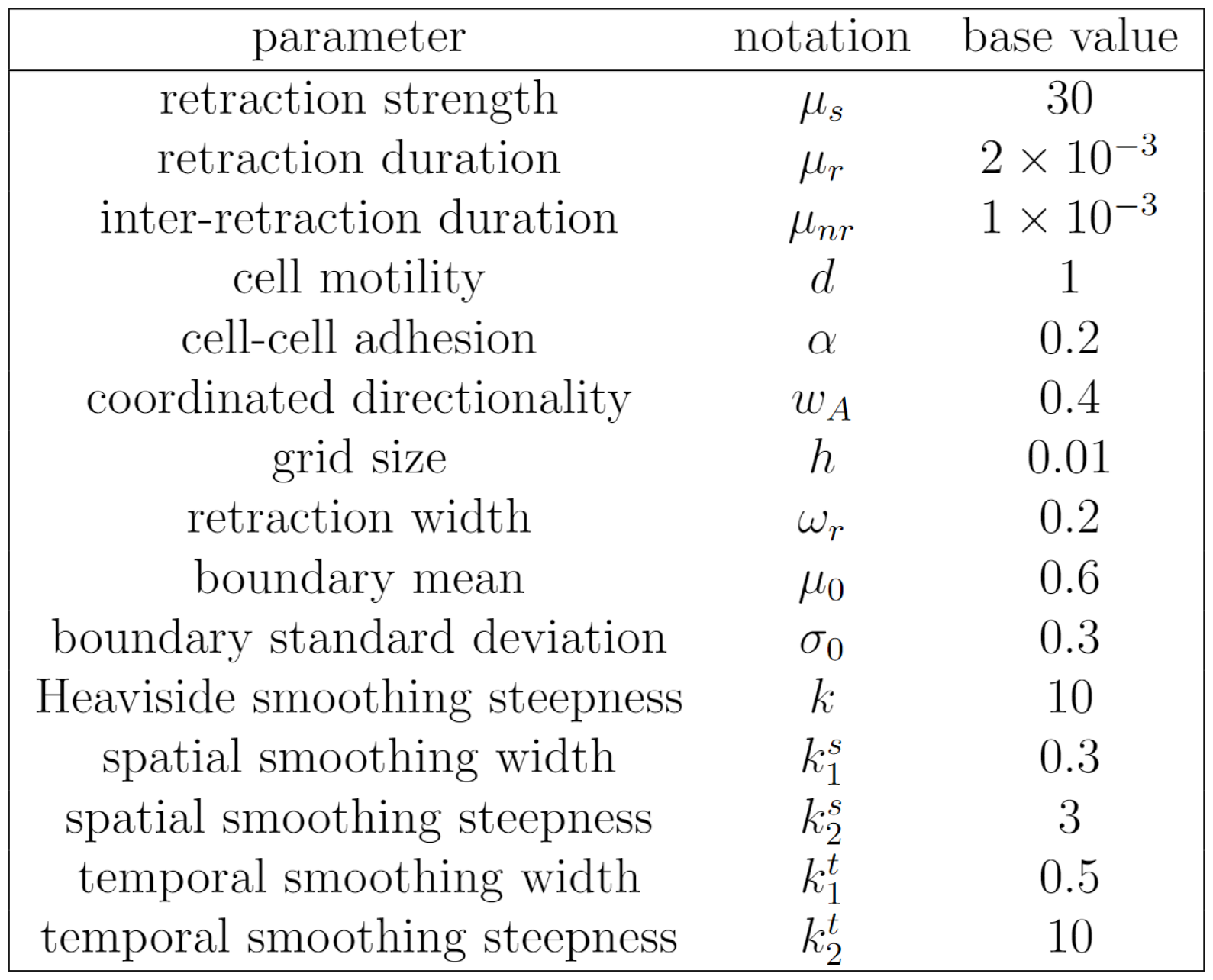
List of model parameters and their base values.

**S5 Fig.**
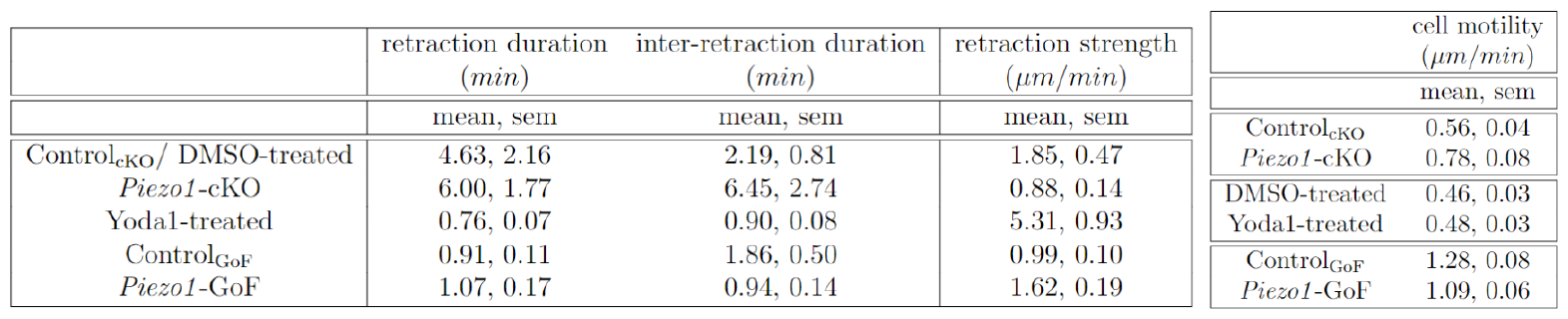
PIEZO1 activity affects single cell migration. Mean and standard error of mean (sem) of single cell migration dataset (retraction duration, inter-retraction duration, retraction strength and cell motility) under different experimental conditions. Retraction duration data was derived by kymograph measurements, retraction strength derived from cell shape analysis and cell motility data from tracking cells during single cell migration assays [22].

**S6 Fig.**
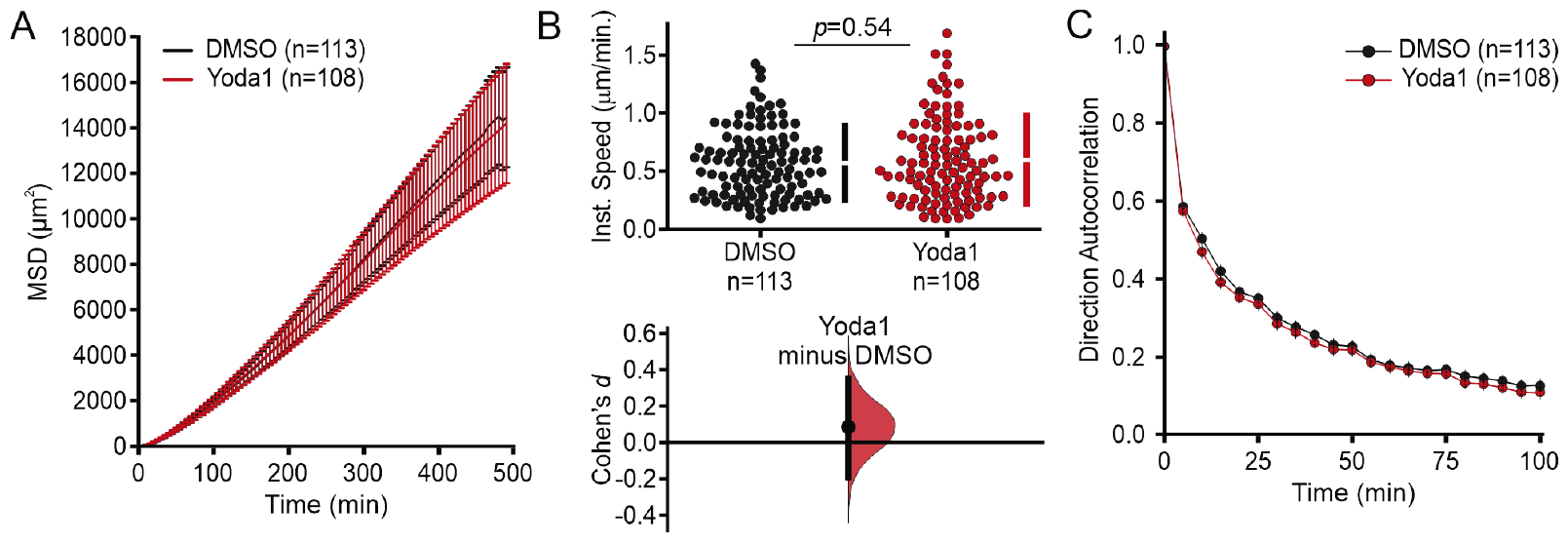
Yoda1 has no effect on single cell migration. **(A)** Mean Squared Displacement (MSD) analysis of Yoda1-treated keratinocytes. Average MSD plotted as a function of time. **(B)** Cumming plot illustrating quantification of the average instantaneous speed from individual Yoda1-treated keratinocytes plotted against DMSO-treated Control (Cohen’s *d* = 0.08; *p* value calculated via two-sample t-test). **(C)** Average direction autocorrelation of Yoda1-treated keratinocytes relative to DMSO-treated control cells plotted as a function of time. *n* in A-C denotes the number of tracked individually migrating keratinocytes for each condition. Related to Table 2; S5 Fig.

**S7 Fig.**
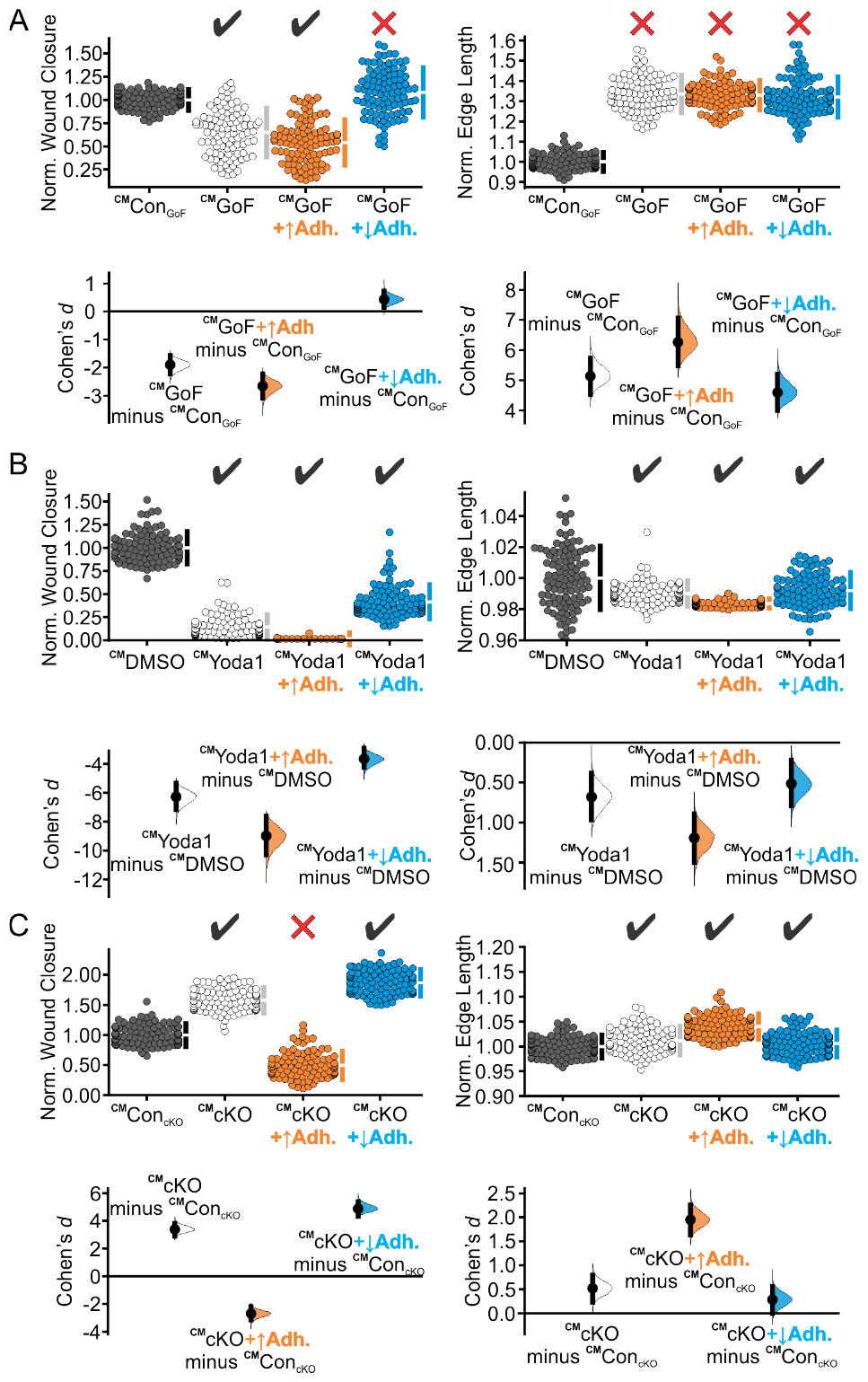
Varying cell-cell adhesion fails to match all the experimental trends. **(A)** Cumming plots showing simulation results in which we use our calibrated model (^CM^) to predict how PIEZO1 affects wound closure (*left column*) and wound edge length (*right column*) in simulated Control_GoF_ monolayers (*gray*), *Piezo1* -GoF monolayers without altered adhesion parameters (*white*), *Piezo1* -GoF monolayers with increased cell-cell adhesion (*orange)* and decreased cell-cell adhesion (*blue)*. **(B)** Similar to A but using simulation results from DMSO-treated monolayers (*gray*), Yoda1-treated monolayers without altered adhesion parameters (*white*), Yoda1-treated monolayers with increased cell-cell adhesion (*orange)* and decreased cell-cell adhesion (*blue)*. **(C)** Similar to C but using simulation results from Control_cKO_ monolayers (*gray*), *Piezo1* -cKO monolayers without altered adhesion parameters (*white*), and *Piezo1* -cKO monolayers with increased cell-cell adhesion (*orange)* and decreased cell-cell adhesion (*blue)*. In A-C, n = 100 simulation results for each condition. To account for differences between control cases, data are normalized by rescaling to the mean of the corresponding control. Larger normalized wound closure indicates faster wound closure, while a smaller normalized wound closure indicates slower wound closure. Similarly, a larger normalized edge length indicates a more featured wound while a smaller normalized edge length indicates a flatter or less featured wound. Black check marks at the top of each plot condition indicate that simulation results match experimental trends while a red cross indicates the simulations fail to match the experiment results. Related to Table 3. For comparison with experimental data see Fig 1B, 1G and 1H.

**S8 Fig.**
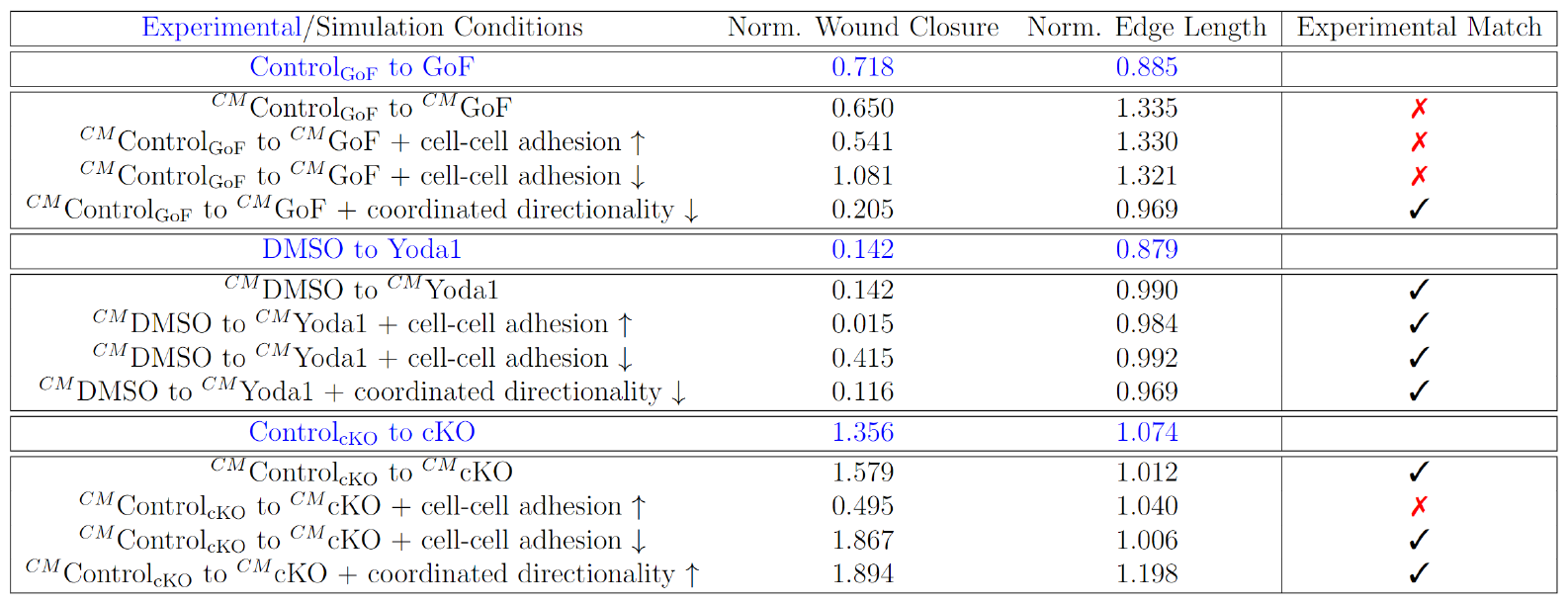
Numerical comparisons between simulations and experiments on wound closure and edge length. The table presents simulation results (in black, see Table 3 for qualitative results) obtained using the calibrated model (^CM^) to predict the impact of PIEZO1 on normalized wound closure and normalized edge length, altering adhesion and coordinated directionality parameters. The simulation results are quantitatively compared with the corresponding experimental results (in blue, see Table 1 for qualitative results). Model predictions are indicated in red font with a cross mark (✗) when they do not align with the experimental trends of increasing or decreasing values. Conversely, a check mark (✓) indicates that model predictions are consistent with the experimental trends.

**S9 Fig.**
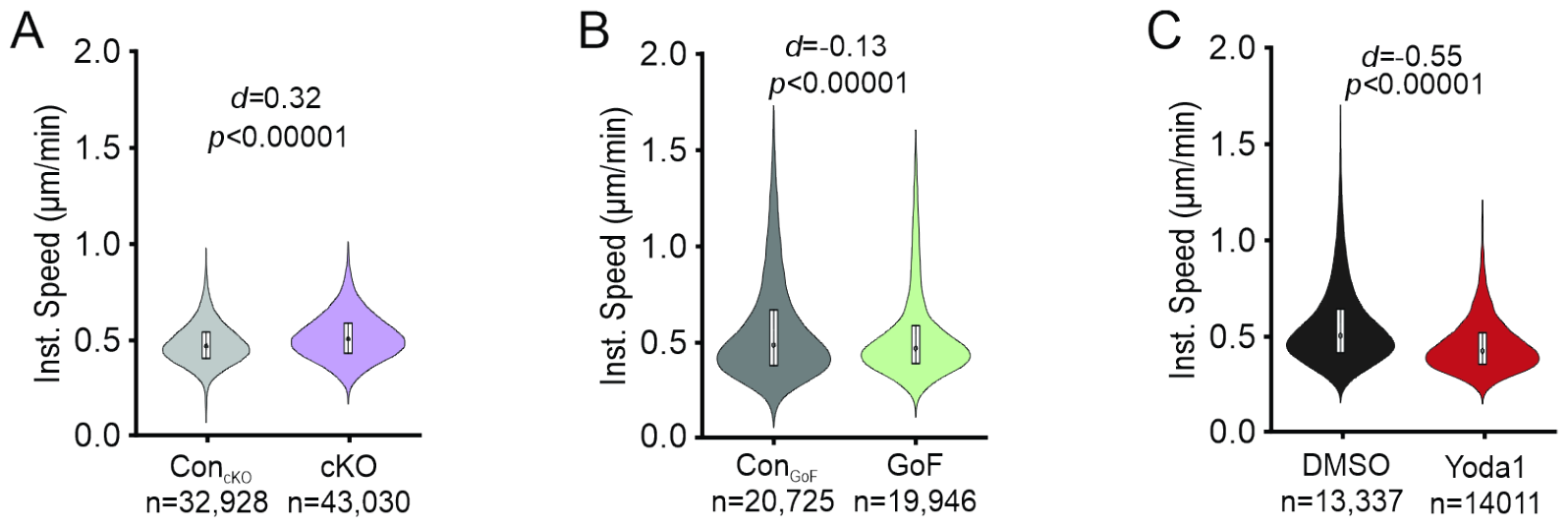
PIEZO1 inhibits keratinocyte speed during collective cell migration. Violin plots quantifying the average instantaneous cell speed of tracked cells in **(A)** Control_cKO_ vs. *Piezo1* -cKO, **(B)** Control_GoF_ vs. *Piezo1* -GoF, and **(C)** DMSO-treated and 4 *μ*M Yoda1-treated keratinocytes monolayers. For A-C, *p* value calculated via Mann-Whitney Test. For A-C, plotted n denotes the number of individual cell trajectories. See also Fig 4.

**S10 Fig.**
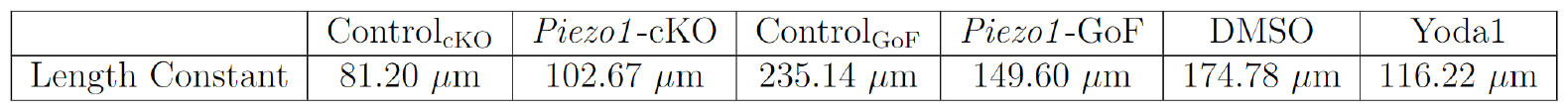
PIEZO1 reduces the length scale of spatial autocorrelation in keratinocytes. Summary table showing the length constant, or the distance at which the spatial autocorrelation value is estimated to reach 0.37, for each experimental condition. Length constants were calculated by fitting a curve to the respective experimental dataset. See also Fig 5.

**S11 Fig.**
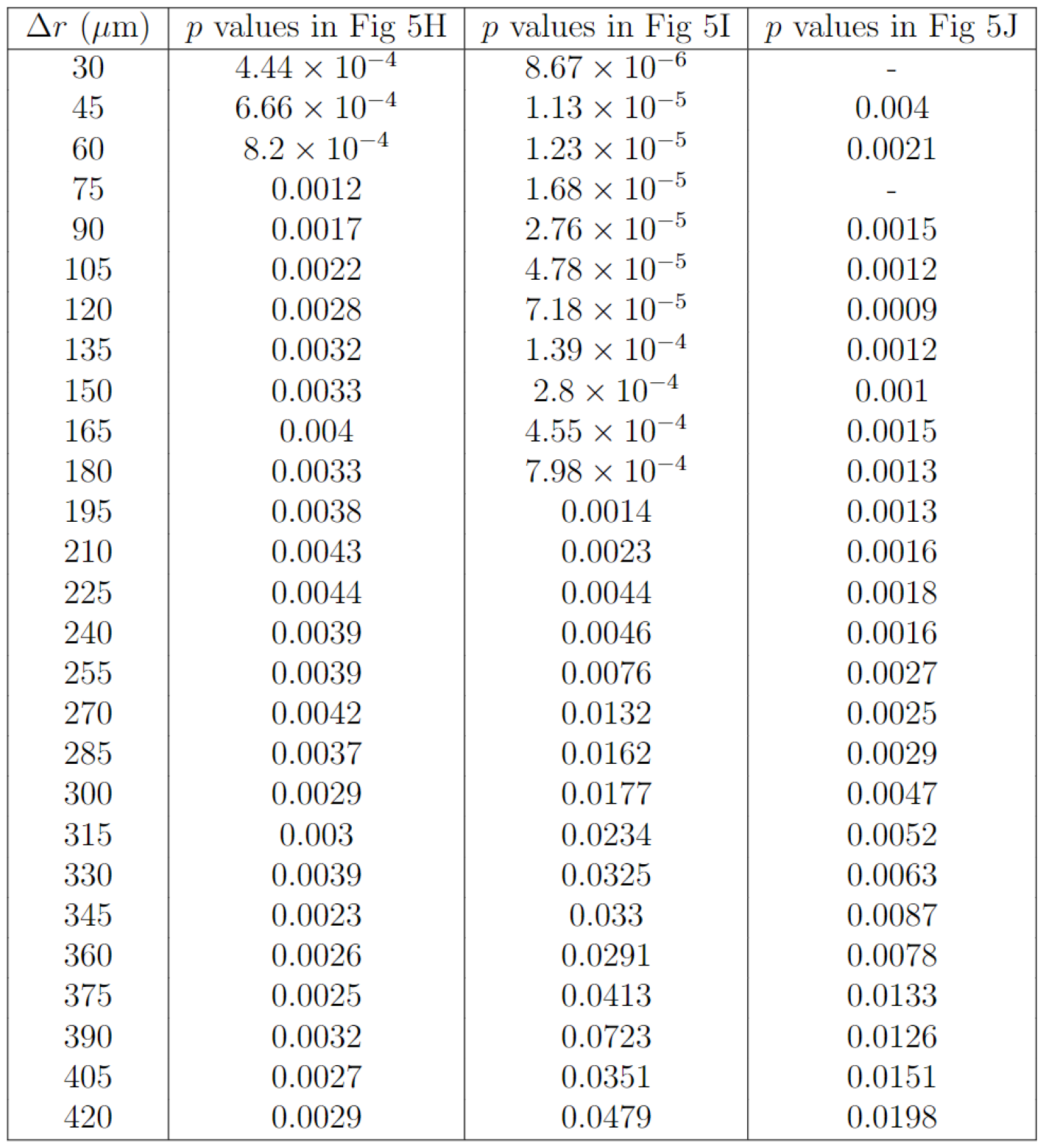
Specific *p* values for plotted points seen in Fig 5 H-J.

**S12 Fig.**
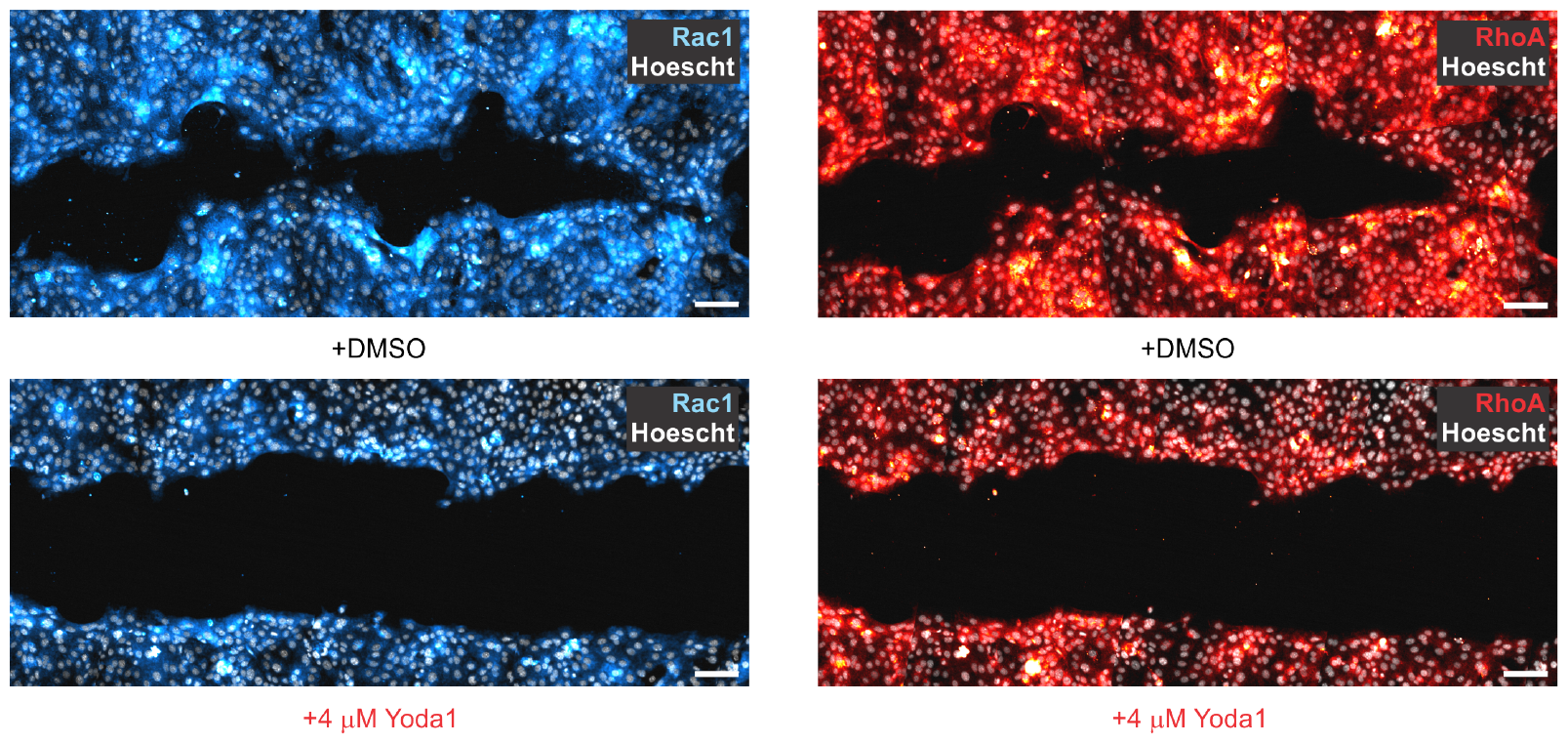
Increased PIEZO1 activity regulates Rho GTPase levels within collectively migrating monolayers. To explore a possible relationship between PIEZO1 and Rho GTPases we performed immunocytochemistry experiments for the Rho GTPases RhoA and Rac1 within healing monolayers. Scratch wounds were generated in keratinocyte monolayers and then immediately treated with either 4 *μ*M Yoda1, or the equivalent amount of solvent DMSO. Keratinocyte monolayers were allowed to collectively migrate for 24 hours with the respective drug in the bath media before fixing and labeling monolayers. Shown above, representative images of healing keratinocyte monolayers immuno-labeled with antibodies against Rac1 (*blue, left panels*), and RhoA (*red, right panels*) 24 hours after wounding and treating monolayers with DMSO (*top*) and 4 *μ*M Yoda1 (*bottom*). Increasing PIEZO1 activity through Yoda1-treatment decreases Rac1 and RhoA staining suggesting that PIEZO1 activity regulates Rho GTPase expression during keratinocyte collective migration. Scale bar=100 *μ*m.

**S13 Fig.**
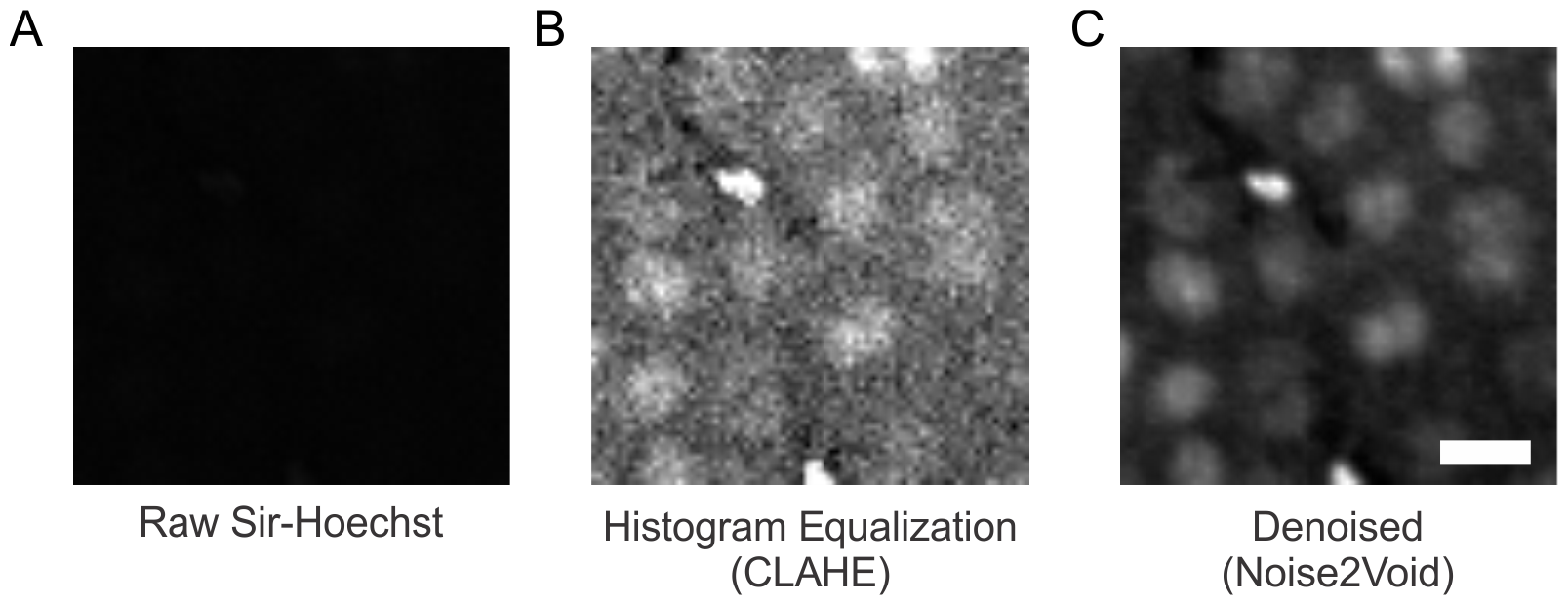
Image processing pipeline for nuclei images. Representative images of processing steps to boost signal-to-noise ratio of **(A)** raw SiR-Hoechst images by first performing **(B)** histogram equalization using Contrast Limited Adaptive Histogram Equalization (CLAHE). **(C)** For some images, the denoising algorithm Noise2Void was used to further increase the signal-to-noise ratio of nuclei. Note: all images adjusted to the same brightness and contrast settings. Scale bar = 20 μm.

**S14 Fig.**
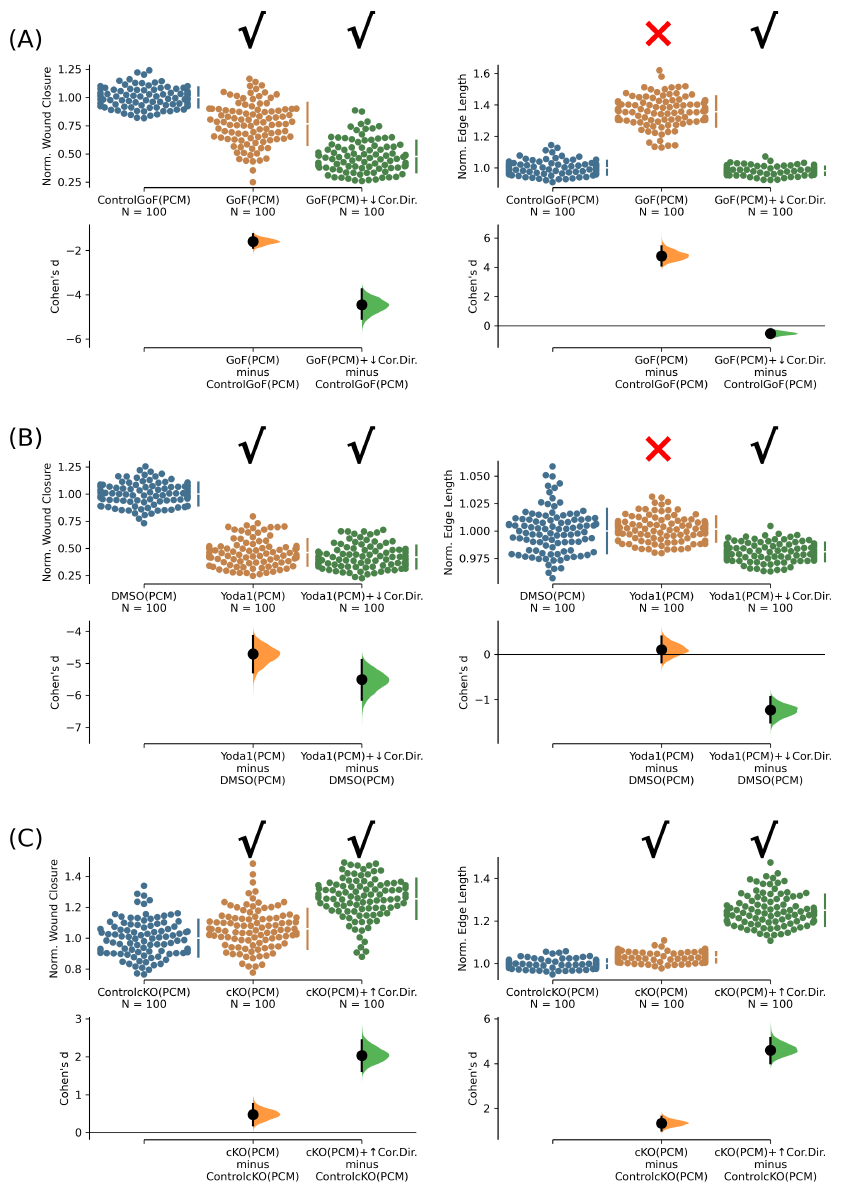
The phenomenological continuum model is consistent with the original prediction that PIEZO1 hinders coordinated directionality in wound closure. **(A)** Cumming plots showing simulation results using the calibrated phenomenological continuum model (denoted as PCM, see Section 7 in S1 Text) to predict how PIEZO1 affects normalized wound closure (*left*) and wound edge length (*right*) in simulated Control_GoF_ monolayers (*blue*), *Piezo1* -GoF monolayers without altered coordinated directionality parameters (*orange*), and *Piezo1* -GoF monolayers with cell coordinated directionality decreased (*green*). See Methods Section for the details on model parameters adjustment. **(B)** Similar to A but using simulation results from DMSO-treated monolayers (*blue*), Yoda1-treated monolayers without altered coordinated directionality parameters (*orange*), and Yoda1-treated monolayers with coordinated directionality decreased (*green*). **(C)** Similar to A but using simulation results from Control_cKO_ monolayers (*blue*), *Piezo1* -cKO monolayers without altered coordinated directionality parameters (*orange*), and *Piezo1* -cKO monolayers with coordinated directionality increased (*green*). In A-C, n = 100 simulation results for each condition, and CM denotes “Calibrated Model”, specifically using the phenomenological continuum model in Section 7 in S1 Text. To account for differences between control cases, data are normalized by rescaling to the mean of the corresponding control. Larger normalized wound closure indicates faster wound closure, while a smaller normalized wound closure indicates slower wound closure. Similarly, a larger normalized edge length indicates a more featured wound edge while a smaller normalized edge length indicates a flatter or less featured wound edge. Black check marks at the top of each plot condition indicate that simulation results match experimental trends while a red cross indicates simulation fails to match the experiment trends. For comparison with experimental data see Fig 1B, 1G and 1H.

**S15 Fig.**
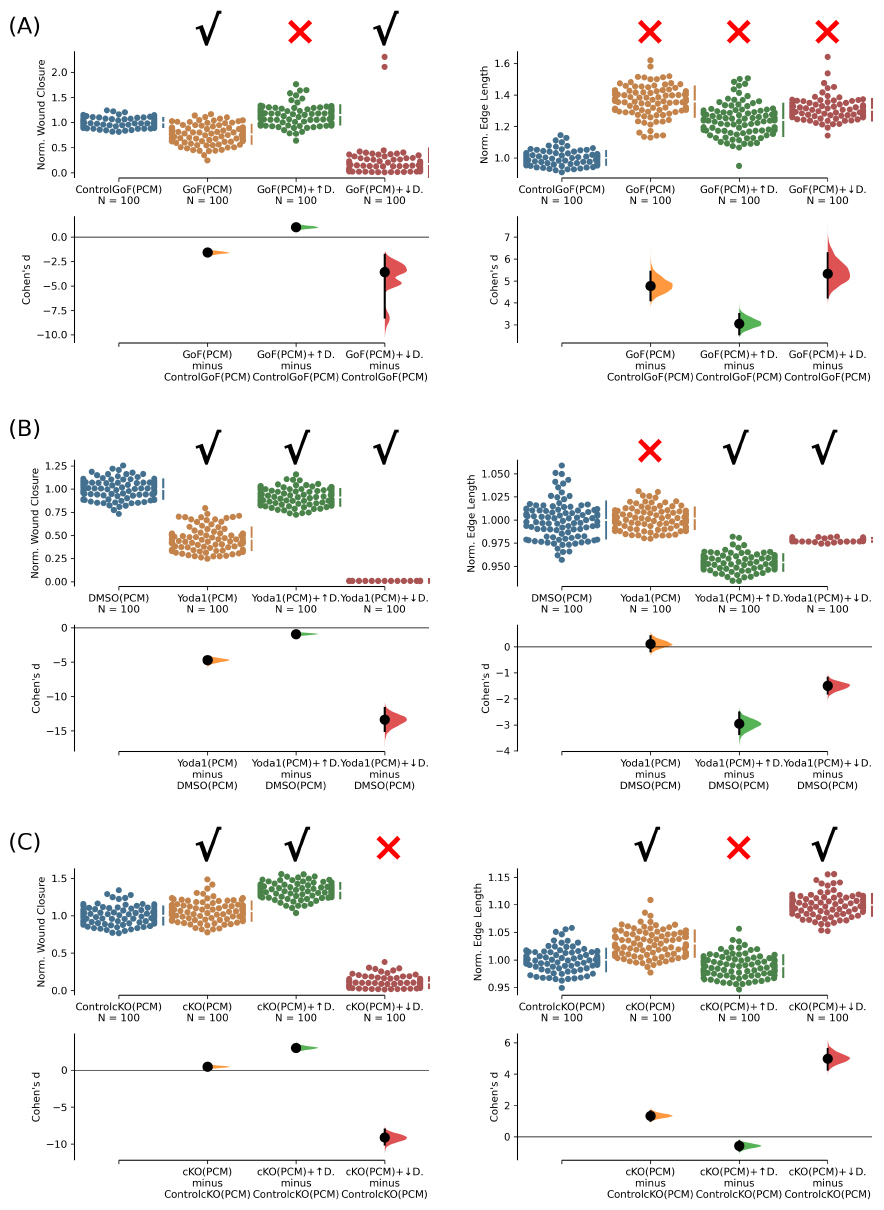
Varying the diffusion coefficient in response to changes in cell-cell adhesion fails to match all the experimental trends. **(A)** Cumming plots showing simulation results in which we use the phenomenological continuum model (denoted as PCM, see Section 7 in S1 Text) to predict how PIEZO1 affects wound closure (*left*) and wound edge length (*right*) in simulated Control_GoF_ monolayers (*blue*), *Piezo1* -GoF monolayers without altered the diffusion coefficient (*orange*), *Piezo1* -GoF monolayers with increased diffusion coefficient (*green*) and decreased diffusion coefficient (*red*). The magnitude of diffusion coefficient models the combined effects of cell motility and cell-cell adhesion. **(B)** Similar to A but using simulation results from DMSO-treated monolayers (*blue*), Yoda1-treated monolayers without altered the diffusion coefficient (*orange*), Yoda1-treated monolayers with increased diffusion coefficient (*green*) and decreased diffusion coefficient (*red*). **(C)** Similar to C but using simulation results from Control_cKO_ monolayers (*blue*), *Piezo1* -cKO monolayers without altered the diffusion coefficient (*orange*), and *Piezo1* -cKO monolayers with increased diffusion coefficient (*green*) and decreased diffusion coefficient (*red*). In A-C, n = 100 simulation results for each condition. To account for differences between control cases, data are normalized by rescaling to the mean of the corresponding control. Larger normalized wound closure indicates faster wound closure, while a smaller normalized wound closure indicates slower wound closure. Similarly, a larger normalized edge length indicates a more featured wound while a smaller normalized edge length indicates a flatter or less featured wound. Black check marks at the top of each plot condition indicate that simulation results match experimental trends while a red cross indicates simulation fails to match the experiment result. Related to S14 Fig. For comparison with experimental data see Fig 1B, 1G and 1H.

**S16 Fig.**
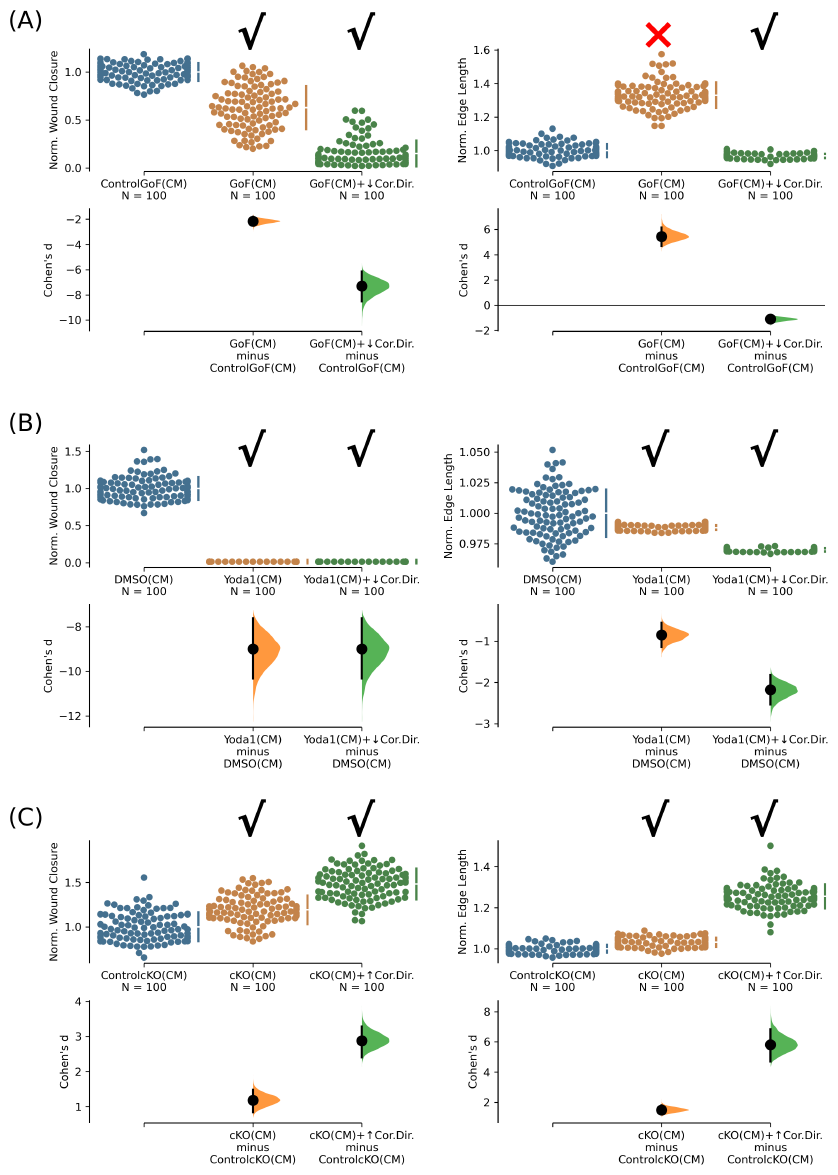
Original model calibrated by monolayer cell motilities is consistent with the original prediction that PIEZO1 hinders coordinated directionality in wound closure. **(A)** Cumming plots showing simulation results from the calibrated model (CM) using cell motilities from monolayer experiments (for data see S9 Fig, for detailed model calibration see Section 8 in S1 Text) to predict how PIEZO1 affects normalized wound closure (*left*) and wound edge length (*right*) in simulated Control_GoF_ monolayers (*blue*), *Piezo1* -GoF monolayers without altered coordinated directionality parameters (*orange*), and *Piezo1* -GoF monolayers with coordinated directionality decreased (*green*). See Methods Section for the details on model parameters adjustment. **(B)** Similar to A but using simulation results from DMSO-treated monolayers (*blue*), Yoda1-treated monolayers without altered coordinated directionality parameters (*orange*), and Yoda1-treated monolayers with coordinated directionality decreased (*green*). **(C)** Similar to A but using simulation results from Control_cKO_ monolayers (*blue*), *Piezo1* -cKO monolayers without altered coordinated directionality parameters (*orange*), and *Piezo1* -cKO monolayers with coordinated directionality increased (*green*). In A-C, n = 100 simulation results for each condition, and CM denotes “Calibrated Model”, specifically our original model using cell motilities from monolayer experiments (Section 8 in S1 Text). To account for differences between control cases, data are normalized by rescaling to the mean of the corresponding control. Larger normalized wound closure indicates faster wound closure, while a smaller normalized wound closure indicates slower wound closure. Similarly, a larger normalized edge length indicates a more featured wound while a smaller normalized edge length indicates a flatter or less featured wound. Black check marks at the top of each plot condition indicate that simulation results match experimental trends while a red cross indicates simulation fails to match the experiment trends. For comparison with experimental data see Fig 1B, 1G and 1H.

**S17 Fig.**
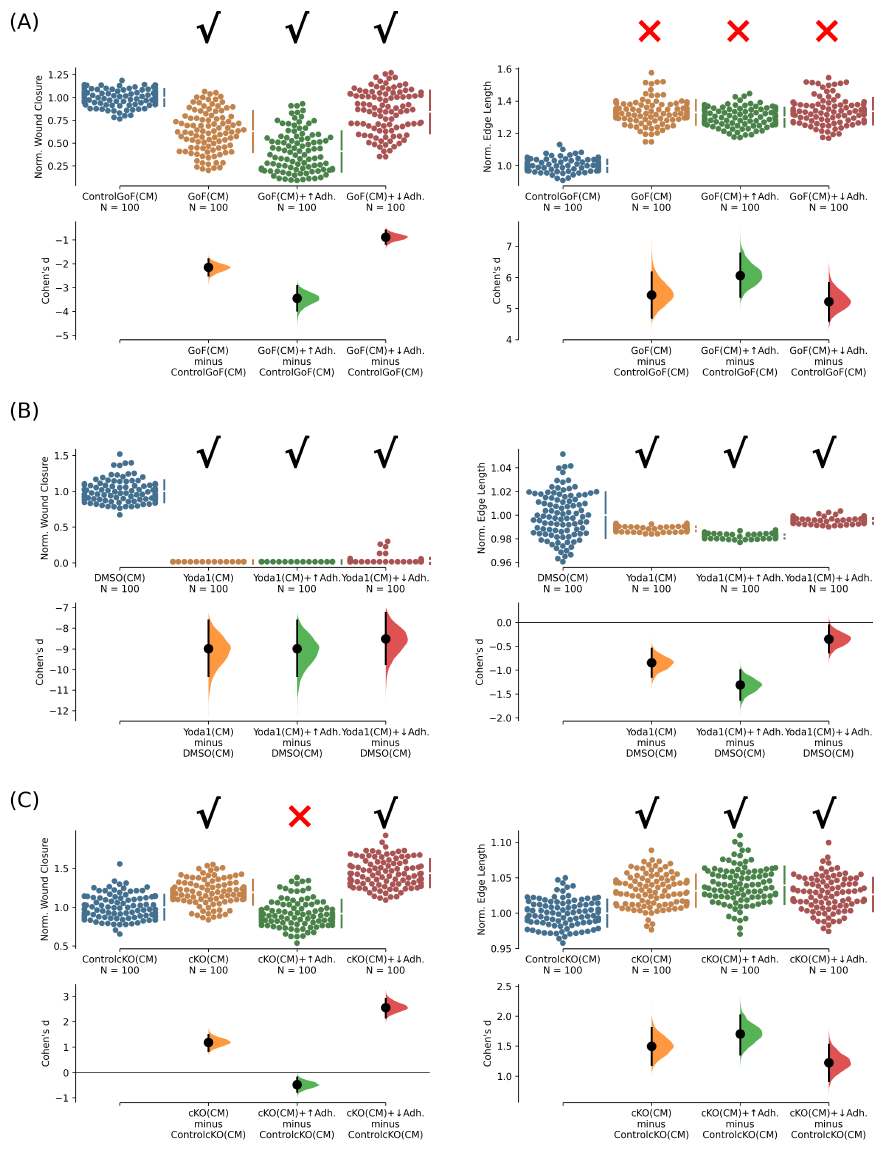
Varying cell-cell adhesion in the original model, calibrated by the monolayer cell motilities, fails to match some experimental trends. **(A)** Cumming plots showing simulation results in which we use our calibrated model using cell motilities from monolayer experiments (denoted as CM, see Section 8 in S1 Text) to predict how PIEZO1 affects wound closure (*left*) and wound edge length (*right*) in simulated Control_GoF_ monolayers (*blue*), *Piezo1* -GoF monolayers without altered the cell-cell adhesion parameter (*orange*), *Piezo1* -GoF monolayers with increased cell-cell adhesion (*green*) and decreased cell-cell adhesion (*red*). **(B)** Similar to A but using simulation results from DMSO-treated monolayers (*blue*), Yoda1-treated monolayers without altered the cell-cell adhesion parameter (*orange*), Yoda1-treated monolayers with increased cell-cell adhesion (*green*) and decreased cell-cell adhesion (*red*). **(C)** Similar to C but using simulation results from Control_cKO_ monolayers (*blue*), *Piezo1* -cKO monolayers without altered the cell-cell adhesion parameter (*orange*), and *Piezo1* -cKO monolayers with increased cell-cell adhesion (*green*) and decreased cell-cell adhesion (*red*). In A-C, n = 100 simulation results for each condition. To account for differences between control cases, data are normalized by rescaling to the mean of the corresponding control. Larger normalized wound closure indicates faster wound closure, while a smaller normalized wound closure indicates slower wound closure. Similarly, a larger normalized edge length indicates a more featured wound while a smaller normalized edge length indicates a flatter or less featured wound. Black check marks at the top of each plot condition indicate that simulation results match experimental trend while a red cross indicates simulation fails to match the experiment result. Related to S16 Fig. For comparison with experimental data see Fig 1B, 1G and 1H.

**S18 Fig.**
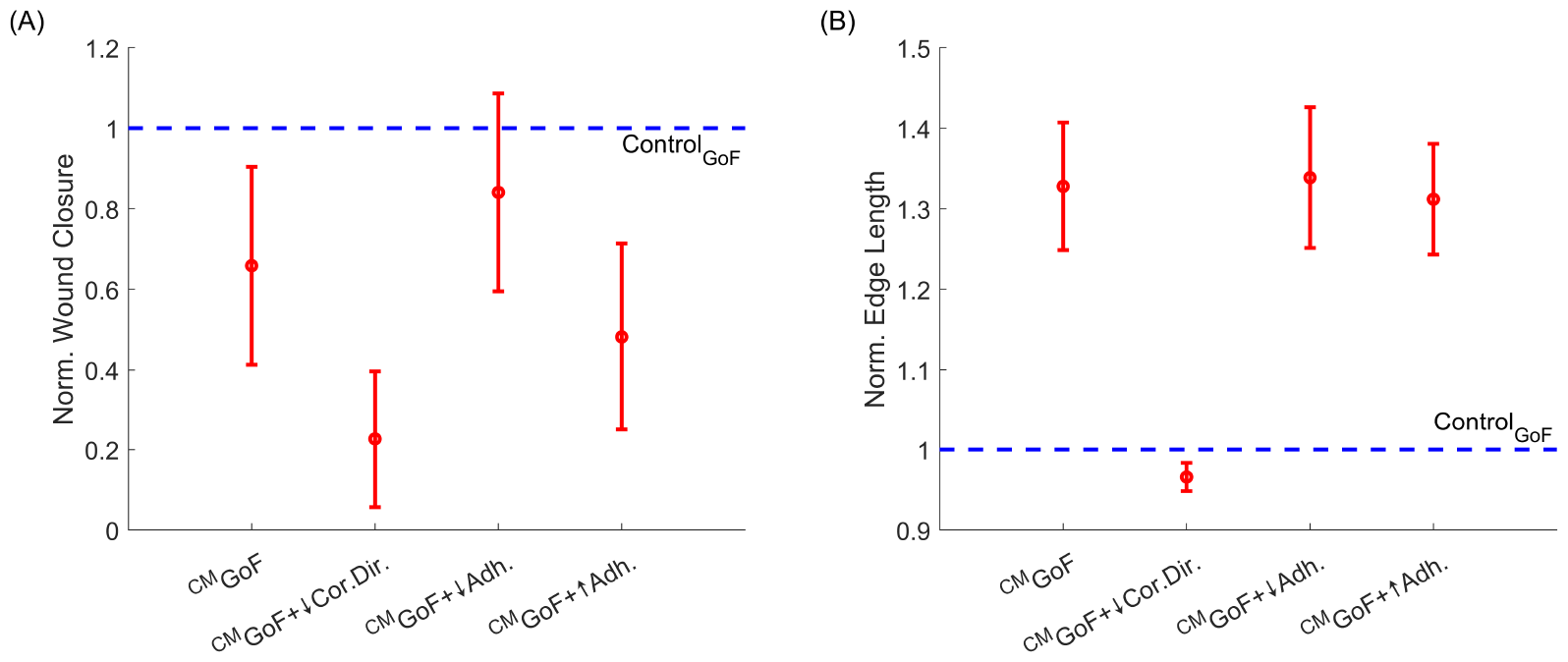
Varying the magnitudes of the retraction processes in the original model (see text) yields results that are consistent with the original prediction that PIEZO1 hinders coordinated directionality in wound closure. **(A)** Dot plots illustrate the mean of 2700 simulation results from the model using three values of the magnitudes of the retraction processes. They depict how PIEZO1 influences normalized wound closure in *Piezo1* -GoF monolayers compared to simulated Control_GoF_ monolayers (*blue dashed line*). The scenarios include *Piezo1* –GoF monolayers without altered coordinated directionality and cell-cell adhesion parameters (*first column*), *Piezo1* -GoF monolayers with decreased coordinated directionality (*second column*), *Piezo1* -GoF monolayers with decreased cell-cell adhesion (*third column*), and *Piezo1* -GoF monolayers with increased cell-cell adhesion (*fourth column*). See Section 8 in S1 Text for the details on the model parameters. Error bars indicate the standard deviation. **(B)** Similar to A but measuring the changes in normalized edge length instead of normalized wound closure.

**S19 Fig.**
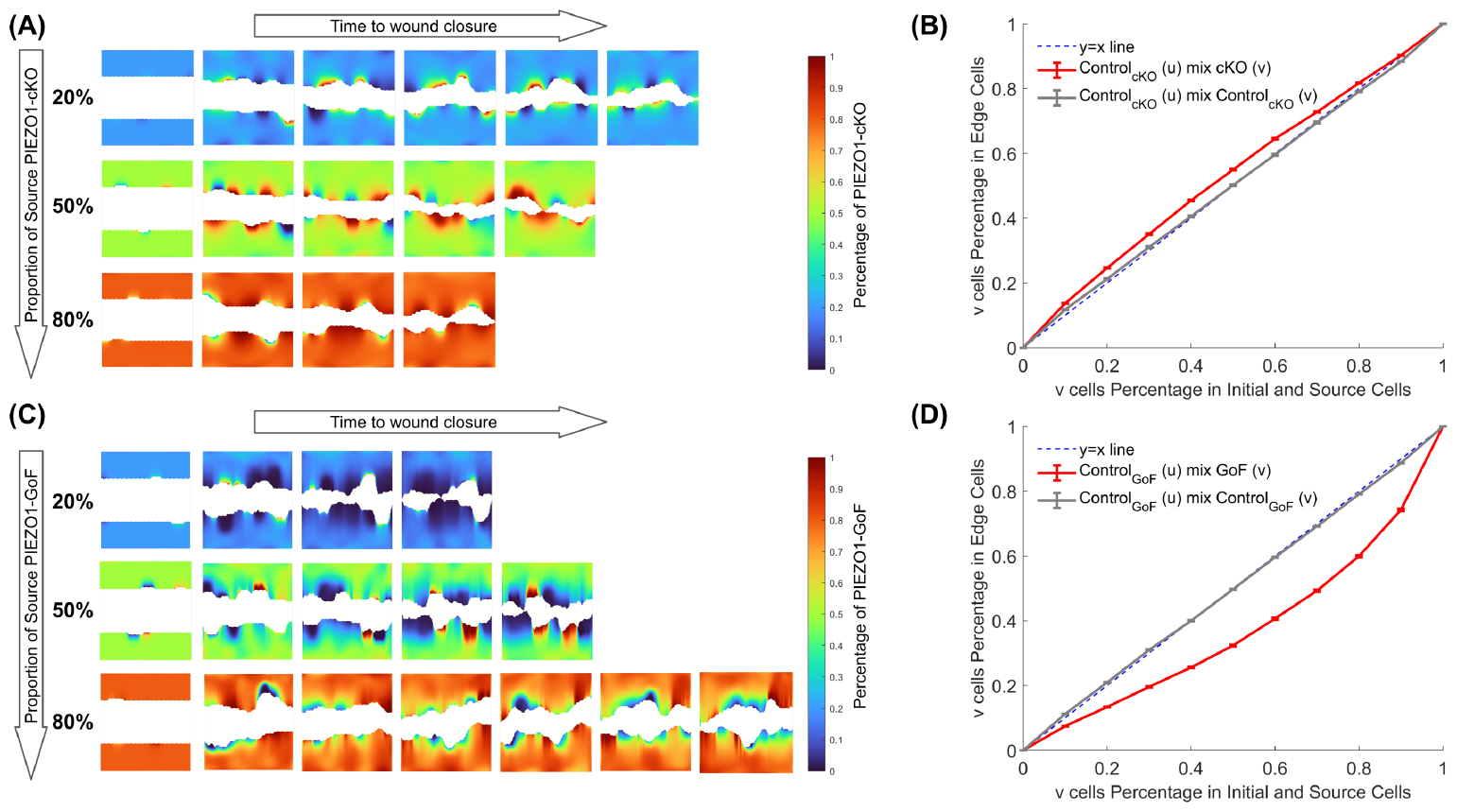
The distribution of edge cells in mixed collective migration correlates with the level of PIEZO1 activity. **(A) and (B)** show that cells with reduced PIEZO1 activity are overrepresented at the leading edge in mixed cell migration. (A) Snapshot of wound healing progression for a mixture of Control_cKO_ (*u* cells) and *Piezo1* -cKO (*v* cells) captured at equidistant time intervals, under varied initial and source cell conditions with *Piezo1* -cKO (*v* cells) percentages of 20% (*top*), 50% (*middle*), and 80% (*bottom*). Colored areas represent cell monolayers, with colors indicating the spatial distribution of *Piezo1* -cKO (*v* cells) percentage, while plain white areas denote cell-free space. (B) Line graphs illustrate the mean of 100 simulation results, displaying the percentage of *Piezo1* -cKO (*v* cells) cells in edge cells versus the percentage in initial and source cells. The red solid curve represents the mixing of *Piezo1* -cKO (*v* cells) with Control_cKO_ (*u* cells), the gray solid curve represents the scenario of mixing the same wild-type Control_cKO_ cells (i.e., u and *v* cells are both Control_cKO_), and the blue dashed line signifies the *y* = *x* line. Error bars indicate the standard error of the mean. **(C) and (D)** Similar to (A) and (B), but involving mixtures of *Piezo1* -GoF (*v* cells) with their respective wild-type control (*u* cells). *Piezo1* -GoF (*v* cells) are underrepresented at the leading edge during migration.

